# Structural Optimization of CHI3L1 Inhibitors with Improved Pharmacokinetics and Functional Activity in 3D Glioblastoma Models

**DOI:** 10.64898/2026.01.28.702243

**Authors:** Baljit Kaur, Hossam Nada, Moustafa T. Gabr

**Affiliations:** Department of Radiology, Molecular Imaging Innovations Institute (MI3), Weill Cornell Medicine, New York, NY 10065, USA

**Author notes:** To whom correspondence should be addressed: Moustafa T. Gabr.

**Keywords:** GBM, CHI3L1, GBM spheroid, drug design, PK profiling

## Abstract

Chitinase-3-like protein 1 (CHI3L1) is a key driver of glioblastoma (GBM) progression and an emerging therapeutic target. Building on the CHI3L1 inhibitor **11g**, we optimized the scaffold through medicinal chemistry to assess structure-property relationships and improve pharmacokinetics. Using microscale thermophoresis (MST) and computational studies, we validated **10p**, which exhibits a CHI31 binding affinity (*K*_d_) of 13.22 µM. Notably, **10p** overcomes previous developability hurdles by achieving a kinetic solubility of 758 µM, a five-fold improvement over **11g**. It further demonstrates high metabolic stability across species and no hERG inhibition. In 3D GBM spheroid models, **10p** significantly reduced tumor viability, mass, and migration, exceeding the efficacy of prior analogues. Collectively, these findings establish **10p** as a potent CHI3L1 inhibitor with a superior pharmacokinetic profile and robust functional activity, marking it as a promising candidate for further GBM drug development.

**Graphical Abstract:** 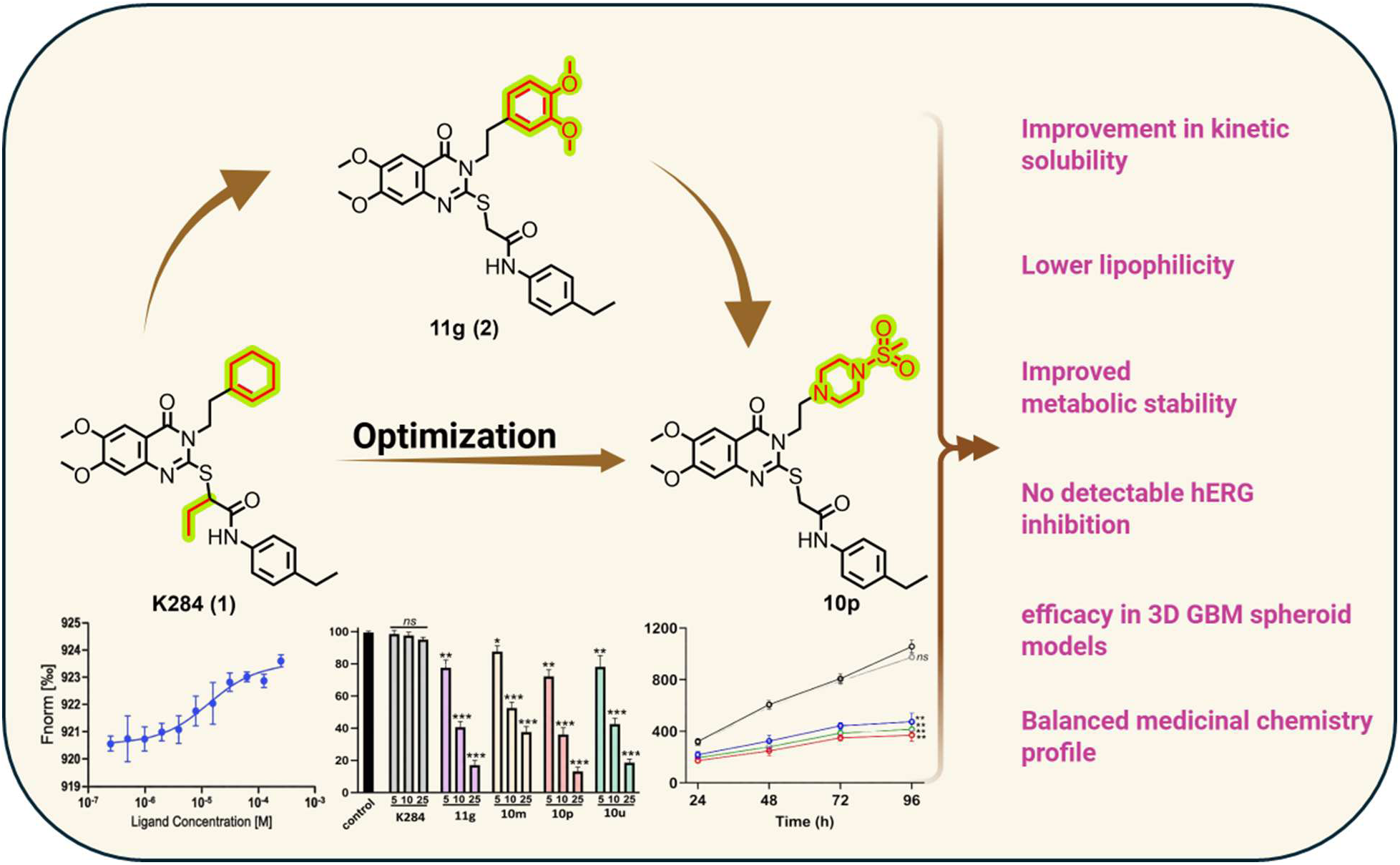

## Introduction

Glioblastoma (GBM) is an aggressive malignancy with limited therapeutic options, partly due to the difficulty of achieving effective modulation of disease-relevant molecular targets within the central nervous system.^1–6^ Traditional interventions offer only temporary reprieve, as tumors almost invariably recur in a more aggressive form.^7,8^ This resilience stems from the tumor’s reliance on complex molecular networks that orchestrate proliferation, invasion, angiogenesis, and immune evasion, rendering single-pathway therapies largely ineffective.^9–12^ A paradigm shift is therefore needed: instead of targeting downstream effects, therapies must disrupt the central regulators that enable GBM’s survival and adaptability.^13,14^ Among these central regulators, Chitinase-3-like protein 1 (CHI3L1, YKL-40) has emerged as a critical driver of GBM malignancy.^15–18^ CHI3L1 is abundantly expressed not only in tumor cells but also in the surrounding stromal and immune compartments,^19–29^ where it coordinates invasion, vascular remodeling, and immune suppression.^30^ High CHI3L1 levels strongly correlate with aggressive tumor phenotypes and poor patient survival, suggesting that GBM leverages this protein to maintain its malignant edge.^18^ Unlike conventional targets that influence single pathways, CHI3L1 functions as a molecular hub, making it an attractive candidate for therapeutic intervention. The small-molecule inhibitor **K284** (**1**)was previously reported to interfere with CHI3L1-mediated signaling, providing an initial chemical framework for CHI3L1 modulation.^31–34^ In our earlier work, medicinal chemistry optimization of this scaffold led to the identification of compound **11g (2)** (Figure 1),^35^ which exhibited improved CHI3L1 engagement and functional activity in three-dimensional GBM spheroid models. This study established the quinazolinone-based scaffold as a viable platform for CHI3L1 inhibition and enabled preliminary structure–activity relationship (SAR) analysis.

**Figure 1.**
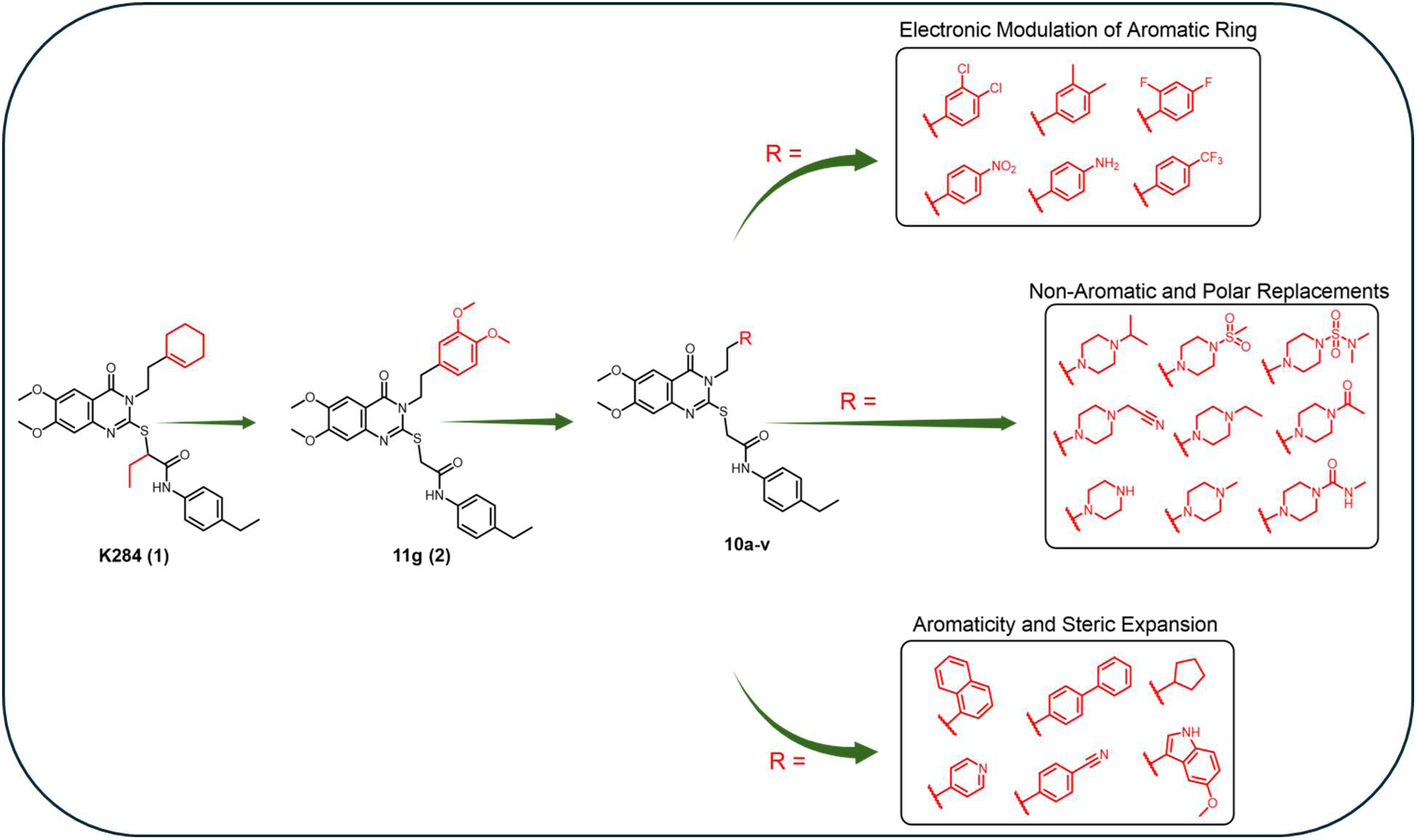
Evolution of CHI3L1 inhibitor design and focused substituent optimization. The previously reported CHI3L1 inhibitor **K284** (**1**) served as the initial scaffold for medicinal chemistry optimization, leading to the identification of compound **11g** (**2**) with improved CHI3L1 engagement. In the present study, further optimization in compounds **10a-v** was focused on systematic modification of the 3,4-dimethoxyphenethyl substituent of **11g**. Variations were designed to probe electronic effects, aromatic surface and topology, and the influence of polarity and aromaticity on CHI3L1 binding and physicochemical properties.

Despite improved biological activity, compound **11g** displayed limitations in aqueous solubility, metabolic stability, and overall pharmacokinetic behavior. These findings indicated that further optimization was required to better understand the structural determinants governing drug-like properties within this series. Analysis of SAR trends from the initial study suggested that the substituent corresponding to the 3,4-dimethoxyphenethyl moiety played a critical role in CHI3L1 interaction while occupying a region of the binding interface that appeared tolerant to structural modification. Based on this observation, the present study focuses on systematic variation of this substituent to more comprehensively define its contribution to CHI3L1 binding and modulation of physicochemical properties. A diverse set of replacements was selected to probe key molecular features, including electronic effects, steric demand, aromatic surface area, and overall polarity (Figure 1). Substituted phenyl rings bearing electron-donating and electron-withdrawing groups were examined to assess how changes in electronic distribution influence CHI3L1 engagement while maintaining a comparable aromatic framework. To further explore the role of aromaticity and ring composition, heteroaromatic systems were incorporated to introduce localized polarity and potential hydrogen-bonding interactions, as well as to modulate π-stacking and dipole characteristics. In parallel, extended aromatic motifs were evaluated to probe the tolerance of this region to increased aromatic surface area and conformational rigidity. Non-aromatic replacements were also included to determine whether aromatic character at this position is essential for CHI3L1 interaction or whether alternative hydrophobic or polar functionalities could be accommodated. In addition, polar and basic substituents were incorporated to systematically increase hydrophilicity and ionizability, enabling assessment of their impact on solubility and in vitro metabolic behavior, while preserving the core structural features required for CHI3L1 engagement.

Rather than prioritizing maximal gains in potency, this work emphasizes evaluation of structure–property relationships within the **K284**-derived scaffold. By examining how targeted modifications influence physicochemical and in vitro ADME parameters while maintaining biological activity, this study aims to define the practical optimization boundaries of this chemical series.

Herein, we report the biological and physicochemical evaluation of a focused set of CHI3L1 inhibitors. The results provide insight into scaffold-dependent constraints and inform future medicinal chemistry strategies for the development of CHI3L1-directed small-molecule inhibitors. amenable to further modification. This observation provided the rationale for a second-stage optimization effort focused exclusively on this substituent, with the objective of probing the tolerance of this region to electronic, steric, and topological variation.

## Materials and methods

### Chemistry

#### General procedures for chemistry

All reagents, solvents, and materials were obtained from commercial suppliers and used without further purification. ^1^H NMR spectra were recorded on a Bruker 500 MHz spectrometer, with chemical shifts (δ) reported in ppm relative to an internal standard. Signal multiplicities are denoted as s, d, dd, t, q, td, dt, ddd, br, and m. Flash chromatography was performed using a CombiFlash system with RediSep RF silica cartridges, and reaction progress was monitored by TLC on silica gel 60 F₂₅₄ plates.

#### Synthetic procedures

##### General procedure A: for the synthesis of intermediates 6a–l

6,7-Dimethoxy-2H-benzo[d][1,3]oxazine-2,4(1H)-dione (**3**, 1.0 mmol) was dissolved in DMF, and the corresponding amines (**4a–l**, 1.5 mmol) were added. The mixture was stirred at room temperature overnight until completion (TLC). The solvent was removed under reduced pressure, and the crude products **5a–l** were used directly in the next step. Intermediates **5a–l** (1.0 mmol) were dissolved in ethanol, treated with aqueous NaOH (2.2 mmol), followed by carbon disulfide (4.0 mmol). The reaction was heated at 60 °C overnight, then cooled and acidified with 1 M HCl to precipitate the product. The solid was filtered and washed with acetone to obtain intermediates **6a–l**.

##### General procedure B: for the synthesis of intermediates 9

4-Ethylaniline (**7**, 1.0 mmol) in dry DCM was reacted with 2-bromoacetyl bromide (**8**, 1.2 mmol) at 0 °C for 30 min. After completion (TLC), the reaction mixture was extracted with DCM, washed with water and brine, dried, and concentrated. Purification by flash chromatography (hexane:ethyl acetate, 10:1) afforded intermediate **9** in 84% yield.

##### General procedure C: for the synthesis of compounds 10a–l

Intermediates **6a–k** (1.0 mmol) were reacted with amine **9** (1.0 mmol) in acetone in the presence of K₂CO₃ (1.5 mmol) at 60 °C overnight. After solvent removal, the residue was extracted with DCM, washed, dried, and concentrated. Final purification by flash chromatography (hexane:ethyl acetate, 3:2 to 1:1) yielded compounds **10a–k** in 50–80% yield.

###### 2-((3-(3,4-dimethylphenethyl)-6,7-dimethoxy-4-oxo-3,4-dihydroquinazolin-2-yl)thio)-N-(4-ethylphenyl)acetamide (10a)

White solid; yield 76%; ^1^H NMR (500 MHz, CDCl_3_) δ 9.78 (s, 1H), 7.61 (s, 1H), 7.39 – 7.32 (m, 2H), 7.12 – 7.02 (m, 6H), 4.35 – 4.21 (m, 2H), 4.02 (d, J = 2.9 Hz, 8H), 3.12 – 2.95 (m, 2H), 2.58 (q, J = 7.6 Hz, 2H), 2.23 (d, J = 15.9 Hz, 6H), 1.18 (t, J = 7.6 Hz, 3H); ^13^C NMR (126 MHz, CDCl_3_) δ 166.60, 160.50, 155.57, 155.49, 148.93, 142.93, 140.42, 137.00, 135.73, 135.22, 134.81, 130.25, 129.97, 128.41, 126.25, 119.43, 112.79, 106.32, 105.73, 56.51, 56.39, 46.87, 35.89, 33.74, 28.30, 19.70, 19.35, 15.69; HRMS: calcd for C_30_H_33_N_3_O_4_S [M+H] m/z 532.2270, found 532.2269.

###### 2-((3-(3,4-dichlorophenethyl)-6,7-dimethoxy-4-oxo-3,4-dihydroquinazolin-2-yl)thio)-N-(4-ethylphenyl)acetamide (10b)

White solid; yield 55%; ^1^H NMR (500 MHz, CDCl_3_) δ 9.58 (s, 1H), 8.58 (d, *J* = 5.6 Hz, 2H), 7.56 (s, 1H), 7.39 – 7.35 (m, 3H), 7.11 (d, *J* = 8.2 Hz, 2H), 7.02 (s, 1H), 4.39 (dd, *J* = 9.5, 6.5 Hz, 2H), 4.03 (d, *J* = 9.8 Hz, 8H), 3.16 (t, *J* = 8.0 Hz, 2H), 2.58 (t, *J* = 7.6 Hz, 2H), 1.18 (t, *J* = 7.6 Hz, 3H); ^13^C NMR (126 MHz, CDCl_3_) δ 166.17, 160.45, 155.81, 154.67, 149.15, 147.91, 142.88, 140.61, 135.56, 128.47, 124.99, 119.43, 112.50, 106.24, 105.88, 56.54, 56.42, 45.01, 35.91, 33.74, 28.29, 15.67; HRMS: calcd for C_28_H_27_Cl_2_N_3_O_4_S [M+H] m/z 572.1178, found 572.1194.

###### 2-((3-(2,4-difluorophenethyl)-6,7-dimethoxy-4-oxo-3,4-dihydroquinazolin-2-yl)thio)-N-(4-ethylphenyl)acetamide (10c)

White solid; yield 80%; ^1^H NMR (500 MHz, CDCl_3_) δ 9.66 (s, 1H), 7.58 (s, 1H), 7.34 (d, J = 8.3 Hz, 2H), 7.18 (d, J = 6.3 Hz, 1H), 7.10 (d, J = 8.2 Hz, 2H), 7.01 (s, 1H), 6.79 – 6.72 (m, 2H), 4.37 – 4.30 (m, 2H), 4.01 (s, 6H), 3.98 (s, 2H), 3.10 (t, J = 7.7 Hz, 2H), 2.58 (q, J = 7.6 Hz, 2H), 1.17 (dd, J = 7.6, 0.8 Hz, 3H); ^13^C NMR (126 MHz, CDCl_3_) δ 166.44, 160.51, 155.65, 155.36, 148.98, 142.88, 140.45, 135.68, 131.87, 131.82, 131.79, 131.74, 128.42, 119.43, 112.65, 111.54, 111.51, 111.37, 111.34, 106.31, 105.79, 104.18, 103.98, 103.78, 56.52, 56.40, 45.02, 35.85, 28.30, 27.24, 15.68; HRMS: calcd for C_28_H_27_F_2_N_3_O_4_S [M+H] m/z 540.1769, found 540.1767.

###### 2-((6,7-dimethoxy-3-(4-nitrophenethyl)-4-oxo-3,4-dihydroquinazolin-2-yl)thio)-N-(4-ethylphenyl)acetamide (10d)

White solid; yield 62%; ^1^H NMR (500 MHz, CDCl_3_) δ 9.55 (s, 1H), 8.14 – 8.07 (m, 2H), 7.50 (s, 1H), 7.45 – 7.37 (m, 2H), 7.31 – 7.25 (m, 2H), 7.08 – 7.01 (m, 2H), 6.96 (s, 1H), 4.35 – 4.25 (m, 2H), 3.95 (d, *J* = 7.8 Hz, 8H), 3.16 – 3.08 (m, 2H), 2.51 (q, *J* = 7.6 Hz, 2H), 1.11 (t, *J* = 7.6 Hz, 3H); ^13^C NMR (126 MHz, CDCl_3_) δ 166.21, 160.41, 155.78, 154.91, 149.13, 147.15, 144.98, 142.80, 140.58, 135.57, 129.86, 128.46, 124.02, 119.38, 112.54, 106.25, 105.79, 56.54, 56.42, 45.69, 35.87, 33.99, 28.29, 15.66; HRMS: calcd for C_28_H_28_N_4_O_6_S [M+H] m/z 549.1808, found 549.1814.

###### 2-((6,7-dimethoxy-4-oxo-3-(4-(trifluoromethyl)phenethyl)-3,4-dihydroquinazolin-2-yl)thio)-N-(4-ethylphenyl)acetamide (10e)

White solid; yield 70%; ^1^H NMR (500 MHz, CDCl_3_) δ 9.70 (s, 1H), 7.59 (d, J = 7.4 Hz, 3H), 7.44 (d, J = 7.9 Hz, 2H), 7.35 (d, J = 8.3 Hz, 2H), 7.11 (d, J = 8.2 Hz, 2H), 7.03 (s, 1H), 4.40 – 4.28 (m, 2H), 4.02 (d, J = 2.2 Hz, 8H), 3.19 – 3.09 (m, 2H), 2.58 (q, J = 7.6 Hz, 2H), 1.58 (s, 2H), 1.18 (t, J = 7.6 Hz, 3H); ^13^C NMR (126 MHz, CDCl_3_) δ 166.37, 160.47, 155.71, 155.09, 149.07, 142.91, 141.52, 140.52, 135.64, 129.31, 128.44, 125.78, 125.75, 125.72, 125.69, 119.41, 112.66, 106.28, 105.80, 56.53, 56.41, 46.10, 35.89, 34.01, 28.30, 15.68; HRMS: calcd for C_29_H_28_F_3_N_3_O_4_S [M+H] m/z 572.1831, found 572.1889.

###### 2-((3-(4-cyanophenethyl)-6,7-dimethoxy-4-oxo-3,4-dihydroquinazolin-2-yl)thio)-N-(4-ethylphenyl)acetamide (10f)

White solid; yield 79%; ^1^H NMR (500 MHz, CDCl_3_) δ 9.63 (s, 1H), 7.66 – 7.55 (m, 3H), 7.45 – 7.38 (m, 2H), 7.38 – 7.32 (m, 2H), 7.15 – 7.08 (m, 2H), 7.02 (s, 1H), 4.39 – 4.27 (m, 2H), 4.02 (s, 8H), 3.22 – 2.99 (m, 2H), 2.59 (d, *J* = 7.6 Hz, 2H), 1.18 (t, *J* = 7.6 Hz, 3H); ^13^C NMR (126 MHz, CDCl_3_) δ 166.27, 160.45, 155.76, 154.92, 149.11, 142.93, 142.88, 140.57, 135.60, 132.59, 129.78, 128.48, 119.39, 118.70, 112.58, 111.08, 106.26, 105.81, 56.54, 56.42, 45.74, 35.85, 34.26, 28.29, 15.67; HRMS: calcd for C_29_H_28_N_4_O_4_S [M+H] m/z 529.1910, found 529.1889.

###### 2-((6,7-dimethoxy-3-(2-(naphthalen-1-yl)ethyl)-4-oxo-3,4-dihydroquinazolin-2-yl)thio)-N-(4-ethylphenyl)acetamide (10g)

White solid; yield 65%;^1^H NMR (500 MHz, CDCl_3_) δ 9.70 (s, 1H), 8.33 (d, J = 8.5 Hz, 1H), 7.79 (d, J = 8.1 Hz, 1H), 7.70 (d, J = 8.0 Hz, 1H), 7.60 (s, 1H), 7.57 – 7.52 (m, 1H), 7.46 – 7.32 (m, 3H), 7.29 (d, J = 8.3 Hz, 2H), 7.04 (d, J = 8.2 Hz, 2H), 6.98 (d, J = 1.6 Hz, 1H), 4.43 – 4.31 (m, 2H), 3.96 (d, J = 8.5 Hz, 8H), 3.54 – 3.43 (m, 2H), 2.52 (q, J = 7.6 Hz, 2H), 1.11 (t, J = 7.6 Hz, 3H); ^13^C NMR (126 MHz, CDCl_3_) δ 166.46, 160.71, 155.67, 155.57, 155.55, 149.06, 142.85, 140.47, 135.69, 133.90, 133.73, 132.14, 128.80, 128.41, 127.85, 127.30, 126.62, 125.90, 125.64, 123.72, 119.49, 112.76, 106.28, 105.70, 56.55, 56.45, 46.48, 35.97, 31.58, 28.30, 15.69; HRMS: calcd for C_32_H_31_N_3_O_4_S [M+H] m/z 554.2114, found 554.2120.

###### 2-((3-(2-([1,1’-biphenyl]-4-yl)ethyl)-6,7-dimethoxy-4-oxo-3,4-dihydroquinazolin-2-yl)thio)-N-(4-ethylphenyl)acetamide (10h)

White solid; yield 70%; ^1^H NMR (500 MHz, CDCl_3_) δ 9.78 (s, 1H), 7.61 (s, 1H), 7.55 (dd, J = 7.6, 3.6 Hz, 4H), 7.44 – 7.32 (m, 7H), 7.13 – 7.01 (m, 3H), 4.38 (t, J = 8.1 Hz, 2H), 4.02 (d, J = 2.5 Hz, 8H), 3.12 (dd, J = 9.4, 6.8 Hz, 2H), 2.58 (d, J = 7.6 Hz, 2H), 1.18 (t, J = 7.5 Hz, 3H); ^13^C NMR (126 MHz, CDCl_3_) δ 166.42, 160.40, 155.65, 149.03, 140.77, 140.44, 140.02, 136.47, 135.65, 129.40, 128.77, 128.40, 127.51, 127.26, 127.07, 119.47, 112.69, 106.32, 105.64, 56.55, 56.41, 46.71, 33.83, 28.29, 15.67; HRMS: calcd for C_34_H_33_N_3_O_4_S [M+H] m/z 580.2270, found 580.2293.

###### 2-((6,7-dimethoxy-3-(2-(5-methoxy-1H-indol-3-yl)ethyl)-4-oxo-3,4-dihydroquinazolin-2-yl)thio)-N-(4-ethylphenyl)acetamide (10i)

White solid; yield 72%; ^13^C NMR (126 MHz, DMSO) δ 165.04, 159.30, 154.17, 153.38, 152.54, 147.46, 142.45, 138.26, 136.17, 130.81, 127.39, 126.81, 123.15, 118.59, 111.60, 111.18, 110.70, 109.30, 106.10, 104.93, 99.42, 55.20, 55.16, 54.67, 44.35, 36.24, 26.98, 23.29, 15.10; HRMS: calcd for C_31_H_32_N_4_O_5_S [M+H] m/z 573.2172, found 573.2185.

###### 2-((6,7-dimethoxy-4-oxo-3-(2-(pyridin-4-yl)ethyl)-3,4-dihydroquinazolin-2-yl)thio)-N-(4-ethylphenyl)acetamide (10j)

White solid; yield 79%; ^1^H NMR (500 MHz, CDCl_3_) δ 9.68 (s, 1H), 7.59 (s, 1H), 7.43 – 7.32 (m, 4H), 7.26 (s, 1H), 7.18 – 7.09 (m, 3H), 7.02 (s, 1H), 4.34 – 4.26 (m, 2H), 4.05 – 3.97 (m, 8H), 3.08 – 2.97 (m, 2H), 2.58 (d, *J* = 7.6 Hz, 2H), 1.18 (t, *J* = 7.6 Hz, 3H); ^13^C NMR (126 MHz, CDCl_3_) δ 166.35, 160.44, 155.72, 155.00, 149.07, 142.90, 140.52, 137.60, 135.63, 132.76, 130.91, 130.72, 128.44, 128.34, 119.43, 112.64, 106.28, 105.81, 56.53, 56.42, 46.02, 35.88, 33.30, 28.30, 15.68; HRMS: calcd for C_27_H_28_N_4_O_4_S [M+H] m/z 572.1729, found 572.1719.

###### 2-((3-(2-cyclopentylethyl)-6,7-dimethoxy-4-oxo-3,4-dihydroquinazolin-2-yl)thio)-N-(4-ethylphenyl)acetamide (10k)

thick oil; yield 70%; ^1^H NMR (500 MHz, CDCl_3_) δ 9.83 (s, 1H), 7.58 (s, 1H), 7.38 – 7.31 (m, 2H), 7.13 – 7.07 (m, 2H), 7.00 (s, 1H), 4.20 – 4.10 (m, 2H), 4.00 (d, *J* = 2.9 Hz, 8H), 2.58 (q, *J* = 7.6 Hz, 2H), 1.95 – 1.74 (m, 5H), 1.69 – 1.61 (m, 2H), 1.55 (dd, *J* = 7.6, 4.7 Hz, 2H), 1.28 – 1.20 (m, 2H), 1.18 (t, *J* = 7.6 Hz, 3H); ^13^C NMR (126 MHz, CDCl_3_) δ 166.68, 160.52, 155.57, 155.47, 148.84, 142.90, 140.37, 135.75, 128.40, 119.39, 112.79, 106.33, 105.65, 56.48, 56.36, 44.71, 37.78, 35.87, 34.24, 32.44, 28.29, 25.19, 15.69; HRMS: calcd for C_27_H_33_N_3_O_4_S [M+H] m/z 496.2270, found 496.2291.

##### General procedure D: for the synthesis of compound 10m

To a solution of Intermediate **10l** (1 mmol) dissolved in CH_2_Cl_2_ (20 mL) was added TFA (10 mmol) and the solution stirred at room temperature for 24 hours. Concentration of the solution followed by trituration in diethyl ether provided the desired product as a white solid in 75% yield.

###### 4-(2-(2-((2-((4-ethylphenyl)amino)-2-oxoethyl)thio)-6,7-dimethoxy-4-oxoquinazolin-3(4H)-yl)ethyl)piperazin-1-ium (10m)

White solid; yield 75%; ^1^H NMR (500 MHz, DMSO) δ 10.34 (s, 1H), 8.59 (s, 2H), 7.54 (d, *J* = 8.2 Hz, 2H), 7.38 (s, 1H), 7.16 (d, *J* = 8.2 Hz, 2H), 6.94 (s, 1H), 4.20 (d, *J* = 4.6 Hz, 4H), 3.86 (s, 3H), 3.81 (s, 3H), 3.12 (s, 4H), 2.74 (d, *J* = 6.9 Hz, 6H), 1.16 (t, *J* = 7.6 Hz, 3H); ^13^C NMR (126 MHz, DMSO) δ 165.19, 159.48, 154.33, 153.65, 147.58, 142.49, 138.40, 136.25, 127.49, 118.70, 111.08, 106.19, 104.92, 55.32, 55.25, 54.08, 49.02, 42.44, 41.12, 36.33, 27.07, 15.21; HRMS: calcd for C_26_H_33_N_5_O_4_S [M+H] m/z 512.2332, found 512.2333.

##### General procedure E: for the synthesis of compounds 10n–s

Triethylamine (1.5 mmol) was added to a suspension of intermediate **10m** (200 mg, 1 mmol) in dichloromethane (15 mL) and the mixture was stirred briefly. The appropriate alkylating or acylating agent-methyl iodide, ethyl bromide, methylsulfonyl chloride, 2-chloroacetonitrile, dimethylsulfamoyl chloride, or methylcarbamoyl chloride (1.5 mmol) was then added. Reaction progress was monitored by TLC. Upon completion, additional DCM and water were added, and the organic layer was separated, dried over anhydrous sodium sulfate, filtered, and concentrated. Purification by flash chromatography using hexane–ethyl acetate afforded compounds **10n–s** in 60–80% yield.

###### 4-(2-(2-((2-((4-ethylphenyl)amino)-2-oxoethyl)thio)-6,7-dimethoxy-4-oxoquinazolin-3(4H)-yl)ethyl)-N-methylpiperazine-1-carboxamide (10n)

White solid; yield 62%; ^1^H NMR (500 MHz, CDCl_3_) δ 9.72 (s, 1H), 7.56 (s, 1H), 7.34 (d, J = 8.2 Hz, 2H), 7.10 (d, J = 8.2 Hz, 2H), 7.00 (s, 1H), 4.39 (q, J = 4.7 Hz, 1H), 4.30 (s, 2H), 4.01 (d, J = 6.1 Hz, 8H), 3.35 (t, J = 5.0 Hz, 4H), 2.79 (t, J = 7.1 Hz, 5H), 2.70 – 2.48 (m, 6H), 1.18 (t, J = 7.6 Hz, 3H); ^13^C NMR (126 MHz, CDCl_3_) δ 166.47, 160.62, 158.32, 155.62, 148.98, 142.89, 140.47, 135.66, 128.41, 119.40, 112.60, 106.26, 105.77, 56.50, 56.37, 55.21, 53.04, 43.65, 35.97, 28.28, 27.63, 15.66; HRMS: calcd for C_28_H_36_N_6_O_5_S [M+H] m/z 569.2546, found 569.2570.

###### 2-((3-(2-(4-(N,N-dimethylsulfamoyl)piperazin-1-yl)ethyl)-6,7-dimethoxy-4-oxo-3,4-dihydroquinazolin-2-yl)thio)-N-(4-ethylphenyl)acetamide (10o)

White solid; yield 58%;^1^H NMR (500 MHz, CDCl_3_) δ 9.66 (s, 1H), 7.49 (s, 1H), 7.28 (d, J = 8.4 Hz, 2H), 7.03 (d, J = 8.4 Hz, 2H), 6.94 (s, 1H), 4.21 (s, 2H), 3.94 (d, J = 3.6 Hz, 8H), 3.17 (d, J = 5.1 Hz, 4H), 2.72 (s, 8H), 2.63 – 2.46 (m, 6H), 1.11 (t, J = 7.6 Hz, 3H); ^13^C NMR (126 MHz, CDCl_3_) δ 166.48, 160.60, 155.65, 149.00, 142.87, 140.49, 135.66, 128.43, 119.40, 112.60, 106.28, 105.75, 56.51, 56.39, 55.10, 52.86, 46.25, 38.22, 35.99, 28.28, 15.68; HRMS: calcd for C_28_H_38_N_6_O_6_S_2_ [M+H] m/z 619.2372, found 619.2393.

###### 2-((6,7-dimethoxy-3-(2-(4-(methylsulfonyl)piperazin-1-yl)ethyl)-4-oxo-3,4-dihydroquinazolin-2-yl)thio)-N-(4-ethylphenyl)acetamide (10p)

White solid; yield 65%; ^1^H NMR (500 MHz, CDCl_3_) δ 9.79 (s, 1H), 7.57 (s, 1H), 7.32 (d, J = 8.2 Hz, 2H), 7.09 (d, J = 8.2 Hz, 2H), 7.01 (s, 1H), 4.26 (s, 2H), 4.01 (d, J = 2.7 Hz, 8H), 3.16 (t, J = 4.9 Hz, 4H), 2.79 (d, J = 6.7 Hz, 2H), 2.67 (t, J = 4.9 Hz, 4H), 2.62 (s, 3H), 2.58 (d, J = 7.6 Hz, 2H), 1.18 (t, J = 7.6 Hz, 3H); ^13^C NMR (126 MHz, CDCl_3_) δ 166.50, 160.58, 155.72, 149.05, 142.88, 140.57, 135.66, 128.46, 119.40, 112.56, 106.27, 105.76, 56.52, 56.41, 54.93, 52.71, 35.95, 28.27, 15.67; HRMS: calcd for C_27_H_35_N_5_O_6_S_2_ [M+H] m/z 590.2107, found 590.2122.

###### 2-((3-(2-(4-(cyanomethyl)piperazin-1-yl)ethyl)-6,7-dimethoxy-4-oxo-3,4-dihydroquinazolin-2-yl)thio)-N-(4-ethylphenyl)acetamide (10q)

White solid; yield 65%; ^1^H NMR (500 MHz, CDCl_3_) δ 9.66 (s, 1H), 7.50 (s, 1H), 7.30 – 7.25 (m, 2H), 7.03 (d, J = 8.2 Hz, 2H), 6.94 (s, 1H), 4.21 (s, 2H), 3.94 (d, J = 7.0 Hz, 8H), 3.37 (s, 2H), 2.72 – 2.45 (m, 12H), 1.11 (t, J = 7.6 Hz, 3H); ^13^C NMR (126 MHz, CDCl_3_) δ 166.51, 160.61, 155.62, 155.49, 148.97, 142.88, 140.46, 135.68, 128.42, 119.41, 114.73, 112.62, 106.30, 105.75, 56.50, 56.38, 54.94, 52.85, 51.72, 45.80, 35.95, 28.28, 15.68; HRMS: calcd for C_28_H_34_N_6_O_4_S [M+H] m/z 551.2440, found 551.2454.

###### 2-((6,7-dimethoxy-3-(2-(4-methylpiperazin-1-yl)ethyl)-4-oxo-3,4-dihydroquinazolin-2-yl)thio)-N-(4-ethylphenyl)acetamide (10r)

White solid; yield 60%; ^1^H NMR (500 MHz, CDCl_3_) δ 9.70 (s, 1H), 7.50 (s, 1H), 7.27 (d, *J* = 8.1 Hz, 2H), 7.03 (d, *J* = 8.1 Hz, 2H), 6.93 (s, 1H), 4.20 (t, *J* = 7.3 Hz, 2H), 3.94 (d, *J* = 3.7 Hz, 8H), 2.81 – 2.25 (m, 12H), 2.20 (s, 3H), 1.10 (t, *J* = 7.5 Hz, 3H); ^13^C NMR (126 MHz, CDCl_3_) δ 166.60, 160.57, 155.70, 155.59, 148.93, 142.88, 140.46, 135.69, 128.42, 119.40, 112.66, 106.31, 105.71, 56.50, 56.38, 55.14, 54.91, 42.71, 35.96, 28.28, 15.70; HRMS: calcd for C_27_H_35_N_5_O_4_S [M+H] m/z 526.2484, found 526.2488.

###### N-(4-ethylphenyl)-2-((3-(2-(4-ethylpiperazin-1-yl)ethyl)-6,7-dimethoxy-4-oxo-3,4-dihydroquinazolin-2-yl)thio)acetamide (10s)

White solid; yield 78%; ^1^H NMR (500 MHz, CDCl_3_) δ 9.70 (s, 1H), 7.56 (s, 1H), 7.32 (d, *J* = 8.2 Hz, 2H), 7.10 (d, *J* = 8.1 Hz, 2H), 7.01 (s, 1H), 4.25 (t, *J* = 5.9 Hz, 2H), 4.06 – 3.98 (m, 8H), 3.49 – 3.31 (m, 2H), 3.01 (s, 2H), 2.85 (dt, *J* = 22.2, 6.7 Hz, 6H), 2.57 (q, *J* = 7.7 Hz, 4H), 1.34 (s, 3H), 1.32 (s, 3H); ^13^C NMR (126 MHz, CDCl_3_) δ 166.64, 160.62, 155.76, 155.37, 149.06, 142.83, 140.70, 135.63, 128.50, 119.33, 112.54, 106.29, 105.76, 56.52, 56.41, 54.61, 51.70, 51.02, 50.01, 45.67, 42.54, 35.87, 28.25, 15.69, 8.50; HRMS: calcd for C_28_H_37_N_5_O_4_S [M+H] m/z 540.2645, found 540.2665.

##### General procedure F: for the synthesis of compound 10t

Compound **10m** (200 mg, 1.0 mmol) was suspended in dichloromethane (20 mL), and triethylamine (1.5 mmol) was added. The mixture was stirred at room temperature for 10–15 min, followed by the addition of acetic acid (3 mmol) and acetone (500 µL). After stirring for an additional 10 min, sodium triacetoxyborohydride (1.5 mmol) was added. Reaction completion was confirmed by TLC. The mixture was diluted with DCM and washed with saturated sodium bicarbonate solution. The organic phase was dried over sodium sulfate, filtered, concentrated, and purified by flash chromatography (hexane–ethyl acetate) to yield compound **10t** in 68% yield.

###### N-(4-ethylphenyl)-2-((3-(2-(4-isopropylpiperazin-1-yl)ethyl)-6,7-dimethoxy-4-oxo-3,4-dihydroquinazolin-2-yl)thio)acetamide (10t)

White solid; yield 60%; ^1^H NMR (500 MHz, CDCl_3_) δ 9.78 (s, 1H), 7.57 (s, 1H), 7.34 (d, *J* = 8.5 Hz, 2H), 7.09 (d, *J* = 8.4 Hz, 2H), 7.00 (s, 1H), 4.28 (t, *J* = 7.3 Hz, 2H), 4.00 (d, *J* = 3.7 Hz, 8H), 2.78 – 2.52 (m, 13H), 1.18 (t, *J* = 7.6 Hz, 3H), 1.06 (d, *J* = 6.5 Hz, 6H); ^13^C NMR (126 MHz, CDCl_3_) δ 166.60, 160.57, 155.70, 155.58, 148.93, 142.88, 140.43, 135.71, 128.41, 119.41, 112.67, 106.32, 105.72, 56.50, 56.38, 55.19, 48.59, 42.67, 35.98, 28.28, 18.41, 15.69; HRMS: calcd for C_29_H_39_N_5_O_4_S [M+H] m/z 554.2801, found 554.2816.

##### General procedure G: for the synthesis of compound 10u

Triethylamine (1.5 mmol) was added to a suspension of compound **10m** (200 mg, 1.0 mmol) in dichloromethane (500 µL) and stirred briefly. Acetic anhydride (1.5 mmol) was then introduced, resulting in a cloudy reaction mixture. Progress was monitored by TLC. Upon completion, the mixture was diluted with DCM and washed with water. The organic layer was dried over anhydrous sodium sulfate, filtered, and concentrated under reduced pressure. Purification by flash chromatography using hexane–ethyl acetate afforded compound **10u** in 65% yield.

###### 2-((3-(2-(4-acetylpiperazin-1-yl)ethyl)-6,7-dimethoxy-4-oxo-3,4-dihydroquinazolin-2-yl)thio)-N-(4-ethylphenyl)acetamide (10u)

White solid; yield 62%; ^1^H NMR (500 MHz, CDCl_3_) δ 9.65 (s, 1H), 7.49 (s, 1H), 7.31 – 7.24 (m, 2H), 7.06 – 7.00 (m, 2H), 6.94 (s, 1H), 4.22 (s, 2H), 3.97 – 3.90 (m, 8H), 3.51 (s, 2H), 3.35 (s, 2H), 2.69 (s, 2H), 2.50 (dt, *J* = 15.1, 7.4 Hz, 6H), 1.99 (s, 3H), 1.11 (t, *J* = 7.6 Hz, 3H); ^13^C NMR (126 MHz, CDCl_3_) δ 168.91, 166.47, 160.61, 155.65, 149.00, 142.88, 140.50, 135.65, 128.43, 119.37, 112.60, 106.27, 105.75, 56.51, 56.39, 55.18, 53.40, 53.14, 35.97, 28.28, 21.31, 15.67; HRMS: calcd for C_28_H_35_N_5_O_5_S [M+H] m/z 554.2437, found 554.2441.

##### General procedure H: for the synthesis of compound 10v

To a solution of **10d** (1 mmol) in methanol (20 mL) was added 10%-palladium carbon and the mixture was vigorously stirred under a hydrogen atmosphere at room temperature for 6 hr. The catalyst was filtered off through celite, the filtrate was concentrated under reduced pressure, and the obtained residue was purified by flash chromatography using dichloromethane–methanol afforded compound **10v** in 52% yield.

###### 2-((3-(4-aminophenethyl)-6,7-dimethoxy-4-oxo-3,4-dihydroquinazolin-2-yl)thio)-N-(4-ethylphenyl)acetamide (10v)

White solid; yield 52%; ^1^H NMR (500 MHz, CDCl_3_) δ 9.68 (s, 1H), 7.54 (s, 1H), 7.30 – 7.26 (m, 2H), 7.01 (dd, *J* = 23.3, 8.1 Hz, 4H), 6.94 (s, 1H), 6.57 (d, *J* = 7.8 Hz, 2H), 4.23 – 4.17 (m, 2H), 3.97 – 3.90 (m, 8H), 2.89 (dd, *J* = 9.4, 6.6 Hz, 2H), 2.51 (q, *J* = 7.5 Hz, 2H), 1.11 (t, *J* = 7.6 Hz, 3H); ^13^C NMR (126 MHz, CDCl_3_) δ 166.64, 160.51, 155.67, 155.56, 148.90, 142.92, 140.41, 135.79, 129.87, 128.40, 119.45, 115.48, 112.77, 106.33, 105.71, 56.51, 56.39, 47.10, 35.87, 33.29, 28.29, 15.69; HRMS: calcd for C_28_H_30_N_4_O_4_S [M+H] m/z 519.2066, found 519.2047.

#### Biological evaluation

##### Microscale thermophoresis (MST) assay

Microscale thermophoresis (MST) was performed using Monolith NT.115 instrument (NanoTemper Technologies) to evaluate the interaction between test compounds and human CHI3L1. Recombinant His-tagged human CHI3L1 (Cat. #CH1-H5228, Acro Biosystems) was fluorescently labeled using the Monolith His-Tag Labeling Kit RED-tris-NTA 2nd Generation (Cat. #MO-L018) in HEPES buffer (10 mM HEPES, 150 mM NaCl, 0.1% Pluronic F-127). The labeled protein was diluted to a working concentration of 80-60 nM. Test compounds were serially diluted in assay buffer (10 mM HEPES, 150 mM NaCl, 0.1% Pluronic F-127, 1 mM TCEP, 4% DMSO, pH 7.4) over 12-16 concentrations ranging from 500 µM to low nanomolar levels. Labeled CHI3L1 and compound dilutions were mixed in a 1:1 ratio and incubated for 30 minutes at room temperature. Samples were centrifuged at 1000 × g for 30 seconds and loaded into standard capillaries. MST measurements were recorded using 60–80% LED power (red channel) and medium to high IR-laser settings. Binding data were analyzed using MO.Affinity Analysis v2.3 (NanoTemper Technologies) and reported as normalized fluorescence change (Fnorm) upon ligand binding.^36^

##### Evaluation of PK and physicochemical properties

These experiments were conducted following our previously reported methods.^1^ The evaluations included LogD7.4 determination, microsomal stability, kinetic solubility, and cytotoxicity profiling across multiple cell lines. Solubility was assessed using UV-visible spectrophotometry.

##### GBM spheroids evaluation

GBM spheroids were generated following established protocols.^35^ U-87 MG cells (ATCC) were cultured in DMEM supplemented with 4.5 g/L glucose, 2 mM L-glutamine, penicillin, and streptomycin. HMEC-1 cells (ATCC) were maintained in MCDB131 medium enriched with 10 mM L-glutamine, 10 ng/mL FGF, and 1 µg/mL hydrocortisone. For spheroid formation, U-87 MG and HMEC-1 cells were co-cultured with macrophages in low-adhesion 96-well plates at a density of 2 × 10³ cells per well in 100 µL of medium containing test compounds (5, 10, or 25 µM) or control media. After 72 hours of incubation, cell viability was assessed using the CCK-8 assay, with absorbance measured at 450 nm using a microplate reader.

Migration assays were performed using a serum-free, flat-surface model. Geltrex (Gibco) diluted 1:50 in DMEM was added to 12-well plates and incubated overnight at 36 °C. Following removal of the supernatant, individual spheroids were placed in each well and allowed to settle for 90 minutes before fresh medium was added. Spheroids were pre-treated with 50 µM of the test compound for 24 hours prior to analysis. Migration was tracked over three days using ImageJ, and cell death in migrating cells was evaluated by Propidium Iodide staining after 72 hours. Additionally, spheroid mass was quantified using the W8 Physical Cytometer in a non-invasive, label-free manner. Assessment of phospho-STAT3 levels was determined by HTRF phospho-STAT3 kit from Revvity (Cat# 62AT3PET) using the manufacturer’s recommended protocol. All experiments were conducted in triplicate.

##### Molecular Docking and MD simulation

Molecular docking studies were performed using the Maestro suite from Schrödinger (Schrödinger Release 2024.1., Schrödinger, LLC, New York, NY). The three-dimensional crystal structure of CHI3L1 (PDB ID: 8R4X) was retrieved from the Protein Data Bank and prepared using the Protein Preparation Wizard, which included assignment of bond orders, addition of hydrogen atoms, optimization of hydrogen bonding networks, and energy minimization using the OPLS4 force field. Ligands **K284** and **10p** were prepared using LigPrep to generate appropriate ionization states at pH 7.0 ± 2.0 and tautomeric forms. The receptor grid was generated centered on the native ligand binding site with default parameters. Standard precision (SP) docking was performed using Glide, allowing flexible ligand conformations while keeping the receptor rigid. Visualization of the docked poses was carried out using Discovery Studio Visualizer (BIOVIA, Dassault Systèmes).

Molecular dynamics simulations of the **10p**/CHI3L1 complex were conducted using the Desmond molecular dynamics package with the OPLS4 force field. The system was solvated in a TIP3P water model using an orthorhombic periodic boundary box with a minimum distance of 10 Å between the protein and box edges and neutralized with appropriate counter ions. Following energy minimization and equilibration through a series of restrained simulations, production runs were performed for 100 nanoseconds at 300 K and 1.01325 bar using the NPT ensemble with the Nosé-Hoover thermostat and Martyna-Tobias-Klein barostat. MD was carried out at the default setting according to our previously published work. ^35,36^

## Results

### Chemistry

The target compounds **9a-v** were synthesized via a four-step sequence starting from 6,7-dimethoxy-2H-benzo[d][1,3]oxazine-2,4(1H)-dione (**2**). Initial amide formation with various amines (**3a-l**) gave intermediates **4a-l**, which were converted to dithiocarbamate derivatives (**5a-l**) using carbon disulfide and base under mild heating (**Scheme 1**). Separately, 4-ethylaniline (**6**) was acylated with bromoacyl bromide (**7a**) to yield amide intermediates **8**. Final products **10a-l** were obtained through nucleophilic substitution between intermediates **5a-l** and **8** in acetone under basic conditions, followed by chromatographic purification. Overall yields for the final step ranged from 70-80%.

Boc deprotection of compound **10l** afforded the compound 1**0m**. Compounds **9n–s** were synthesized by reacting **10m** with methanesulfonyl chloride or 2-bromoacetonitrile or dimethylsulfamoyl chloride or methyl chloroformate or ethyl iodide or methyl iodide in DCM using triethylamine (TEA) as a base, yielding the corresponding products in 50–80% (Scheme 2). For the synthesis of compounds **10t–u**, bearing ketone and isopropyl substituents, **10m** was treated with acetic acid and acetic anhydride/acetone, followed by reductive alkylation using sodium triacetoxyborohydride in the presence of triethylamine (TEA) in DCM, to afford the desired products (Scheme 2).

**Scheme 1.**
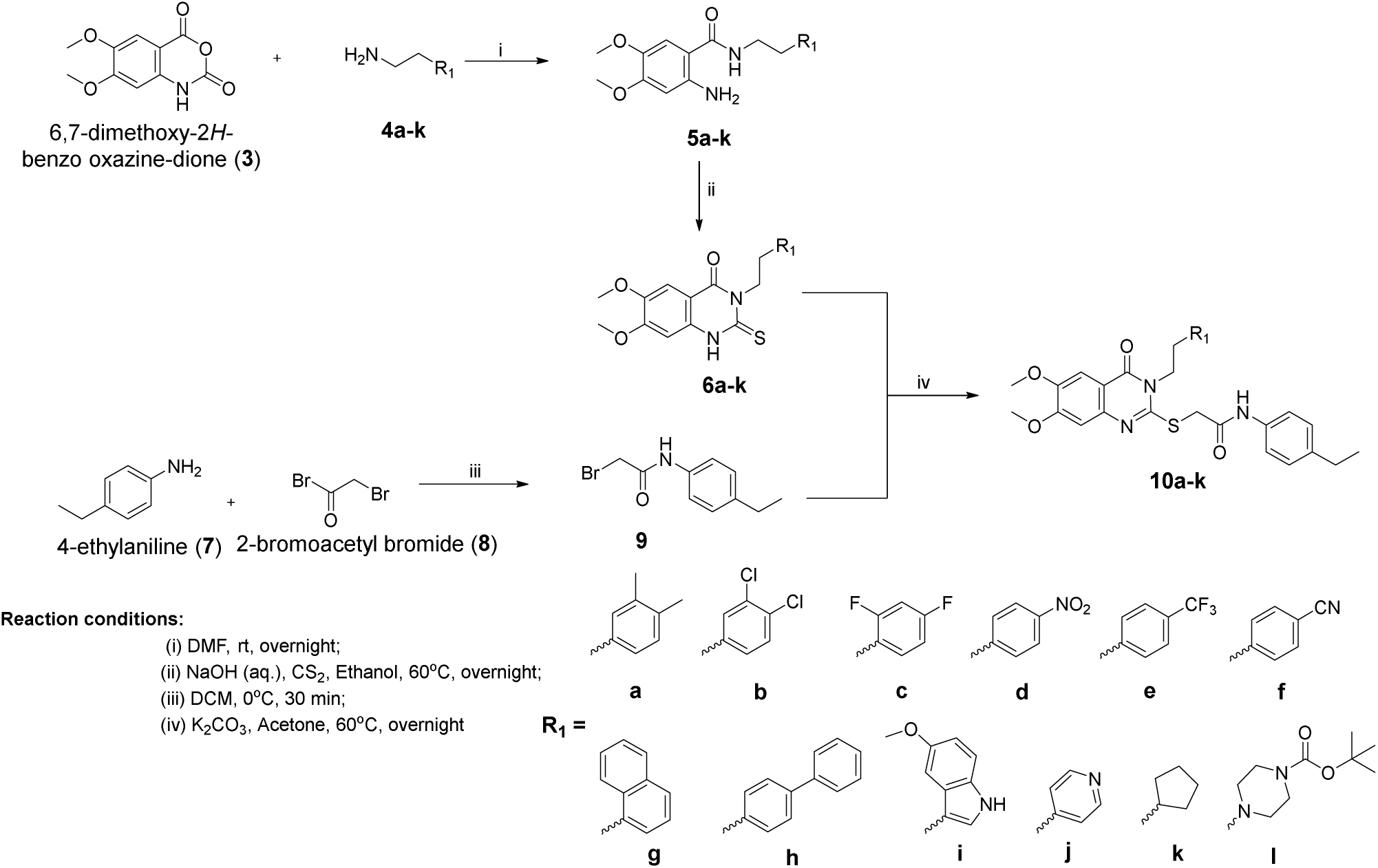
Synthesis of compounds **10a-k** as potential CHI3L1 inhibitors.

**Scheme 2.**
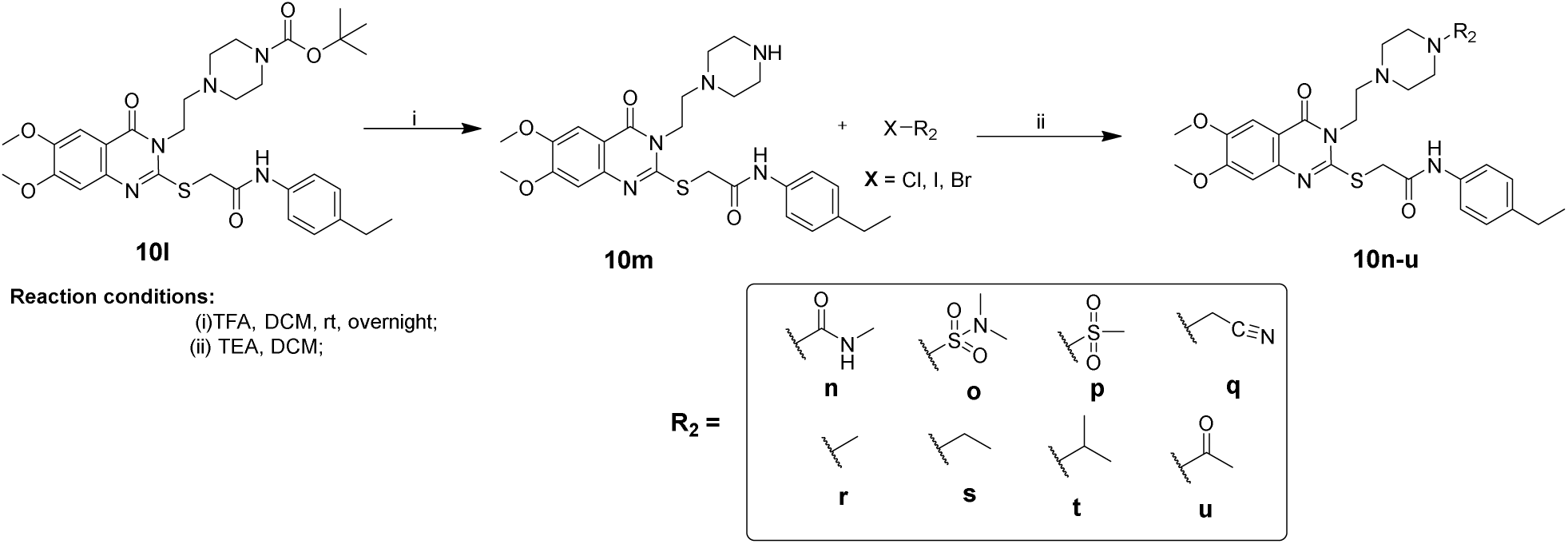
Synthesis of compounds **10n-v** as potential CHI3L1 inhibitors.

**Scheme 3.**
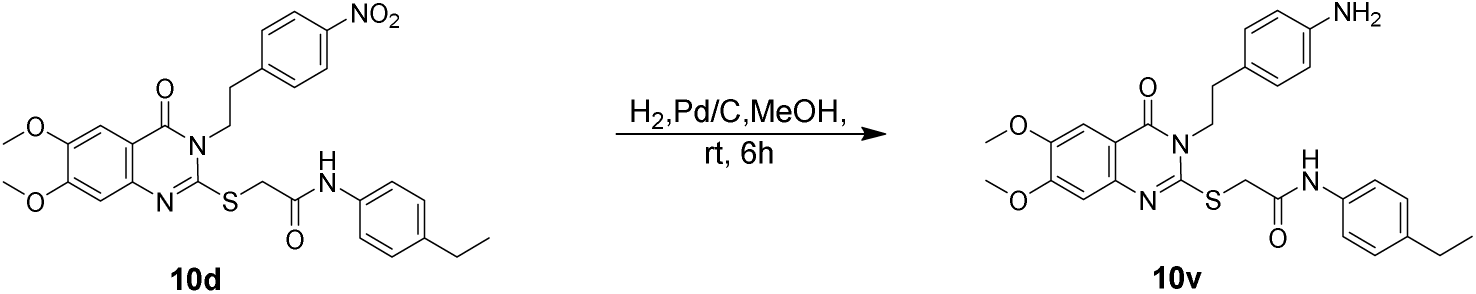
Synthesis of compounds **10w** from **10m**.

Compound **10v** was obtained from **10d** by catalytic hydrogenation of the nitro group to the corresponding amino group using Pd/C in methanol under a hydrogen atmosphere (Scheme 3).

### Biophysical Assessment of CHI3L1 Binding Affinity

#### Concentration-Dependent Binding Analysis using Monolith: Binding Affinity Screening Platform

All the newly synthesized compounds were then subjected to dose-response analyses to confirm their direct binding affinities to CHI3L1. Out of 21 compounds, eight compounds displayed clear dose-dependent binding curves, strongly suggesting specific interactions (Figure 2). Replacement of the 3,4-dimethoxyphenyl group with a biphenyl moiety led to a pronounced reduction in CHI3L1 binding affinity, with the *K*_d_ increasing from 13 µM to 384 µM for compound **10g** (Figure 2), indicating poor tolerance for increased aromatic bulk at this position. Substitution with a *p*-cyano phenyl group in **10f** resulted in a moderate loss of affinity (*K*_d_ = 110 µM), suggesting that strongly electron-withdrawing substituents are only partially accommodated within the binding site. Several additional analogues bearing differently substituted phenyl rings did not yield measurable binding data; however, this was primarily attributed to poor aqueous solubility, as these compounds precipitated under assay conditions, precluding reliable *K*_d_ determination. Consequently, the absence of detectable binding for these analogues should not be interpreted as definitive evidence of a lack of intrinsic affinity. In contrast, replacement of the aromatic moiety with a piperazine group was well tolerated, affording compound **10m** with a *K*_d_ value of 19.1 µM, highlighting the benefit of increased polarity and conformational flexibility at this position.

**Figure 2.**
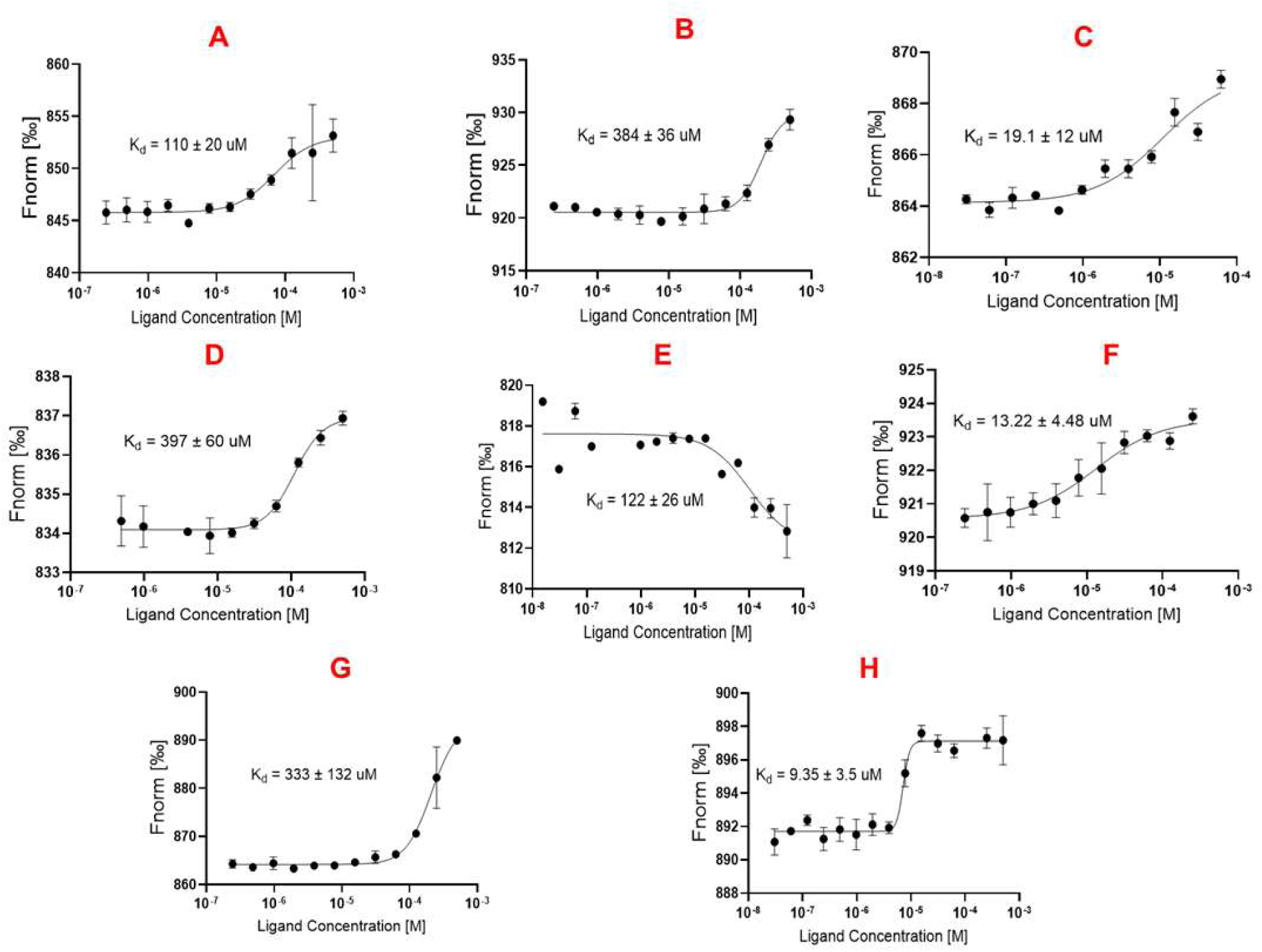
Dose-response curves of (A) **10f**, (B) **10g**, (C) **10m,** (D) **10n,** (E) **10o,** (F) **10p**, (G) **10r,** (H) **10u** to hCHI3L1 measured by MST.

Encouraged by this result, further derivatization of the piperazine secondary amine was undertaken to probe steric and electronic tolerance. Introduction of an *N*-methyl substituent (**10r**) resulted in a substantial decrease in binding affinity (*K*_d_ = 333 µM), while no measurable binding was observed for the corresponding *N*-ethyl and *N*-isopropyl analogues (**10s** and **10t**), indicating a strong sensitivity to steric bulk in this region of the binding pocket. Installation of carbonyl-containing substituents produced divergent effects: the *N*-acyl methylamide (*N*-CONHMe) in **10n** exhibited weak binding (*K*_d_ = 397 µM), whereas introduction of a methyl ketone substituent (*N*–C(=O)Me) in **10u** was well tolerated and yielded the most potent analogue in this series (*K*_d_ = 9.35 µM). Sulfonyl substitution further modulated affinity, with the *N,N*-dimethylsulfonamide analogue (**10o**) displayed reduced binding (*K*_d_ = 122 µM), while the methylsulfonyl derivative (**10p**) restored potency (*K*_d_ = 13.22 µM).

Overall, these SAR trends indicate that the CHI3L1 binding site imposes strict steric constraints and disfavors rigid aromatic expansion, while accommodating flexible, moderately polar substituents. Substituent identity at the piperazine nitrogen plays a critical role in binding, with small, planar, and appropriately polarized groups being favored over bulky alkyl or strongly hydrogen-bonding functionalities. Collectively, these findings define key scaffold-dependent constraints of the **K284** series and provide a framework for future optimization efforts aimed at improving drug-like properties while maintaining effective CHI3L1 engagement.

#### Evaluation of PK profile

The in vitro PK properties of **K284 (1)**, our previously reported CHI3L1 inhibitor **11g (2)**, and the newly developed series (**10m**, **10p**, and **10u**) are summarized in Table 1. While **K284** and **11g** provided useful benchmarks for CHI3L1 inhibition, both compounds exhibit limitations in solubility, metabolic stability, and safety that hindered further optimization. In contrast, the new piperazine analogs show a coordinated improvement in physicochemical and ADME properties, consistent with a more favorable profile for CNS-directed drug development.

**Table 1.** Quantification of CHI3L1 binding affinity for compounds using MST.

| Compound | $K_d$ ($\mu\text{M}$) | Compound | $K_d$ ($\mu\text{M}$) |
| --- | --- | --- | --- |
| **10a** | ND | **10m** | $19.1 \pm 12$ |
| **10b** | ND | **10n** | $397 \pm 60$ |
| **10c** | ND | **10o** | $122 \pm 26$ |
| **10d** | ND<sup>a</sup> | **10p** | $13.22 \pm 4.48$ |
| **10e** | ND | **10q** | ND |
| **10f** | $110 \pm 20$ | **10r** | $333 \pm 132$ |
| **10g** | $384 \pm 36$ | **10s** | ND |
| **10h** | ND | **10t** | ND |
| **10i** | ND | **10u** | $9.35 \pm 3.5$ |
| **10j** | ND | **10v** | ND |
| **10k** | ND |  |  |

<sup>a</sup>Undetermined due to negligible binding.

Across the new series, reduced lipophilicity (LogD_7.4_ = 1.2-1.7) was accompanied by a pronounced enhancement in aqueous solubility under physiological conditions. Notably, **10p** demonstrated a dramatic increase in kinetic solubility (758 µM at pH 7.4), representing an approximately five-fold and nine-fold improvement relative to **11g** and **K284**, respectively (Table 2). This substantial gain in solubility addresses a key developability limitation of earlier CHI3L1 inhibitors and is expected to support improved formulation and systemic exposure.

**Table 2.** Assessment of the in vitro PK properties of K284, 11g, 10m, 10p, and 10u.^a^.

| Compound | **K284** | **11g** | **10m** | **10p** | **10u** |
| --- | --- | --- | --- | --- | --- |
| **LogD<sub>7.4</sub>** | 2.2 | 1.8 | 1.7 | 1.2 | 1.5 |
| **Kinetic solubility ($\mu\text{M}$,**<br>**PBS pH 7.4)** | 81 | 152 | 224 | 758 | 502 |
| **PAMPA permeability**<br>**(cm/s)** | $2.9 \times 10^{-6}$ | $4.6 \times 10^{-6}$ | $2.8 \times 10^{-6}$ | $2.2 \times 10^{-6}$ | $2.5 \times 10^{-6}$ |
| **Human PPB (%)** | 90.3 | 86.2 | 85.1 | 78.2 | 81.3 |
| **Mouse plasma t<sub>1/2</sub> (h)** | 1.4 | 1.9 | 2.1 | 2.9 | 2.8 |
| **Human plasma t<sub>1/2</sub> (h)** | 1.8 | 2.4 | 2.5 | 3.3 | 3.1 |
| **Mouse microsomal t<sub>1/2</sub>**<br>**(h)** | 1.5 | 2.1 | 2.2 | 3.1 | 2.9 |
| **Human microsomal t<sub>1/2</sub>**<br>**(h)** | 1.9 | 2.6 | 2.8 | 3.7 | 3.1 |
| **Intrinsic clearance –**<br>**Mouse (mL/min/mg)** | 19.5 | 16.2 | 15.1 | 11.4 | 12.8 |
| **Intrinsic clearance –**<br>**Human (mL/min/mg)** | 21.6 | 20.3 | 19.5 | 14.2 | 16.3 |
| **hERG IC<sub>50</sub> (μM)** | 20.4 ± 1.3 | 41.5 ± 2.4 | 63.5 ± 5.4 | >100 | >100 |

<sup>a</sup>Values are mean ± SD (n=3).

Improvements in physicochemical properties translated into enhanced metabolic stability in both mouse and human systems. The new analogs, particularly **10p** and **10u**, exhibited extended plasma and microsomal half-lives alongside reduced intrinsic clearance compared with both **K284** and **11g**, indicating decreased susceptibility to hepatic metabolism across species (Table 2). These features suggest a greater likelihood of achieving sustained target engagement in vivo, which is critical for therapeutic intervention in GBM.

Although passive permeability measured by PAMPA was modestly reduced for the new compounds relative to **11g**, all values remained within a range compatible with passive diffusion (Table 2). In the context of markedly improved solubility and metabolic stability, the observed permeability is considered acceptable for CNS exposure and BBB penetration. Importantly, the new series also demonstrated a pronounced reduction in hERG liability, with **10p** and **10u** showing no detectable inhibition at concentrations up to 100 µM, representing a significant safety improvement over both comparator compounds (Table 2).

Collectively, these data highlight **10p** as the most balanced CHI3L1 inhibitor in the series, combining substantially improved solubility, enhanced metabolic stability, and minimized cardiac risk while maintaining permeability consistent with CNS-focused drug development for glioblastoma therapy.

#### Evaluation of the new series as potential GBM therapeutics

Three-dimensional (3D) spheroid models have gained increasing recognition as advanced experimental systems for the evaluation of GBM therapeutics, as they recapitulate tumor architecture and microenvironmental complexity than conventional two-dimensional (2D) monolayer cultures. Unlike 2D systems, 3D spheroids reproduce key physiological features of solid tumors, including gradients of oxygen, nutrients, and signaling molecules, as well as multicellular interactions that govern tumor growth, invasion, and therapeutic response. These features are particularly relevant for GBM, a highly aggressive and heterogeneous brain tumor characterized by extensive vascularization and pronounced infiltration of immune cells, especially macrophages.

To evaluate the functional impact of CHI3L1 inhibition in a physiologically relevant context, we employed a previously established multicellular GBM spheroid model composed of U-87 MG GBM cells, human microvascular endothelial cells (HMEC-1), and macrophages. This model reflects key elements of the GBM tumor microenvironment, including endothelial-tumor crosstalk and immune modulation, and provides a robust platform for assessing the pharmacological effects of CHI3L1-targeting small molecules.

Using this system, we compared the reference compound **K284**, our previously reported inhibitor **11g**, and the newly developed new series (**10m**, **10p**, and **10u**). GBM spheroids were treated with increasing concentrations of each compound for 72 h, and cell viability was assessed (Figure 3). Consistent with prior observations, compound **11g** induced a clear, dose-dependent reduction in spheroid viability, whereas **K284** showed minimal effects at concentrations up to 25 µM. Among the new analogs, **10p** produced viability reductions comparable to, and in some cases slightly exceeding, those observed for **11g** across the tested concentration range. In comparison to **11g**, **10m** and **10u** were slightly less efficient in reducing the viability of the spheroids, indicating reduced functional activity in the 3D GBM context (Figure 3).

**Figure 3.**
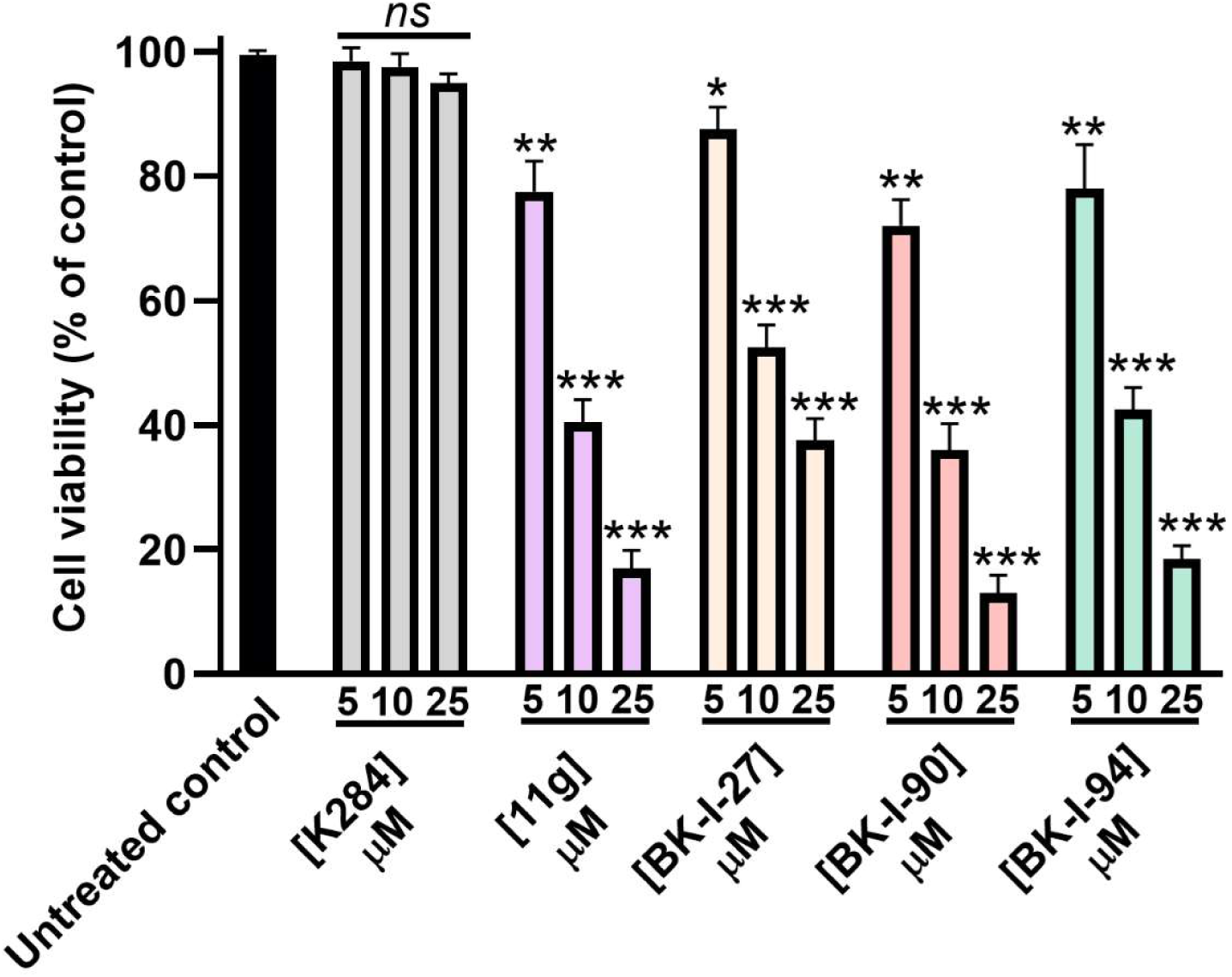
Cell viability of GBM spheroids (as % of untreated control) upon incubation with increasing concentrations of the tested compounds (**11g**, **K284**, **10m**, **10p**, **and 10u**) after 72 h incubation. * *p* < 0.05, ** *p* < 0.01, *** *p* < 0.001, and (*ns*) denotes nonsignificant relative to untreated control. Data are representative of three independent experiments.

To assess gross morphological and structural effects, spheroid mass was quantified following treatment at 10 µM (Figure 4). Treatment with **11g** resulted in a pronounced decrease in spheroid weight (∼50%), consistent with extensive cell death and disruption of spheroid integrity. Notably, both **10p** and **10u** induced a similar or greater reduction in spheroid mass, indicating robust impairment of tumor microtissue cohesion. In contrast, **10m** caused ∼35% reduction in spheroid weight, while **K284** again exhibited minimal impact (Figure 4). The close correspondence between reductions in viability and spheroid mass underscores the strong anti-tumor effects of **10p** and **10u** in this multicellular GBM model.

**Figure 4.**
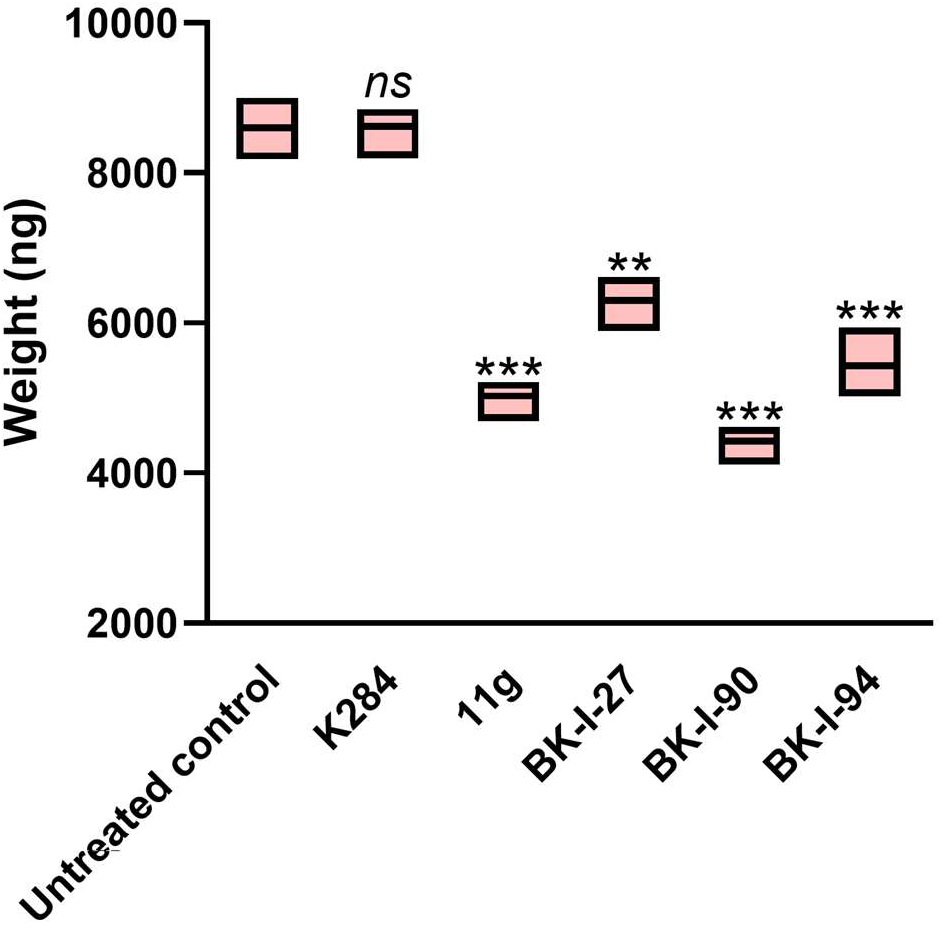
Changes in the weight of GBM spheroids upon incubation with 10 mM of the tested compounds (**11g**, **K284**, **10m**, **10p**, **and 10u**) after 72 h incubation. ** *p* < 0.01, *** *p* < 0.001, and (*ns*) denotes nonsignificant relative to untreated control. Data are representative of three independent experiments.

Given the highly invasive nature of GBM, we further examined compound effects on spheroid migration using a single concentration of 10 µM (Figure 5). As previously reported, compound **11g** significantly suppressed GBM spheroid migration (∼60%), suggesting interference with pathways governing cell motility and invasion. Importantly, **10m**, **10p** and **10u** demonstrated comparable inhibition of spheroid migration relative to **11g**, whereas **K284** had no significant effect on migration under these conditions. These results highlight the ability of **10m**, **10p** and **10u** to combine cytotoxic and anti-invasive activity in a physiologically relevant 3D tumor model.

**Figure 5.**
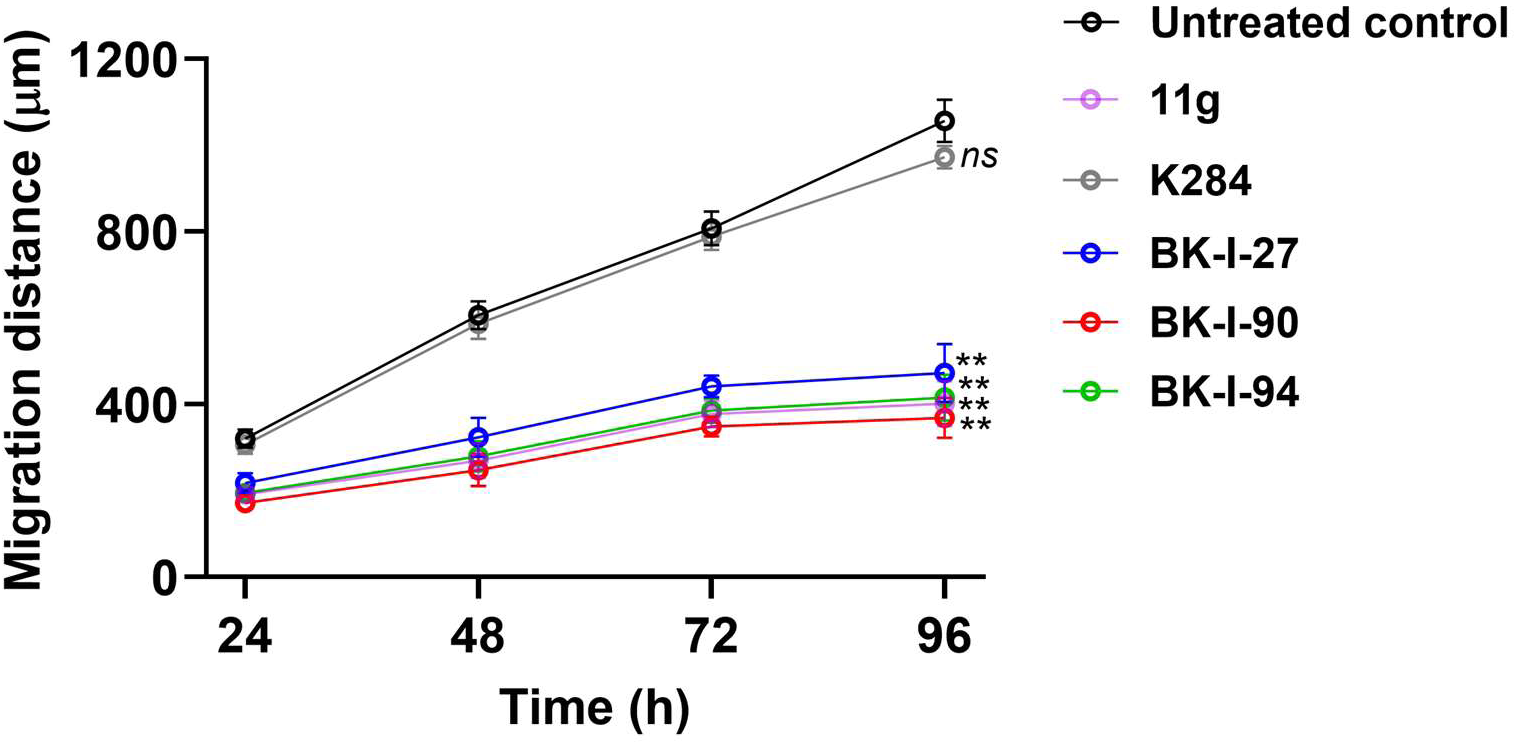
Cell migration distance (mm) of GBM spheroids upon incubation with 10 mM of the tested compounds (**11g**, **K284**, **10m**, **10p**, **and 10u**) after 96 h incubation. ** *p* < 0.01 and (*ns*) denotes nonsignificant relative to untreated control. Data are representative of three independent experiments.

Collectively, these data establish **10p** and **10u** as functionally superior CHI3L1 inhibitors relative to both the literature reference compound **K284** and the previously reported compound **11g**, while 10m displays slightly reduced efficacy in the 3D GBM setting. Building on prior mechanistic insights linking CHI3L1 inhibition to suppression of STAT3 signaling in GBM, these results support the advancement of **10p**, in particular, as a lead compound for further in vivo evaluation. Its strong performance across PK profile, spheroid viability, and invasion assays in a multicellular 3D model suggests effective modulation of CHI3L1-driven tumor biology in a context that closely mimics in vivo GBM behavior.

#### Molecular docking and MD studies

The molecular docking studies revealed distinct binding modes between the parent compound **K284** and its derivative **10p** in the CHI3L1 binding pocket. Both compounds exhibited favorable interactions with key residues in the active site, though with notable differences in their binding orientations and interaction profiles (Figure 6). **K284** demonstrated multiple hydrogen bonding interactions with ARG263, HIS209 and TYR141 along with extensive hydrophobic contacts involving PHE261, GLU290, TRP99, LEU140, and ASP207 (Figure 6A, B). In contrast, **10p** showed a reoriented binding pose with strong interactions involving three hydrogen bonds with ASP207, TRP352 and TYR141. Additional interactions were established between **10p** and MET204, PHE261, THR293, GLU290, ASN100, TRP99 and LEU140 which were predicted to further stabilize the compound within the binding site. Interestingly, both investigated compounds (**K284** and **10p**) maintained interactions with TRP99, TYR141, ASP207 and PHE261 indicating that interactions with these residues is pivotal for the binding.

**Figure 6.**
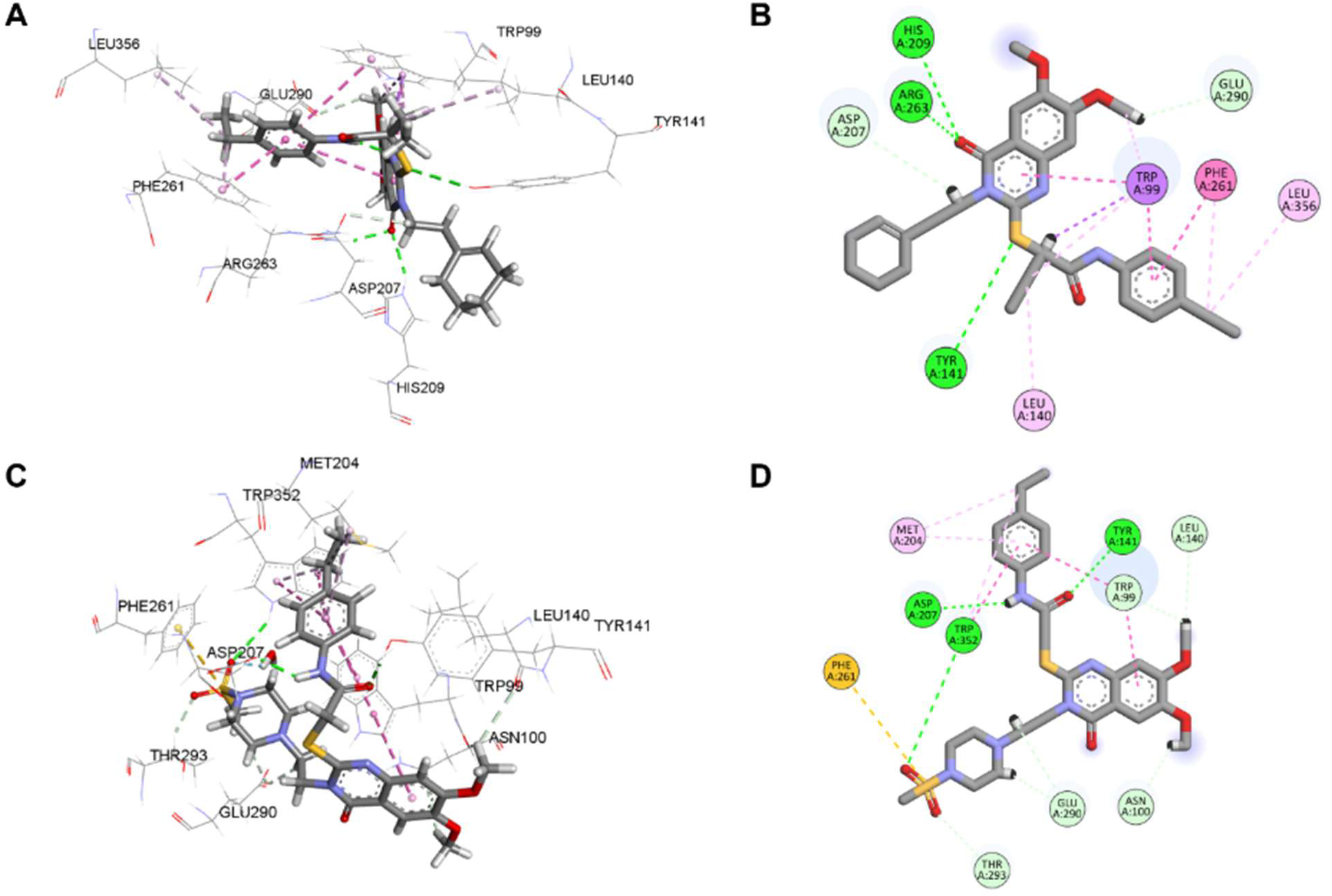
Molecular docking analysis of K284 and 10p binding to CHI3L1. (A) Three-dimensional representation of **K284** in the CHI3L1 binding pocket. (B) Two-dimensional ligand interaction diagram of **K284** showing amino acid residues within the binding pocket. (C) Three-dimensional binding mode of **10p** within the CHI3L1 active site. (D) Two-dimensional interaction map of **10p** showing the network of residues stabilizing the compound.

To assess the stability of the **10p** /CHI3L1 complex, molecular dynamics simulations were performed over 100 nanoseconds. The RMSD analysis demonstrated that the protein-ligand complex achieved equilibration after approximately 10 nanoseconds, maintaining stability throughout the simulation with a mean RMSD of 1.02 Å and minimal deviation (±1σ: 0.08 Å) (Figure 7A). The rolling average showed slight fluctuations around 40 nanoseconds but quickly returned to baseline, indicating transient conformational sampling rather than instability. This low RMSD profile confirms the structural integrity of the complex and suggests strong binding affinity. The ligand contact analysis throughout the simulation revealed persistent interactions with TRP99 and LEU140 throughout the whole simulation (Figure 7B). This persistent contact indicates the predicted importance of these residues in stabilizing the ligand within the binding pocket. The interaction heatmap demonstrates dynamic flexibility in certain contacts while maintaining core stabilizing interactions, which is characteristic of biologically relevant binding and exhibited experimental binding activity.

**Figure 7.**
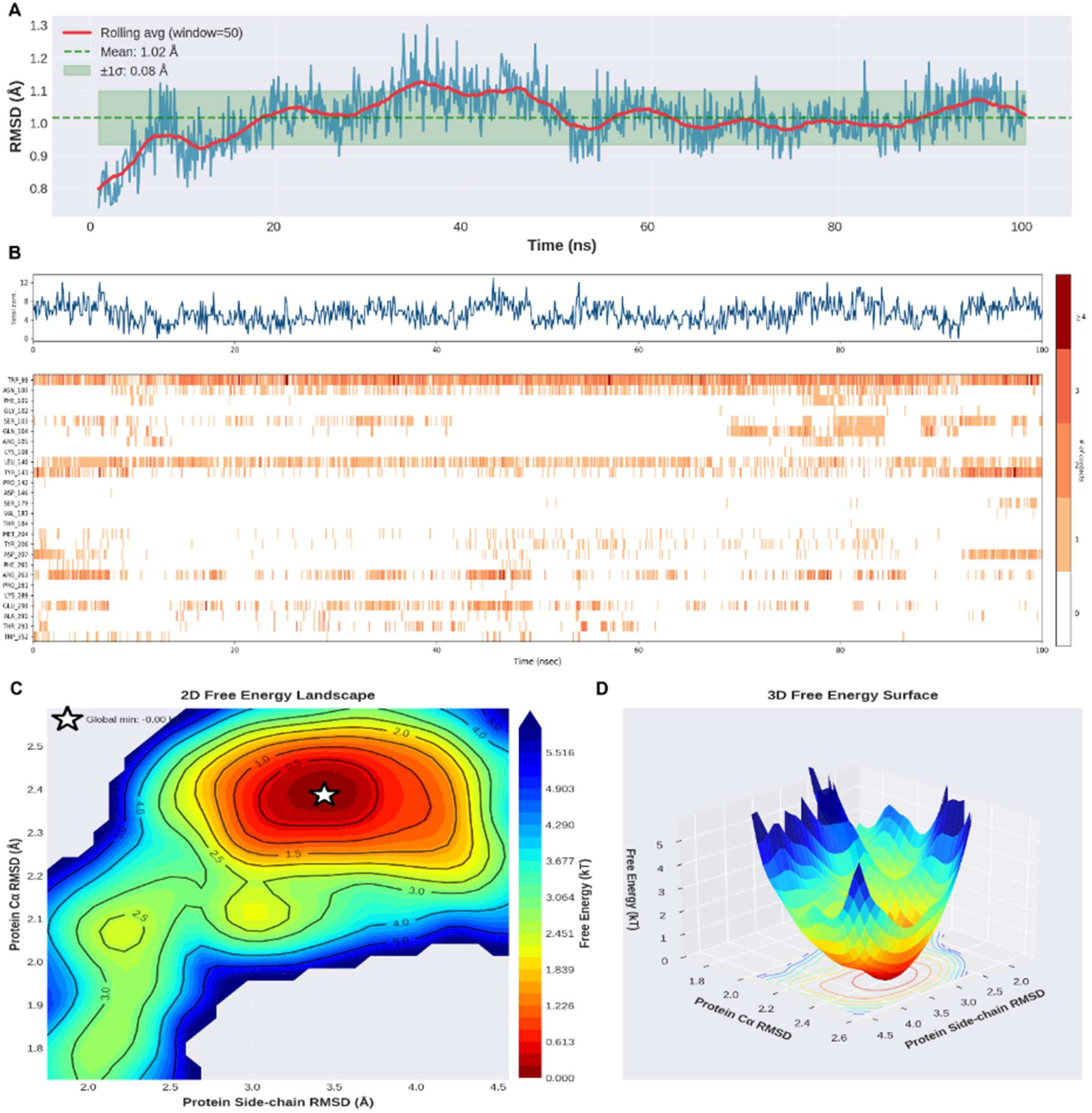
Molecular dynamics simulation analysis of the 10p-CHI3L1 complex over 100 nanoseconds. (A) RMSD of the protein-ligand complex as a function of simulation time. Blue bars represent instantaneous RMSD values, the red line shows the rolling average (window=50 frames), the dashed green line indicates the mean RMSD (1.02 Å), and the shaded green region represents ±1σ standard deviation (0.08 Å). (B) Top panel: ligand RMSD relative to the protein over time (nanoseconds). Bottom panel: residue interaction heatmap showing the frequency and strength of contacts between **10p** and individual CHI3L1 residues throughout the simulation. (C) Two-dimensional free energy landscape projected onto protein Cα RMSD (y-axis) and protein side-chain RMSD (x-axis). (D) Three-dimensional representation of the free energy surface.

The free energy landscape analysis provided additional insights into the conformational preferences of the **10p** /CHI3L1 complex. The 2D free energy landscape (Figure 7C) revealed a single, deep global energy minimum, indicating a highly favorable and stable binding conformation. The presence of this well-defined energy basin with minimal secondary minima suggests limited conformational heterogeneity and strong energetic favorability of the bound state. The 3D free energy surface (Figure 7D) further confirmed this observation, displaying a steep energy funnel converging to the global minimum with energy differences exceeding 5 kJ/mol from higher-energy states. This robust energy landscape indicates that **10p** adopts a thermodynamically stable conformation within the CHI3L1 binding site, with minimal likelihood of dissociation or conformational transitions during physiological timescales. Collectively, these computational studies demonstrate that **10p** is predicted to establish a stable, energetically favorable complex with CHI3L1. The molecular docking identified key binding determinants and showed improved interactions compared to **K284**, while the MD simulations confirmed the stability and persistence of these interactions over extended timescales. The favorable free energy profile and low RMSD fluctuations support the potential of **10p** as a potent CHI3L1 inhibitor warranting further experimental validation.

## Conclusion

This work defines critical structure–activity and structure–property relationships governing CHI3L1 inhibition within a **K284**-derived scaffold. Systematic modification revealed strict steric constraints at the aromatic substituent region, while introduction of a piperazine moiety enabled meaningful improvements in physicochemical and pharmacokinetic properties without loss of target engagement. These findings clarify the practical optimization space of this scaffold and identify design features that effectively mitigate developability limitations.

Notably, targeted structural changes overcame solubility-driven liabilities that restricted earlier CHI3L1 inhibitors, yielding compounds with substantially improved metabolic stability and safety profiles. These gains translated into consistent functional effects in a multicellular 3D GBM spheroid model that captures key aspects of tumor viability and invasion. Rather than relying on enhanced affinity alone, this study demonstrates the value of balanced optimization across binding, developability, and functional response.

Overall, compound **10p** exemplifies a well-balanced CHI3L1 modulator and provides a strong foundation for continued in vivo evaluation. More broadly, the insights reported here establish a rational framework for advancing CHI3L1-targeted small molecules as modulators of GBM-relevant tumor biology.

## Authorship contributions

Baljit Kaur: Writing – original draft, Formal analysis, Data curation, Conceptualization. Hossam Nada: Data curation, Conceptualization. Moustafa Gabr: Supervision, and Funding acquisition.

## Notes

The authors declare no competing financial interest.

