## Supporting Information for "Structural Optimization of CHI3L1 Inhibitors with Improved Pharmacokinetics and Functional Activity in 3D Glioblastoma Models"

**Table of Contents**

| 1. ^1^H NMR, ^13^C NMR of compounds | S2-S23 |
| --- | --- |
| 2. HRMS of compounds | S24-29 |

**1. ^1^H, and ^13^C NMR of compounds of 10a-v**
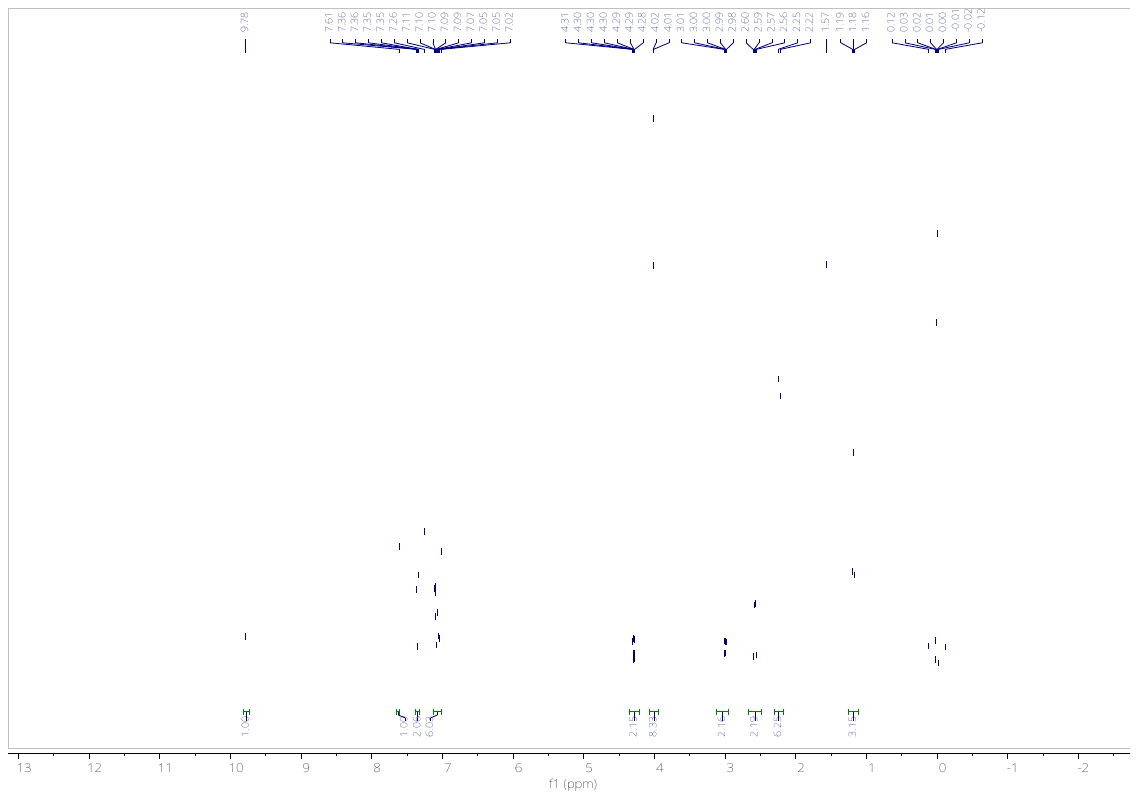

**Figure S1.** ^1^H NMR of **10a** in CDCl_3_.

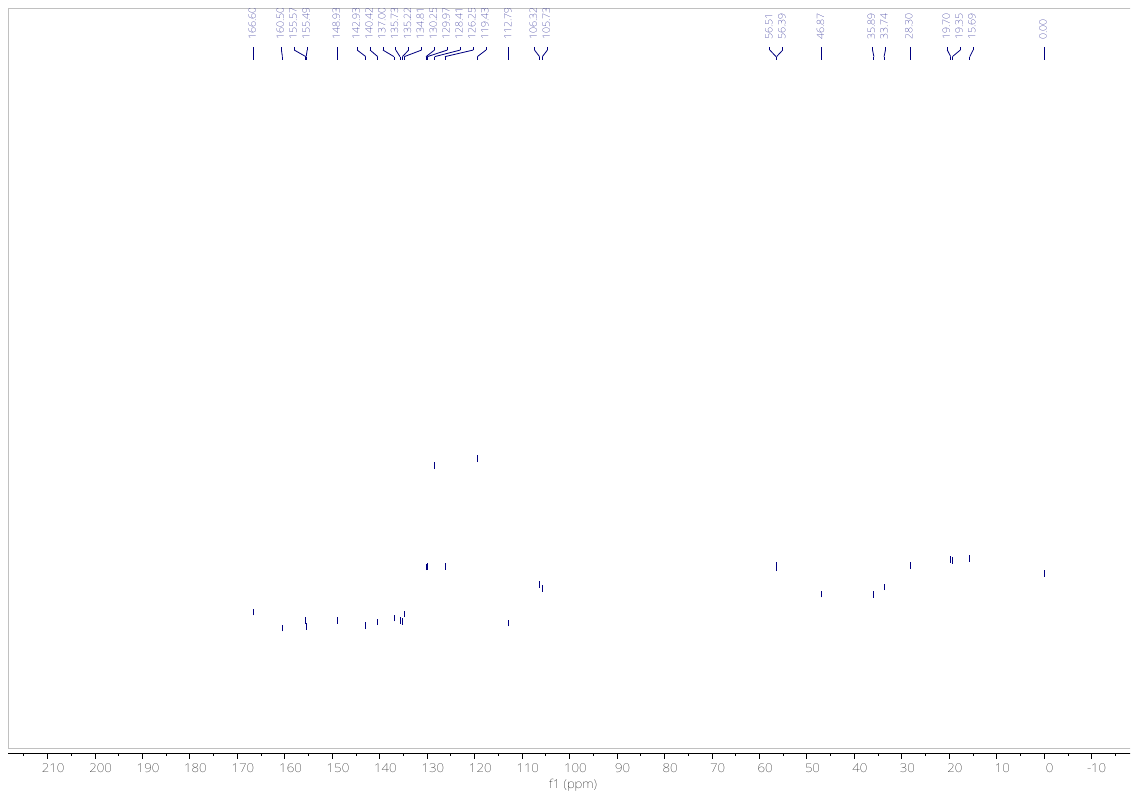

**Figure S2.** ^13^C NMR of **10a** in CDCl_3_.

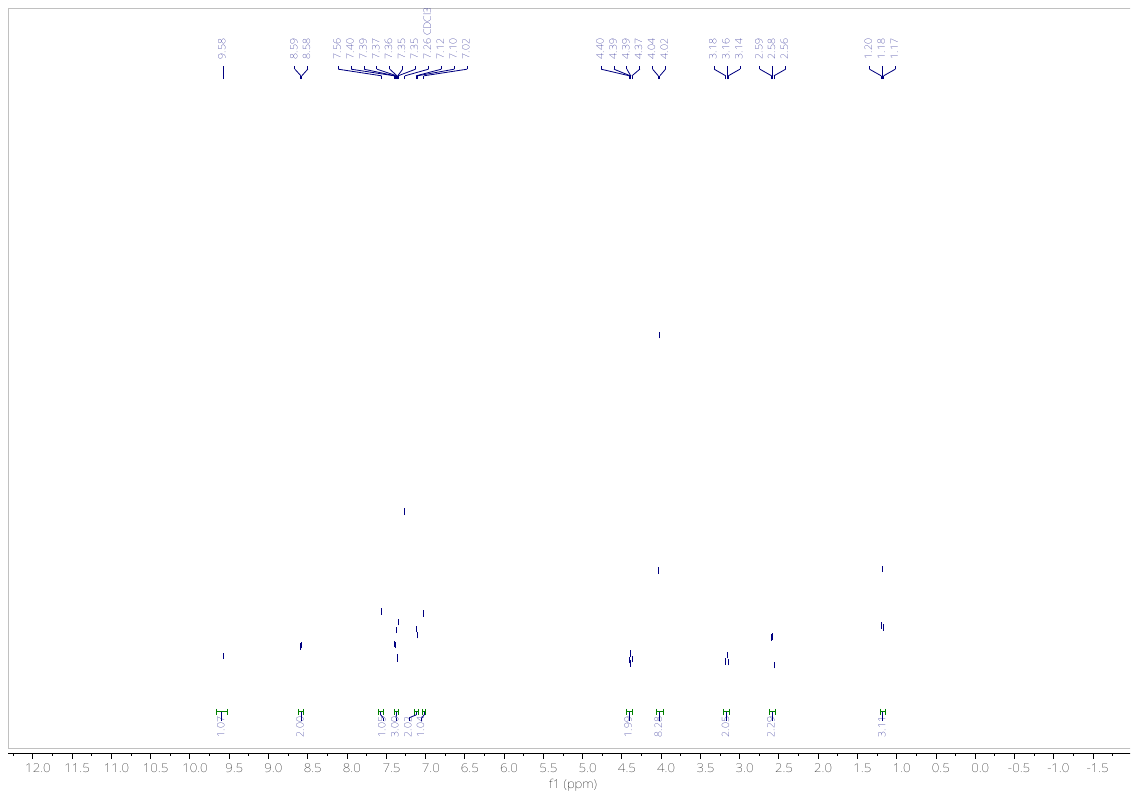

**Figure S3.** ^1^H NMR of **10b** in CDCl_3_.
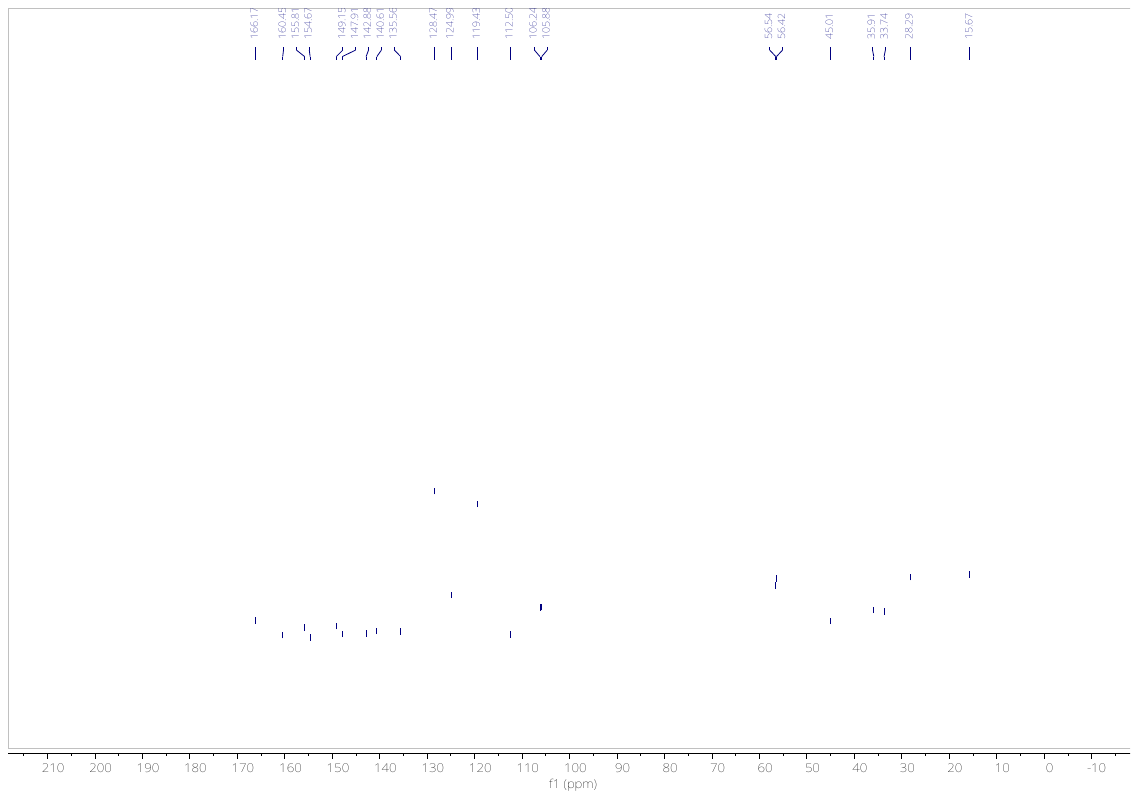

**Figure S4.** ^13^C NMR of **10b** in CDCl_3_.

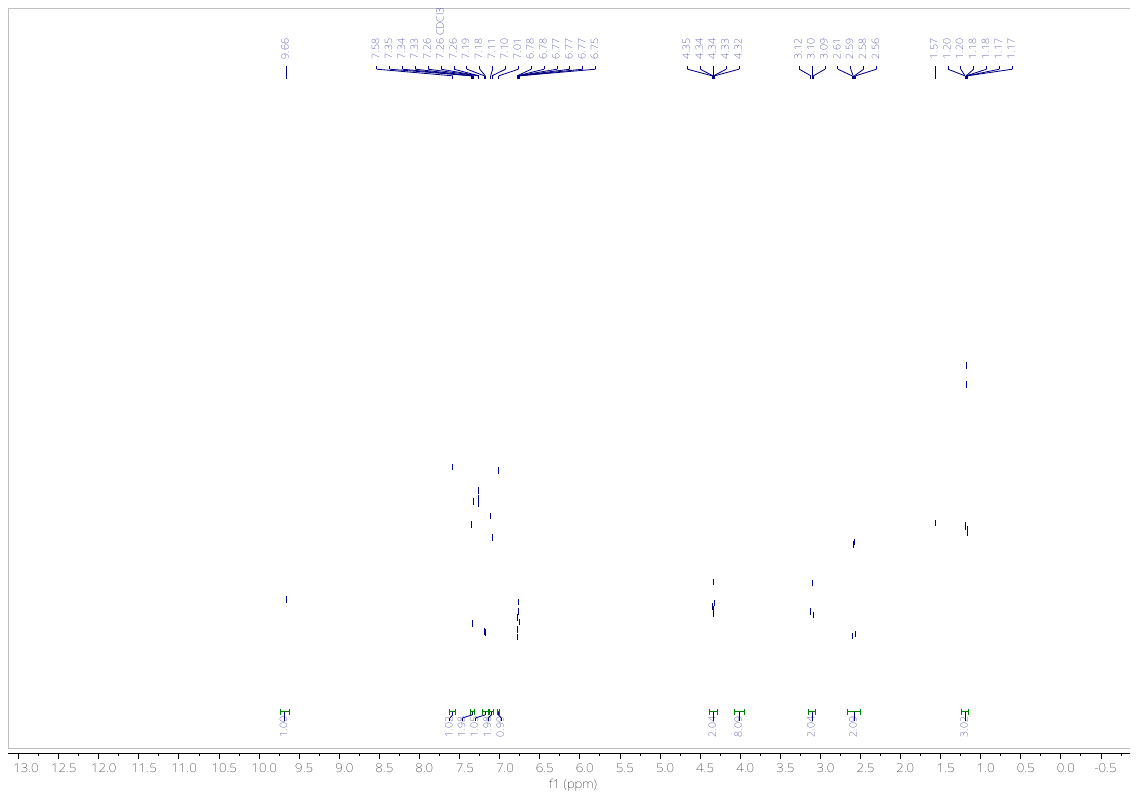

**Figure S5.** ^1^H NMR of **10c** in CDCl_3_.

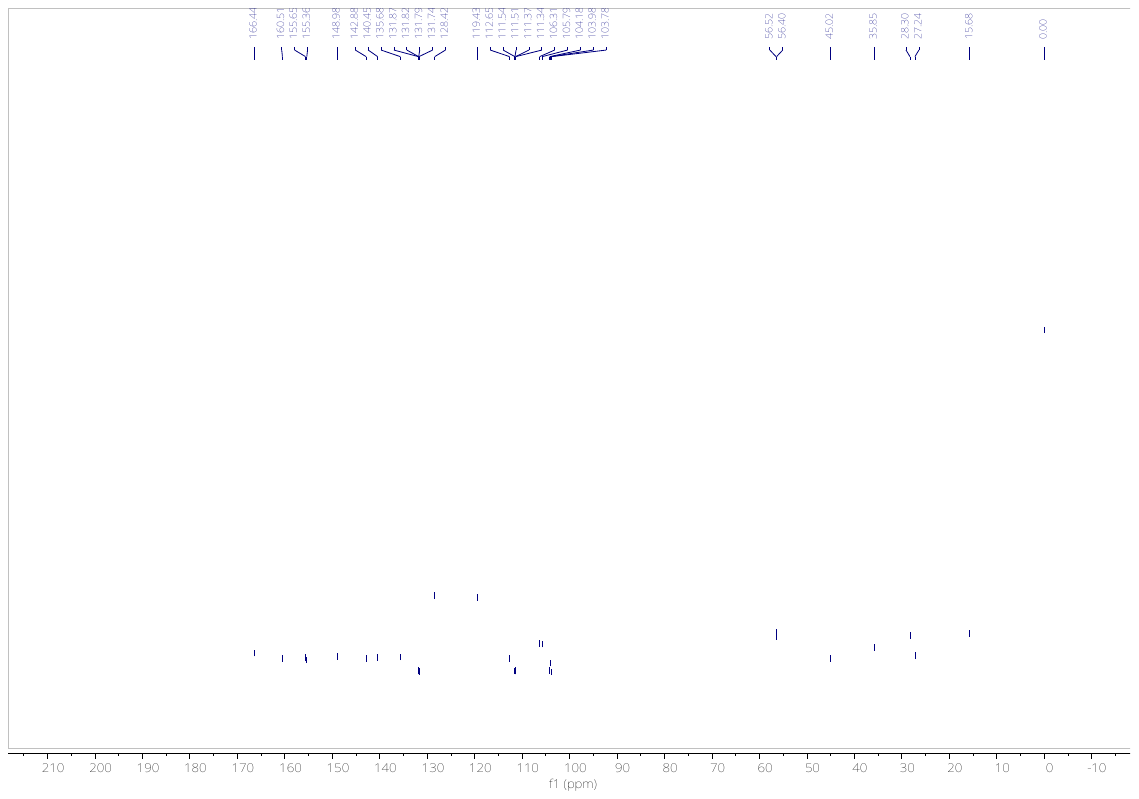

**Figure S6.** ^13^C NMR of **10c** in CDCl_3_.

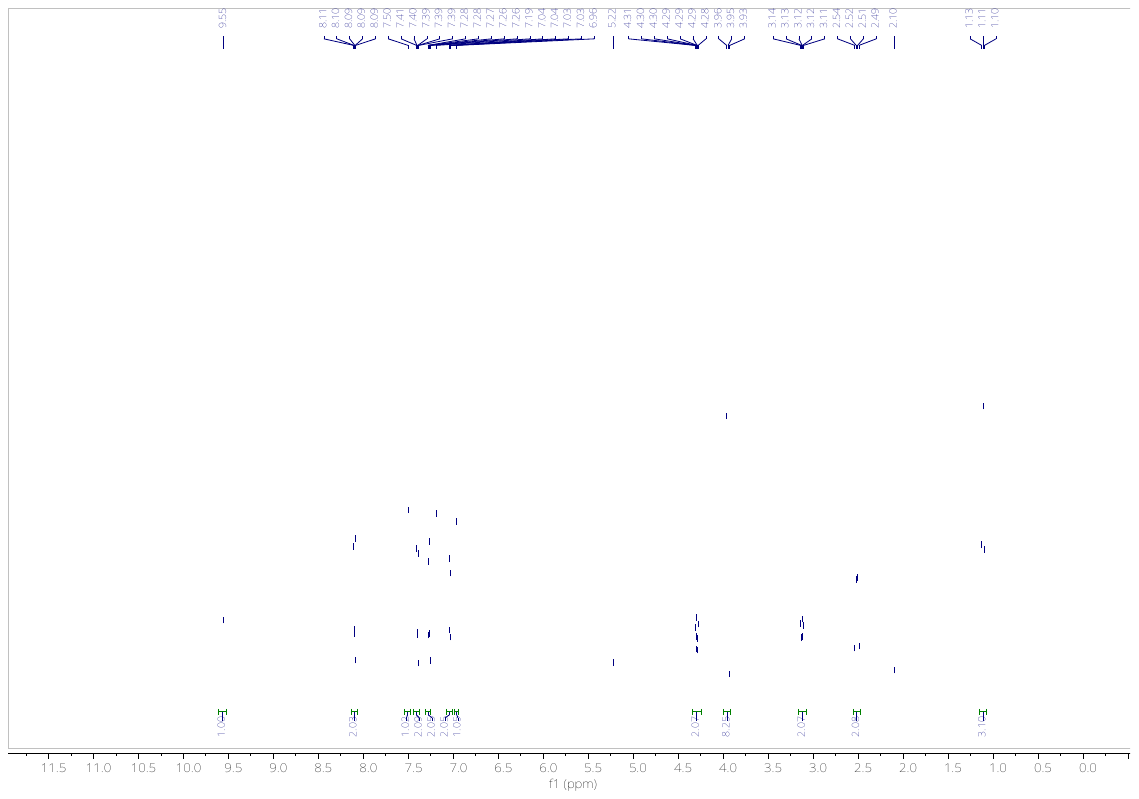

**Figure S7.** ^1^H NMR of **10d** in CDCl_3_.

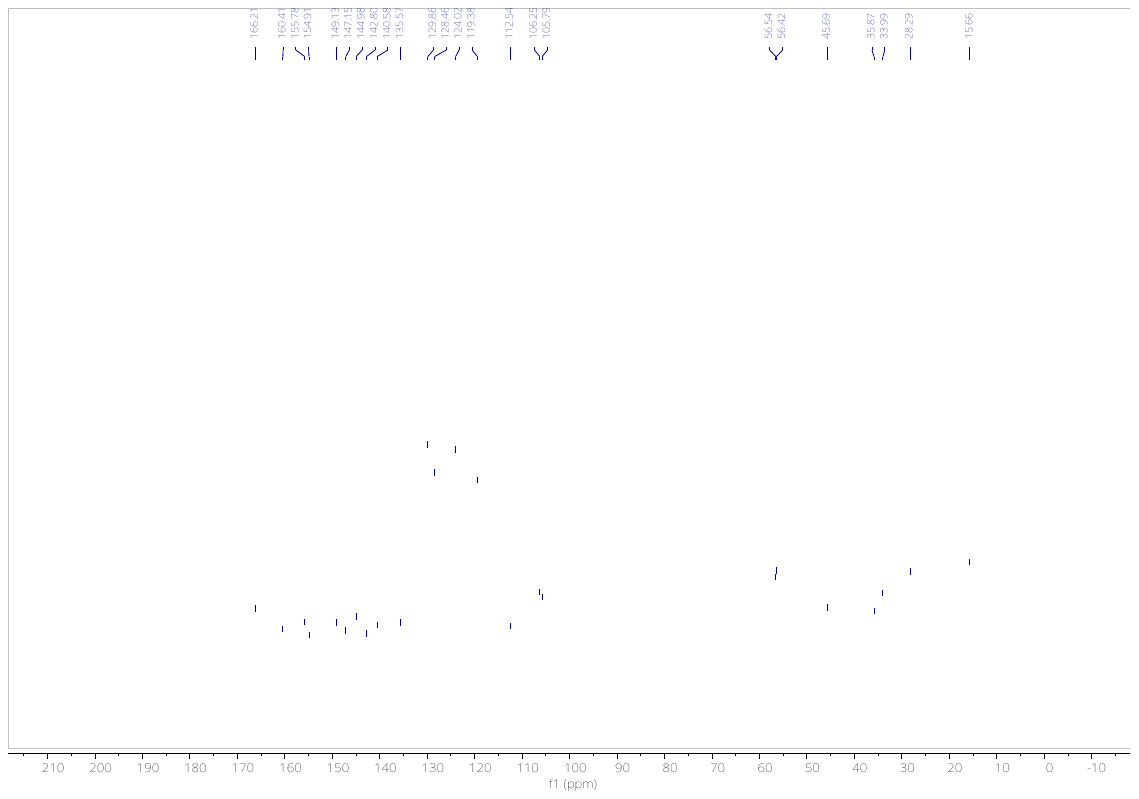

**Figure S8.** ^13^C NMR of **10d** in CDCl_3_.

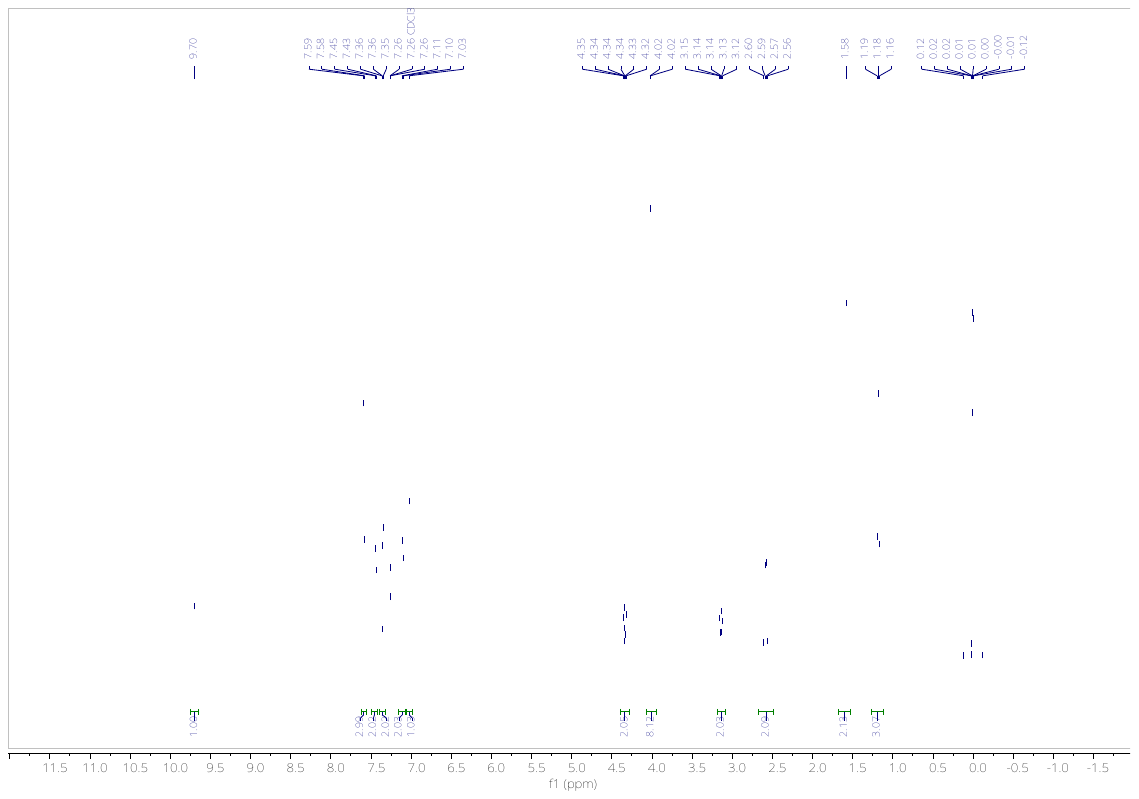

**Figure S9.** ^1^H NMR of **10e** in CDCl_3_.

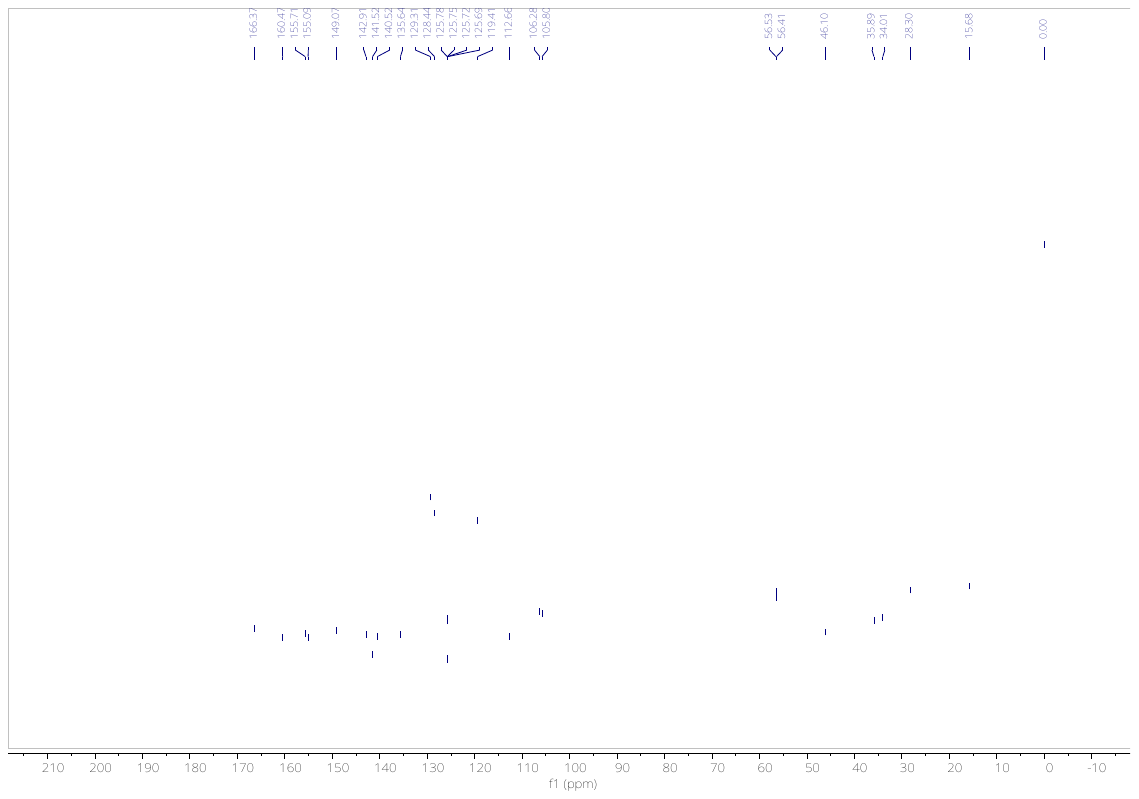

**Figure S10.** ^13^C NMR of **10e** in CDCl_3_.

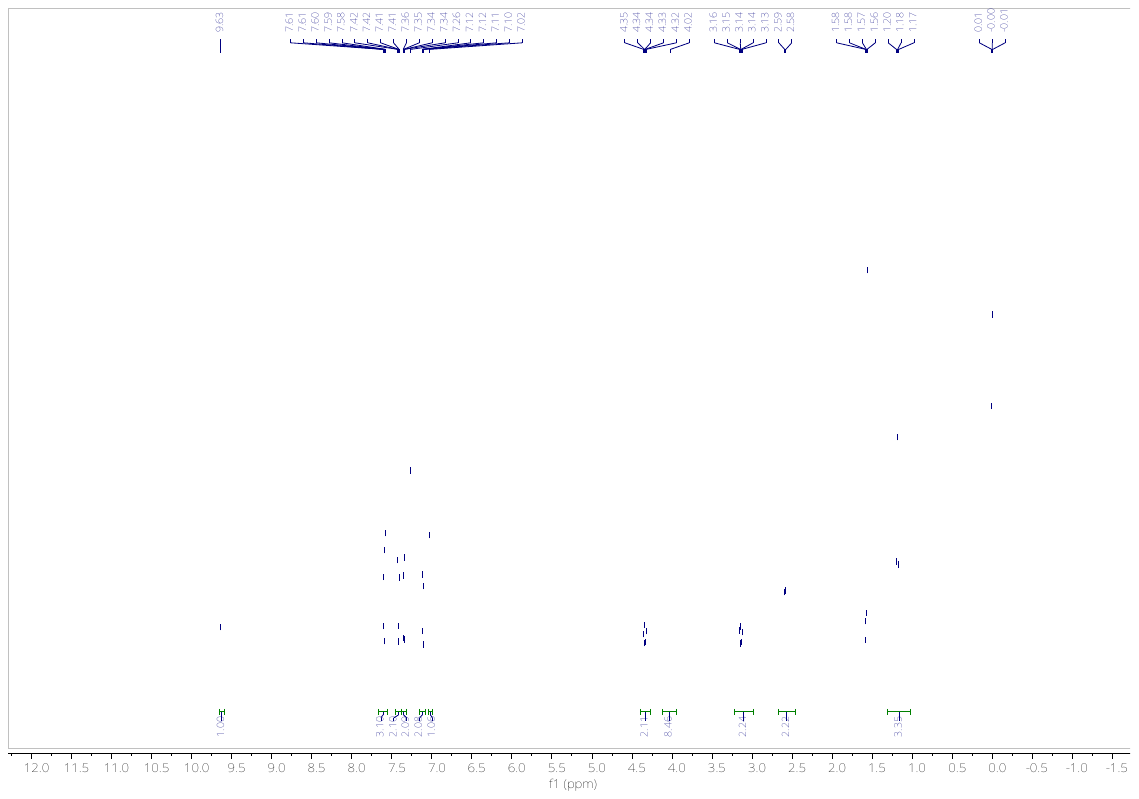

**Figure S11.** ^1^H NMR of **10f** in CDCl_3_.

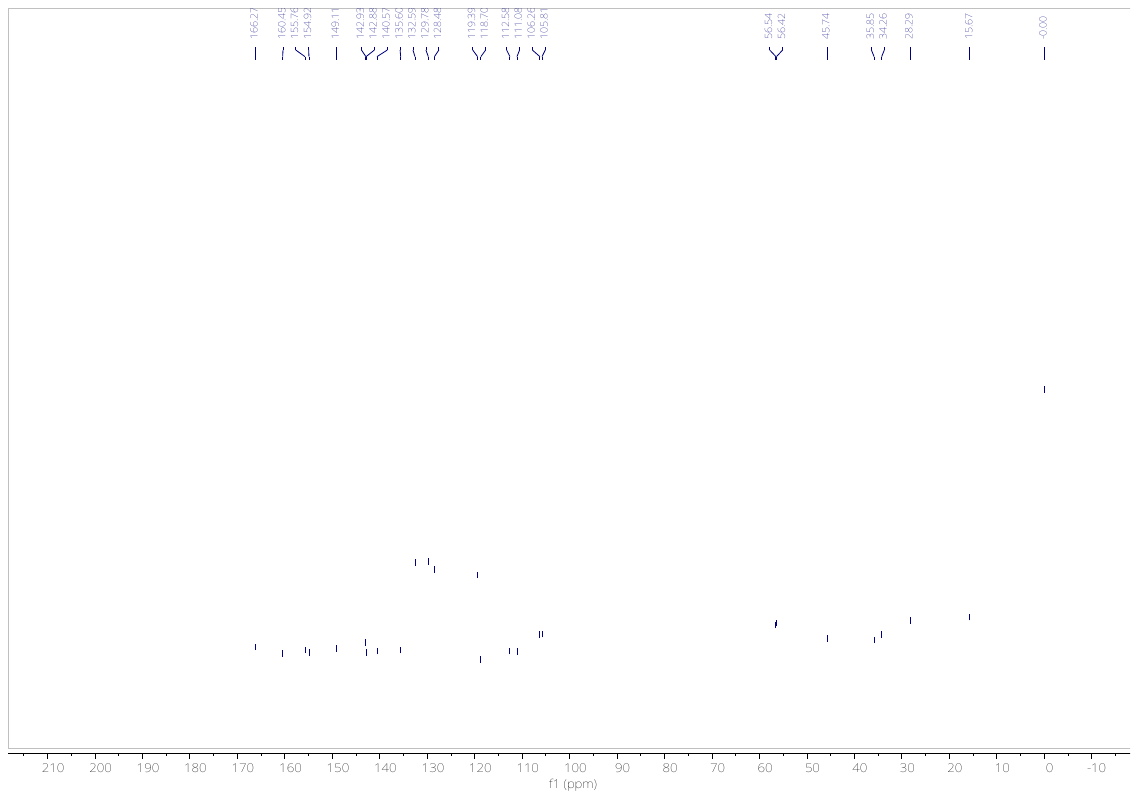

**Figure S12.** ^13^C NMR of **10f** in CDCl_3_.

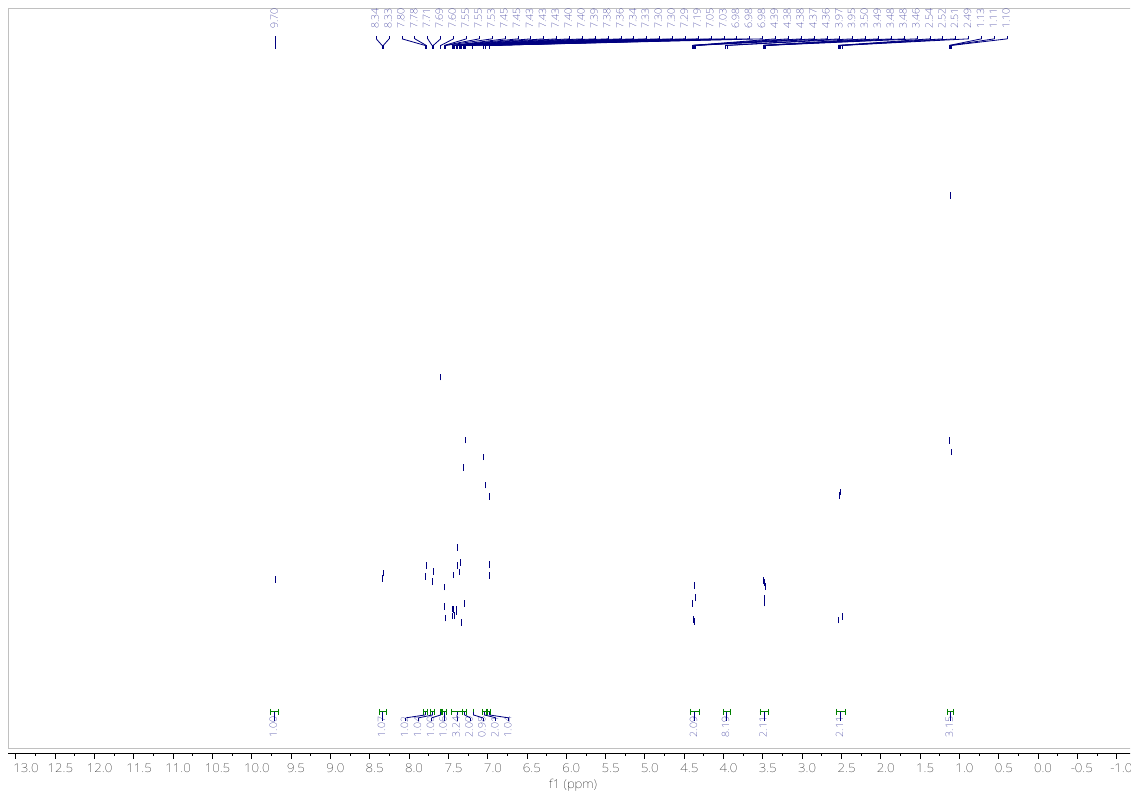

**Figure S13.** ^1^H NMR of **10g** in CDCl_3_.

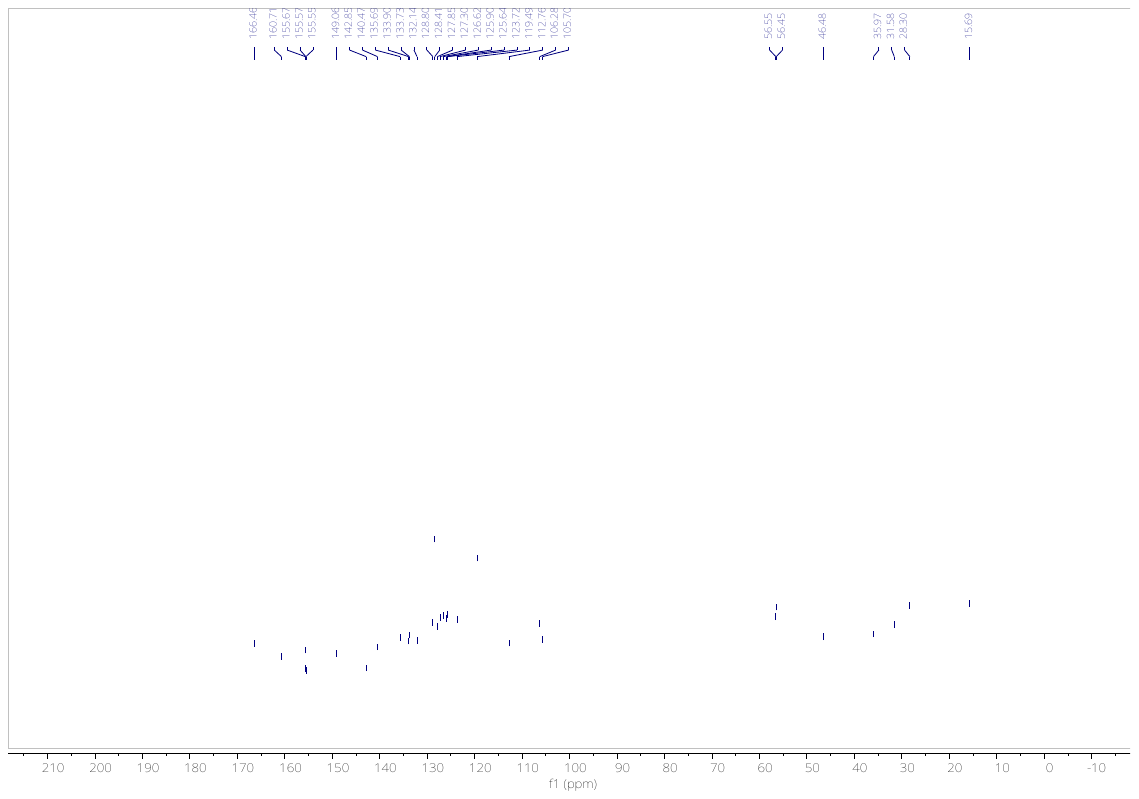
**Figure S14.** ^13^C NMR of **10g** in CDCl_3_.

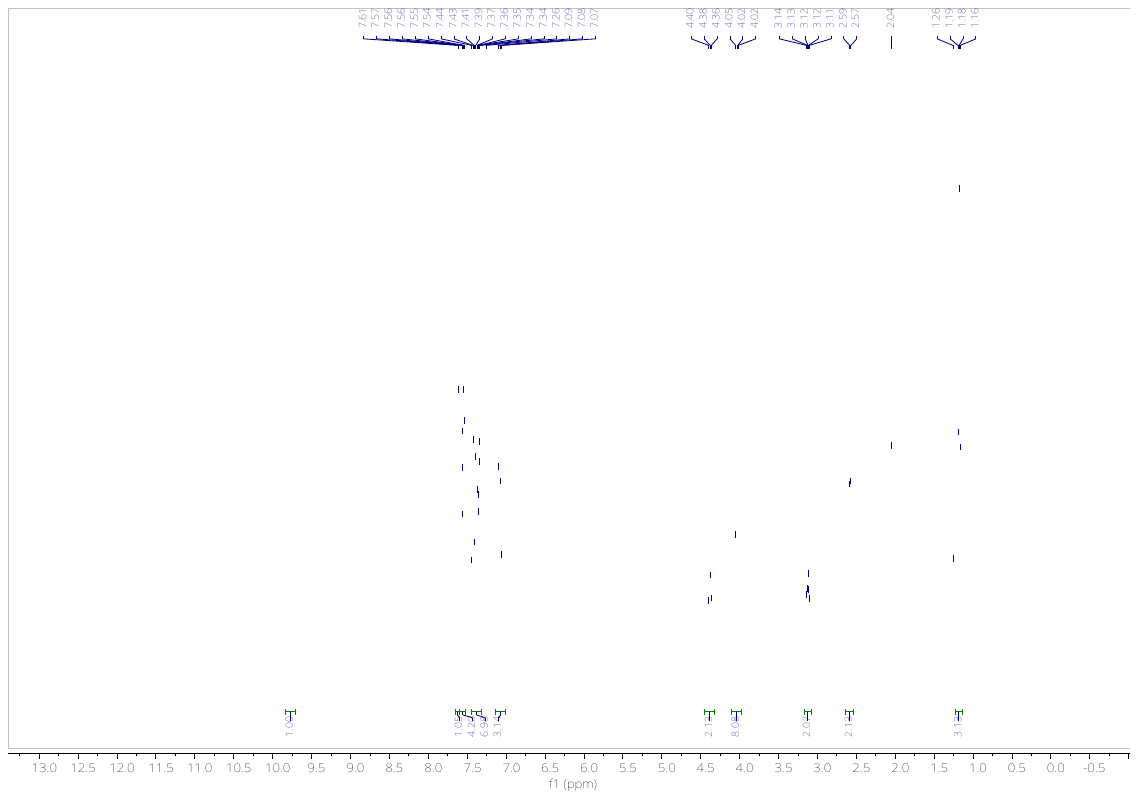

**Figure S15.** ^1^H NMR of **10h** in CDCl_3_.

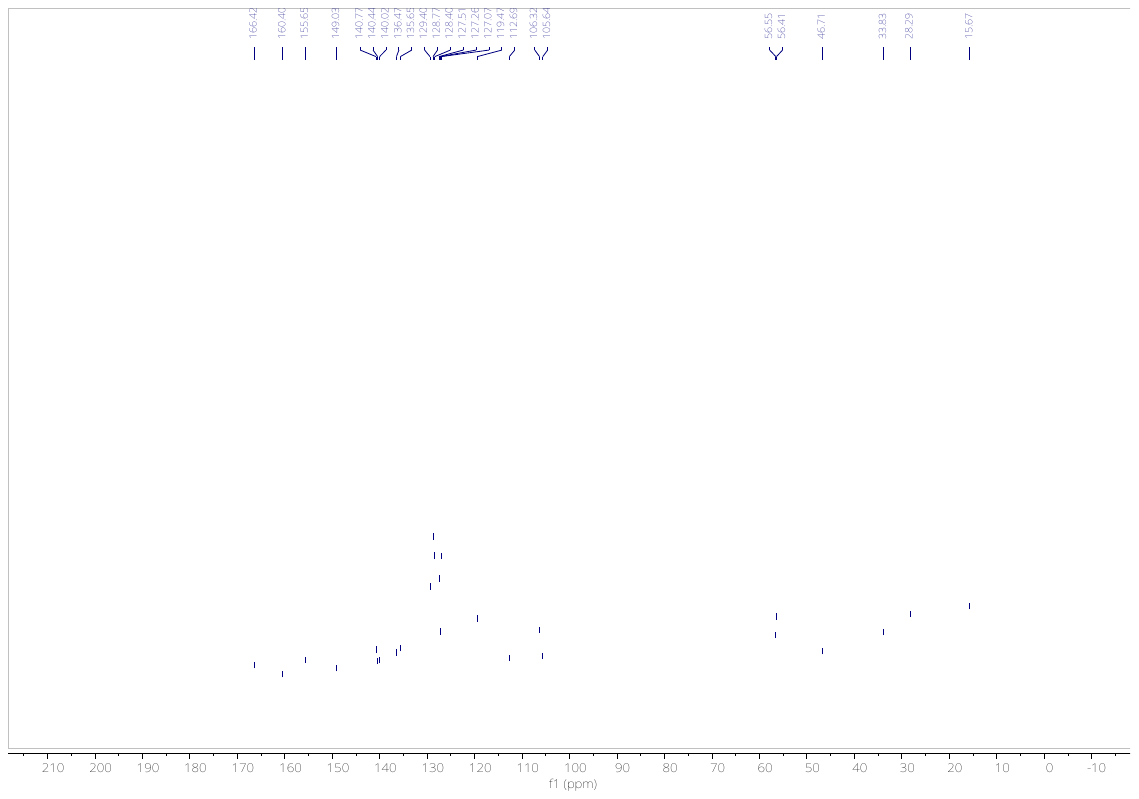

**Figure S16.** ^13^C NMR of **10h** in CDCl_3_.

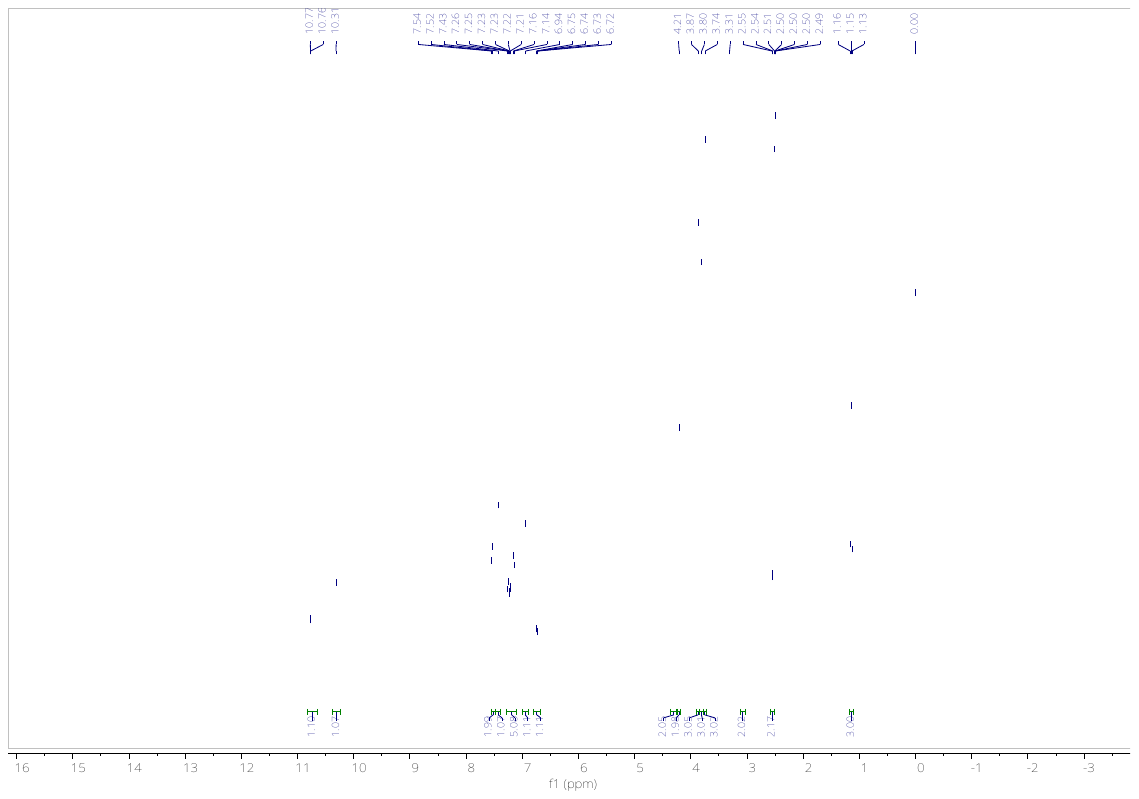

**Figure S17.** ^1^H NMR of **10i** in DMSO-*d*_6_.

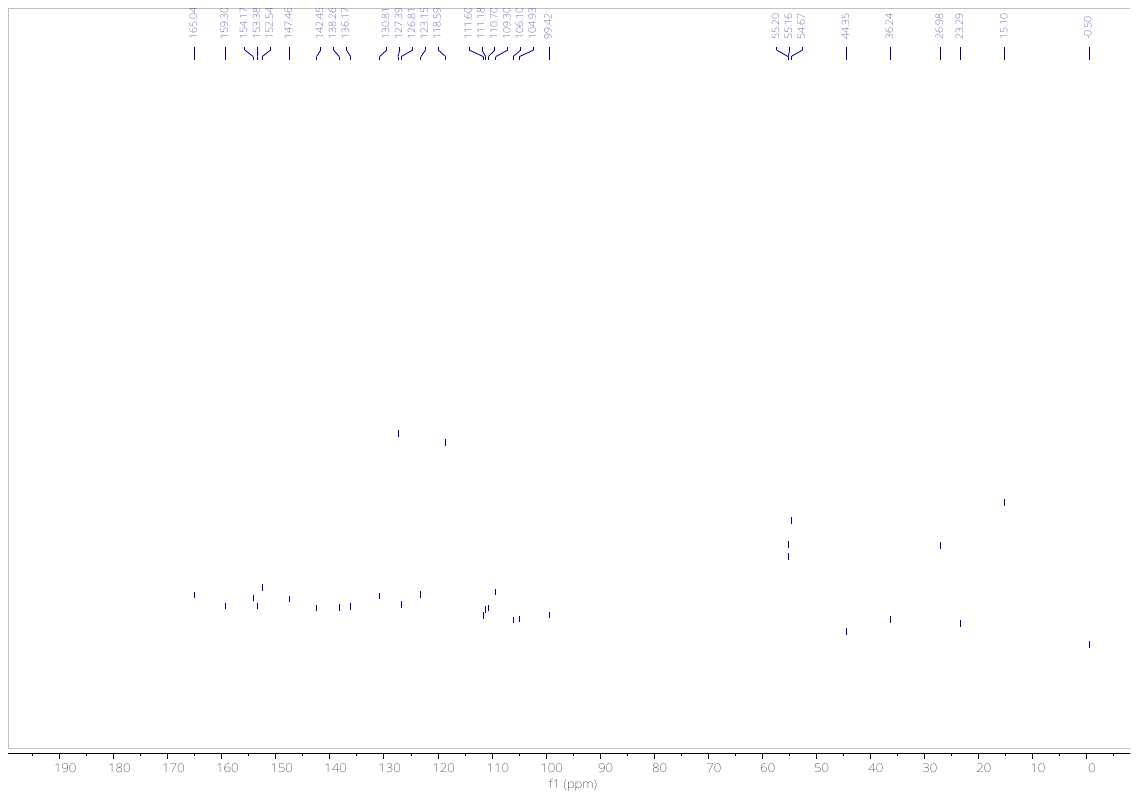

**Figure S18.** ^13^C NMR of **10i** in DMSO-*d*_6_.

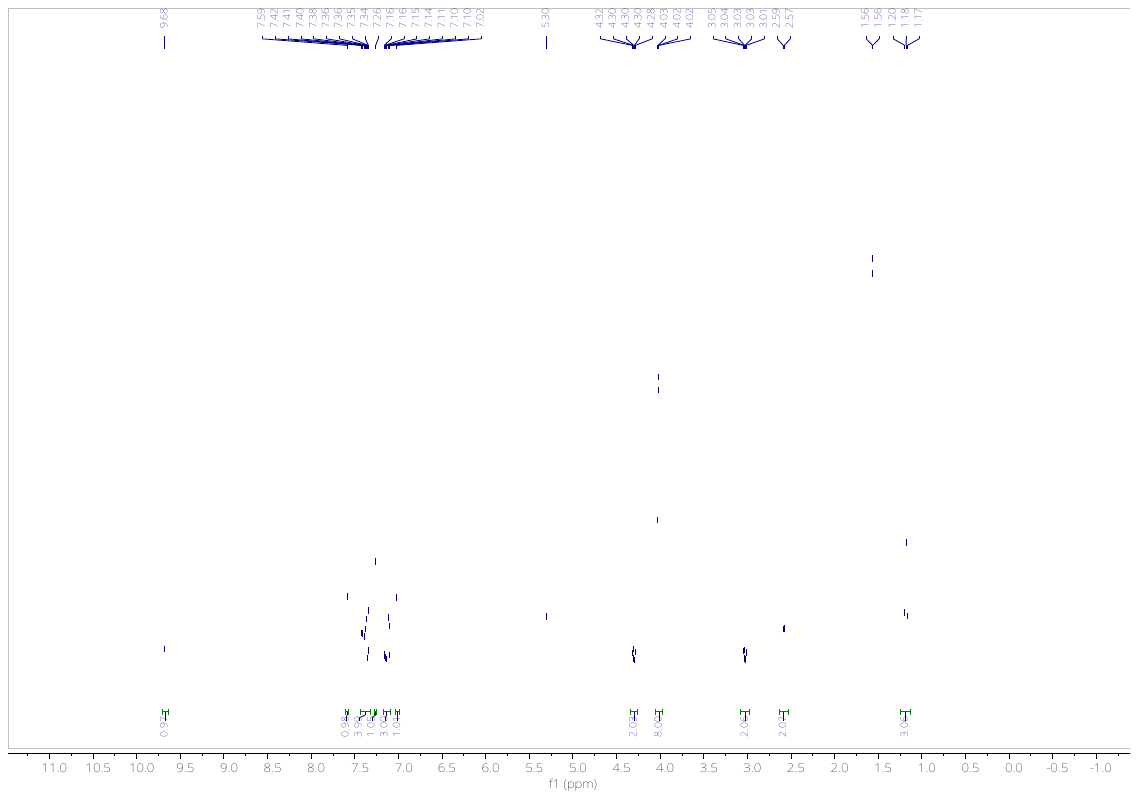

**Figure S19.** ^1^H NMR of **10j** in CDCl_3_.

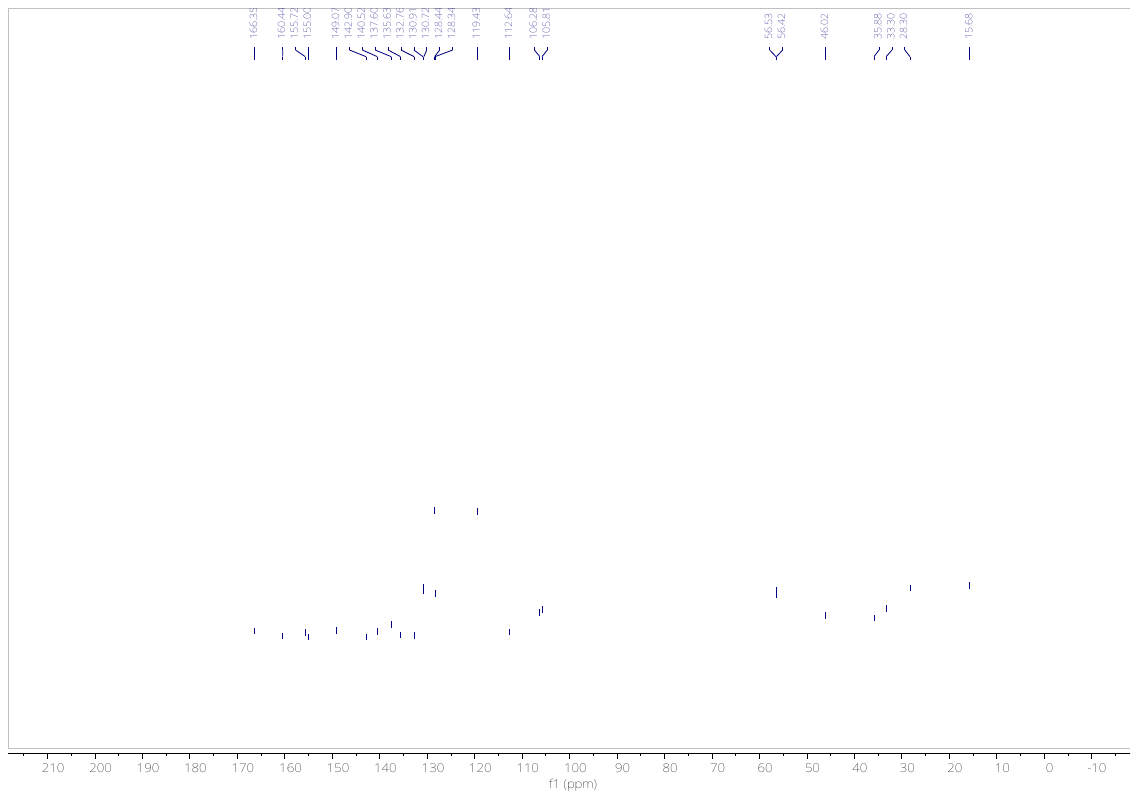
**Figure S20.** ^13^C NMR of **10j** in CDCl_3_.

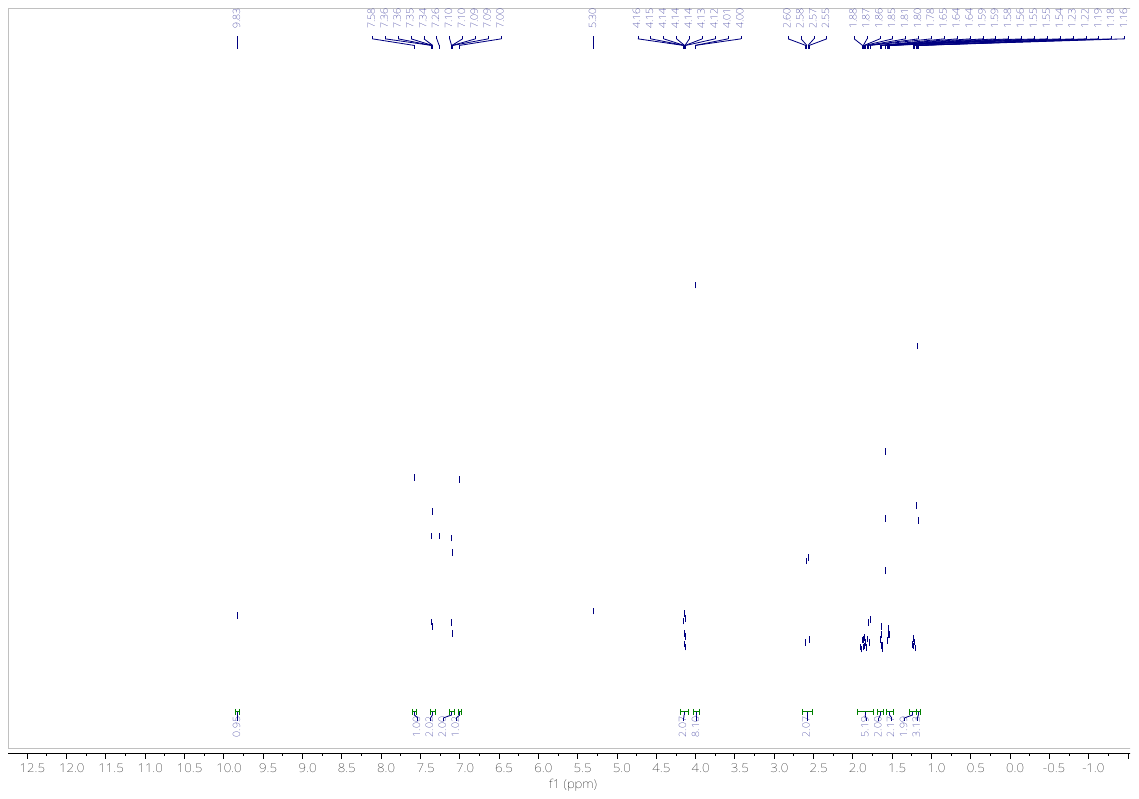

**Figure S21.** ^1^H NMR of **10k** in CDCl_3_.

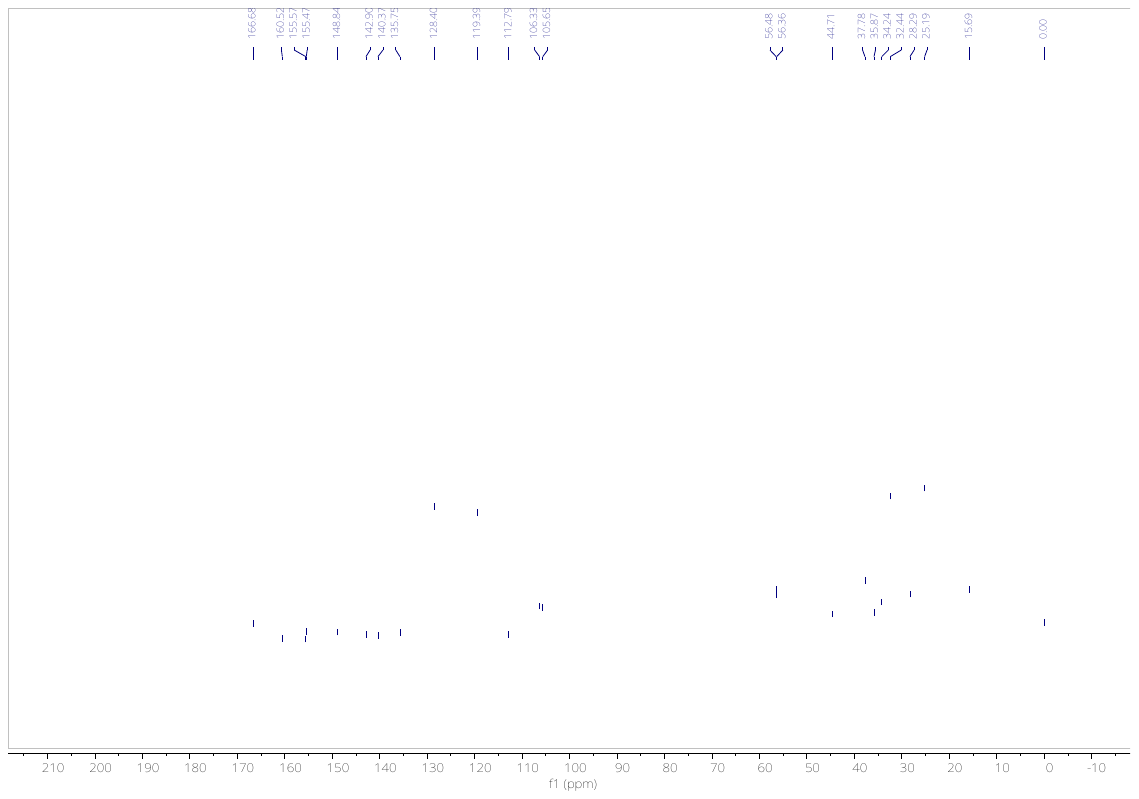

**Figure S22.** ^13^C NMR of **10k** in CDCl_3_.

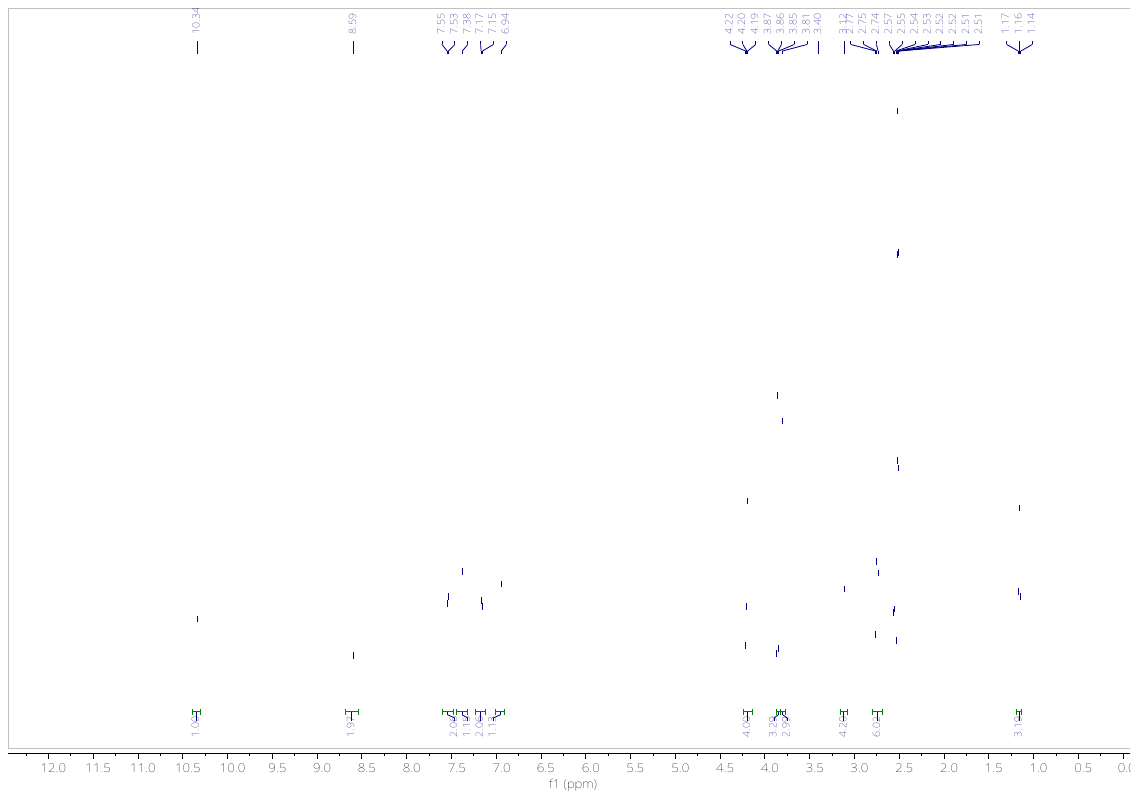

**Figure S23.** ^1^H NMR of **10m** in DMSO-*d*_6_.

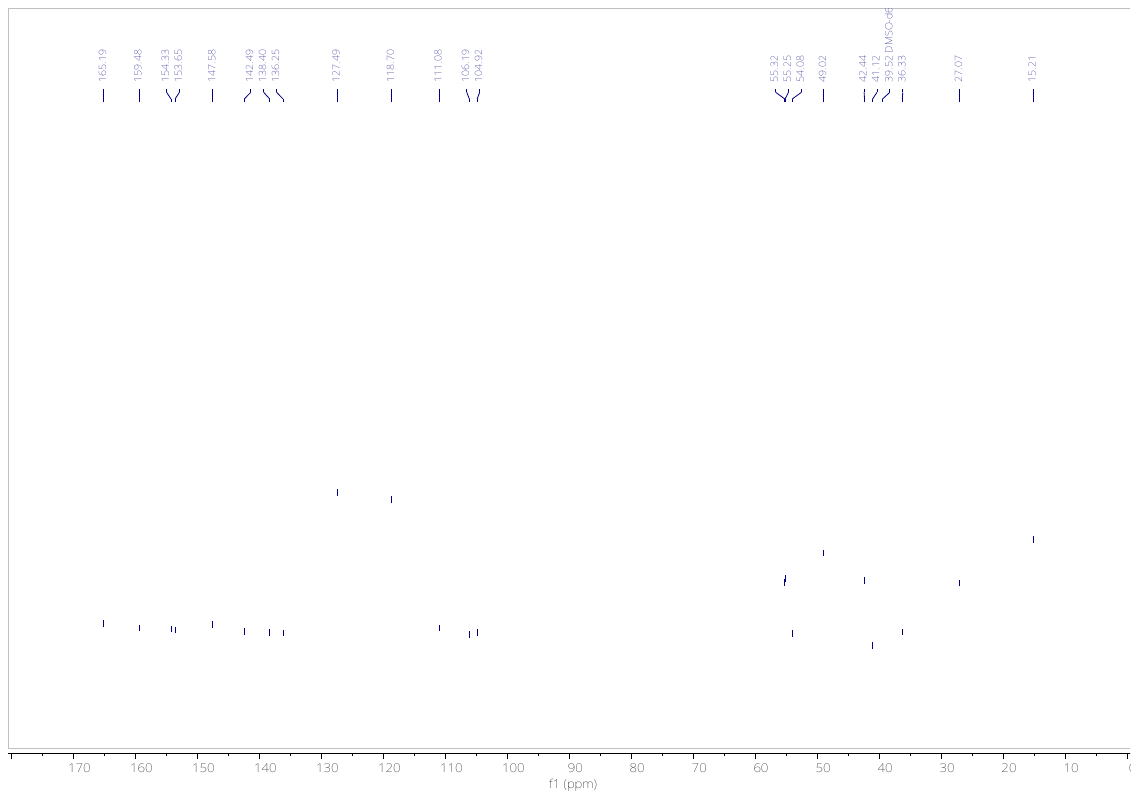

**Figure S24.** ^1^C NMR of **10m** in DMSO-*d*_6_.

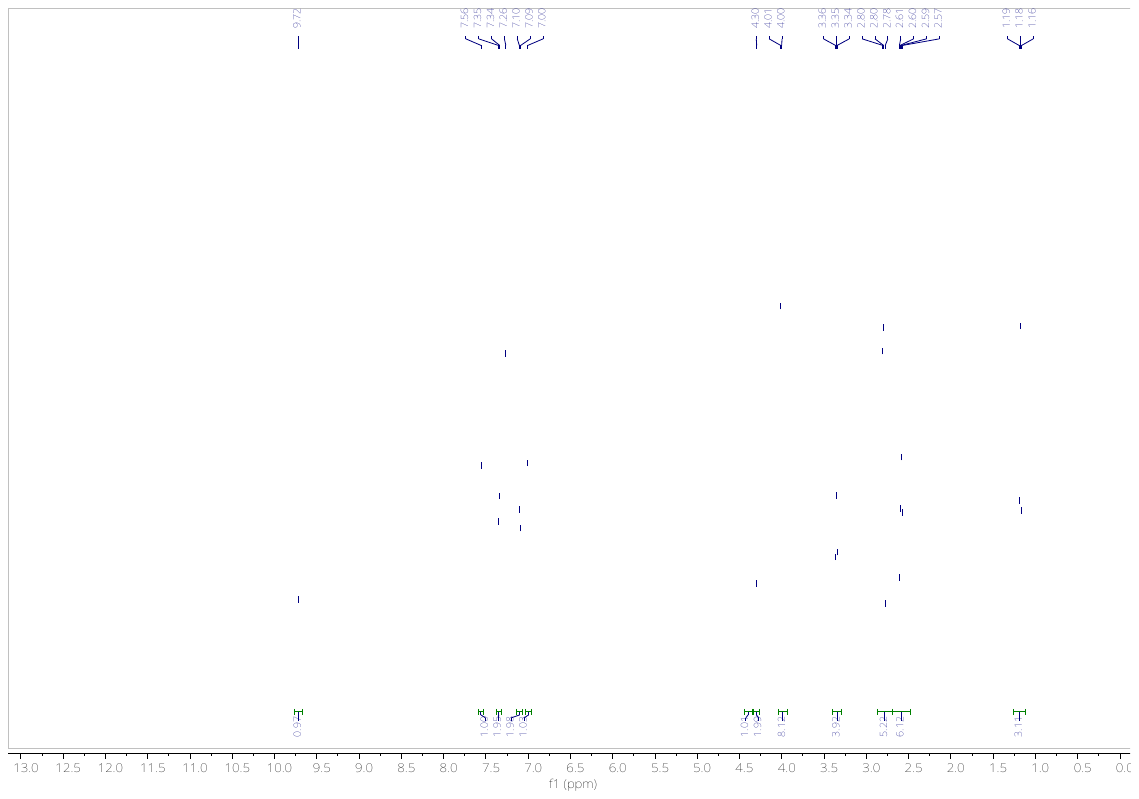
**Figure S25.** ^1^H NMR of **10n** in CDCl_3_.
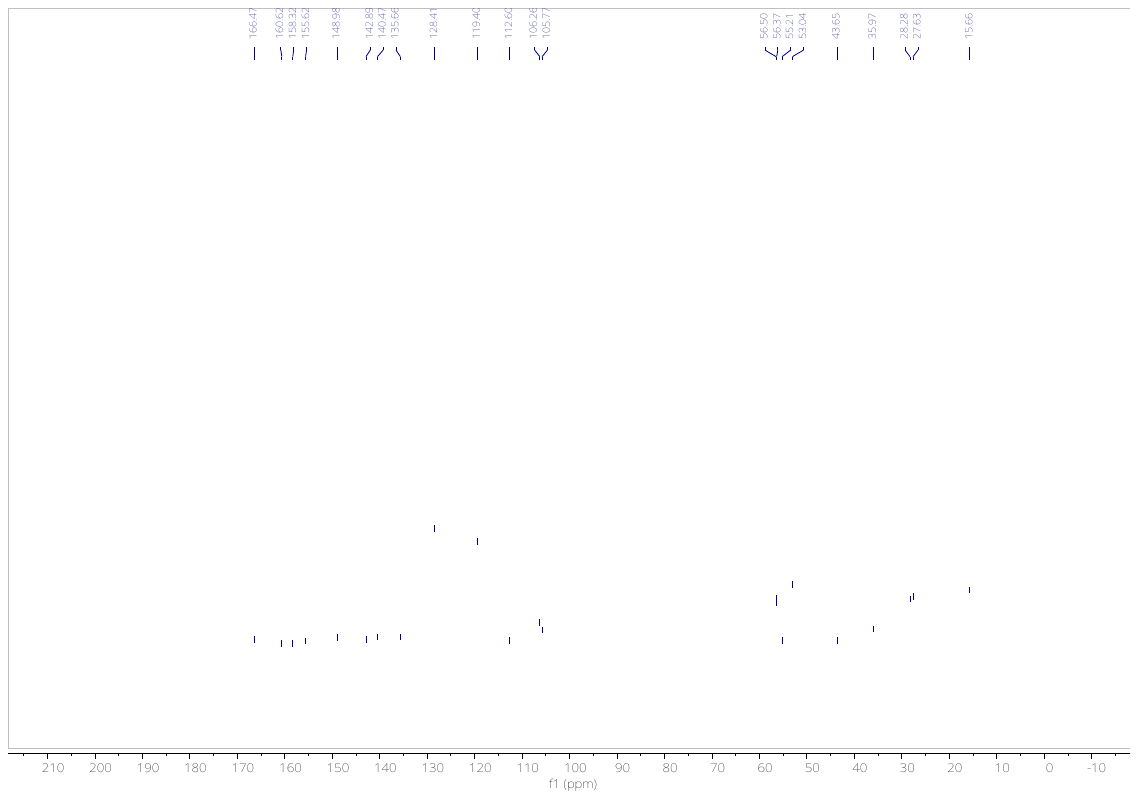

**Figure S26.** ^13^C NMR of **10n** in CDCl_3_.

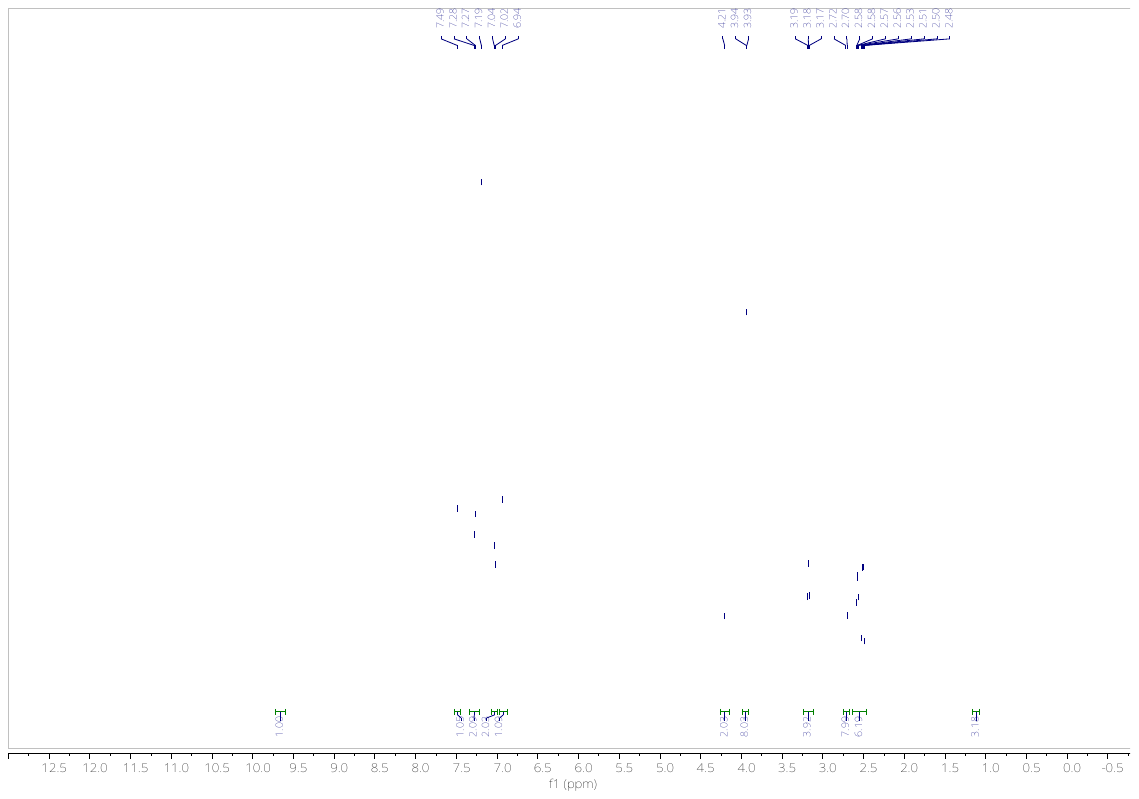

**Figure S27.** ^1^H NMR of **10o** in CDCl_3_.

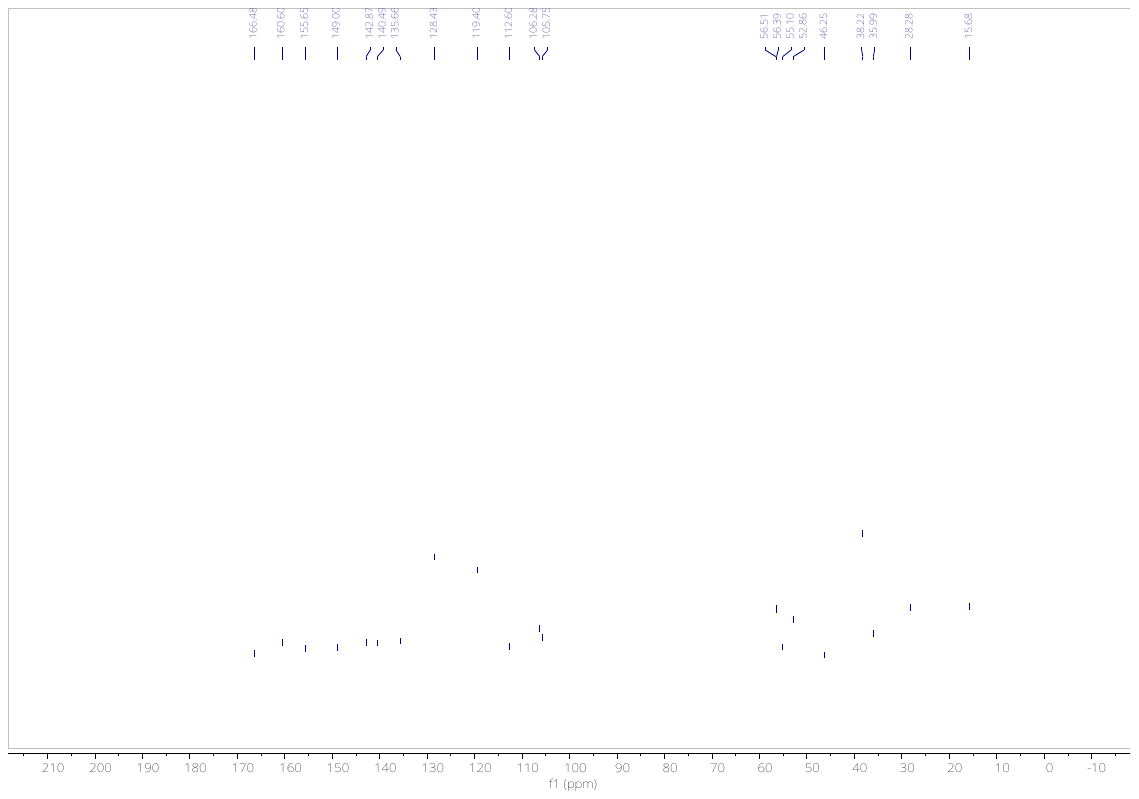

**Figure S28.** ^13^C NMR of **10o** in CDCl_3_.

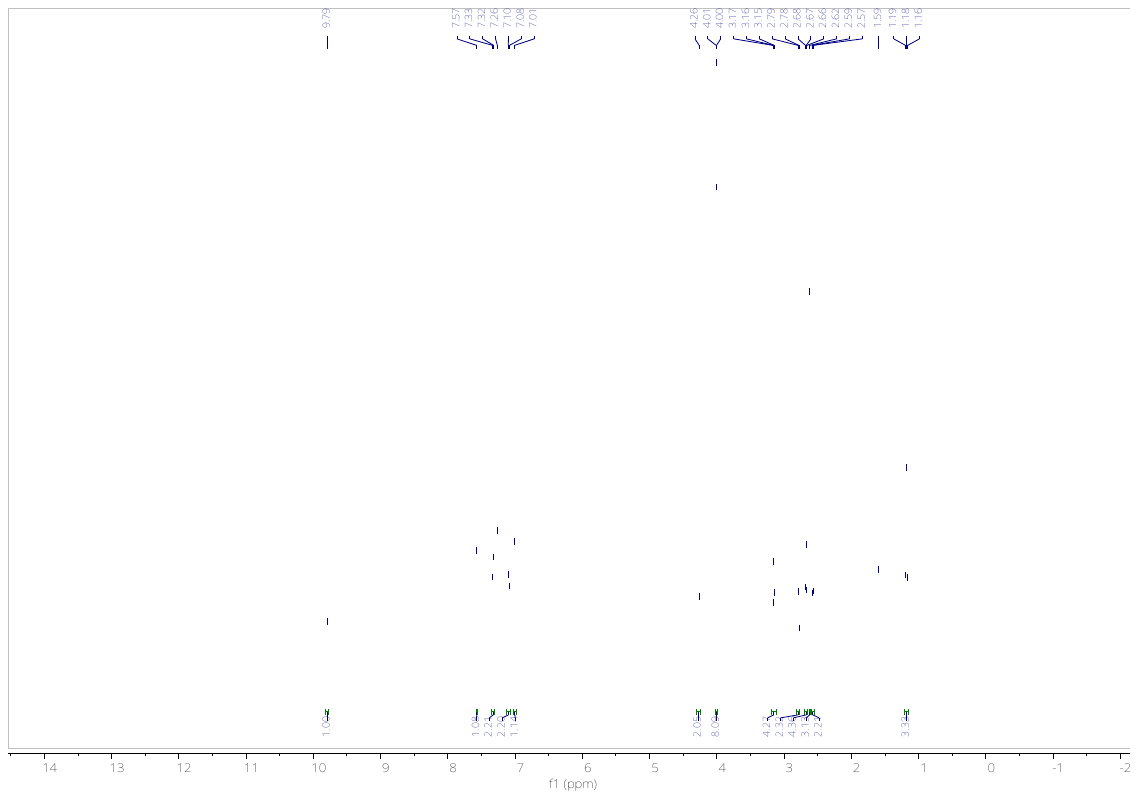

**Figure S29.** ^1^H NMR of **10p** in CDCl_3_.

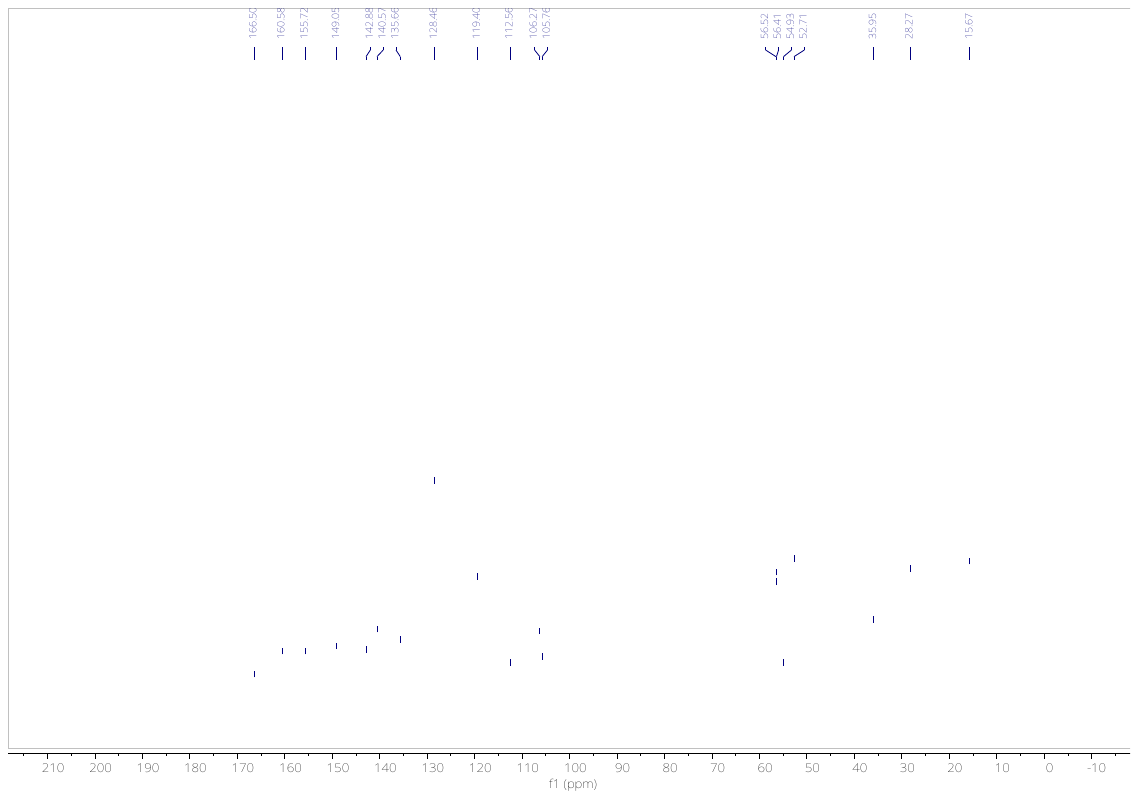

**Figure S30.** ^1^H ^13^C NMR of **10p** in CDCl_3_.

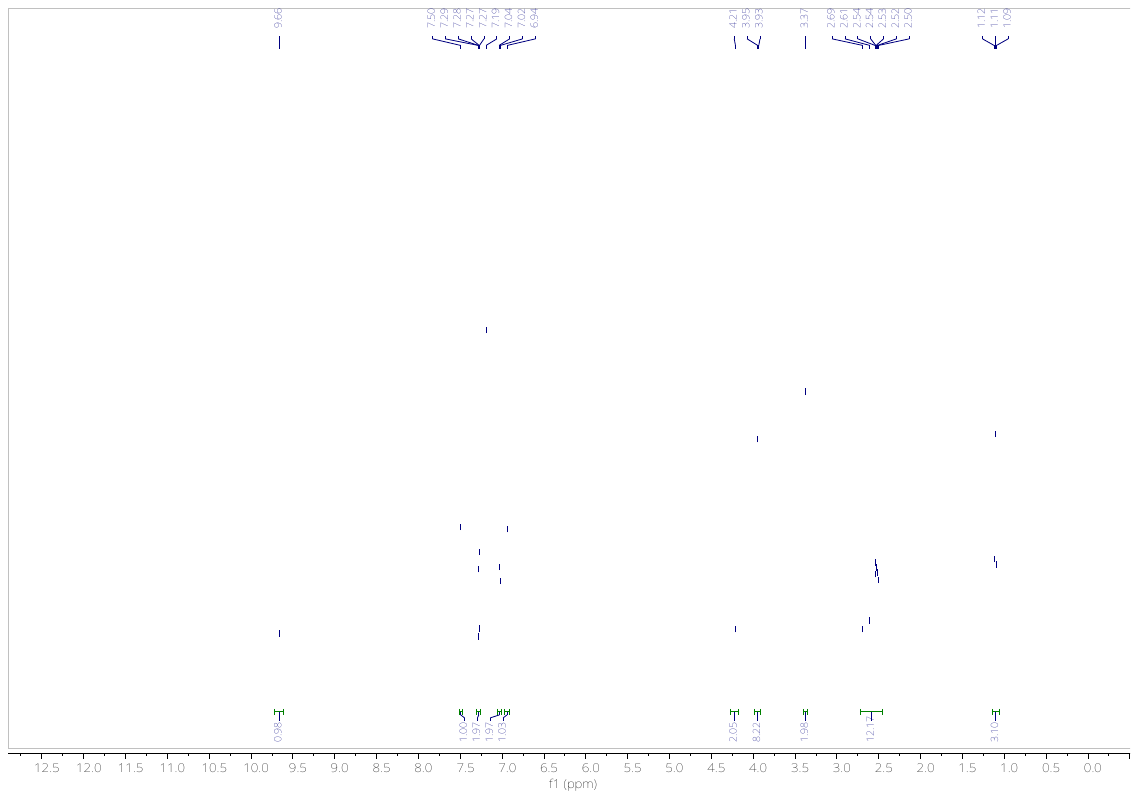

**Figure S31.** ^1^H NMR of **10q** in CDCl_3_.

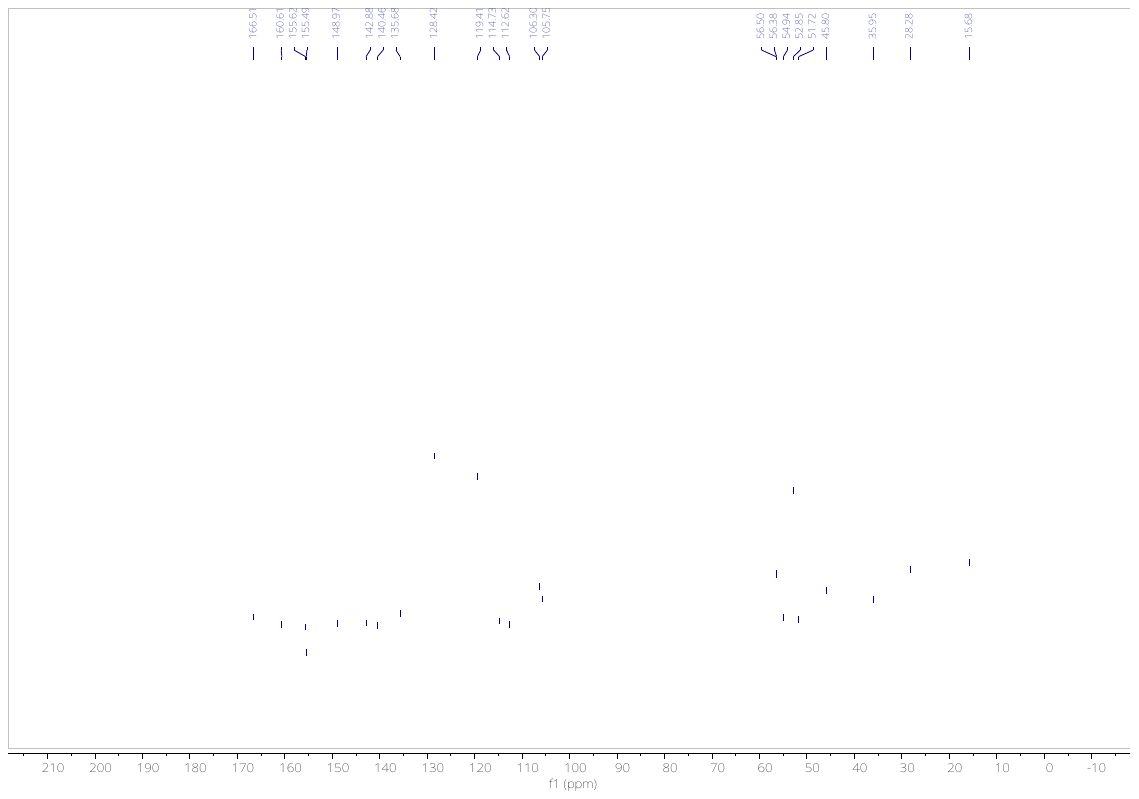

**Figure S32.** ^13^C NMR of **10q** in CDCl_3_.

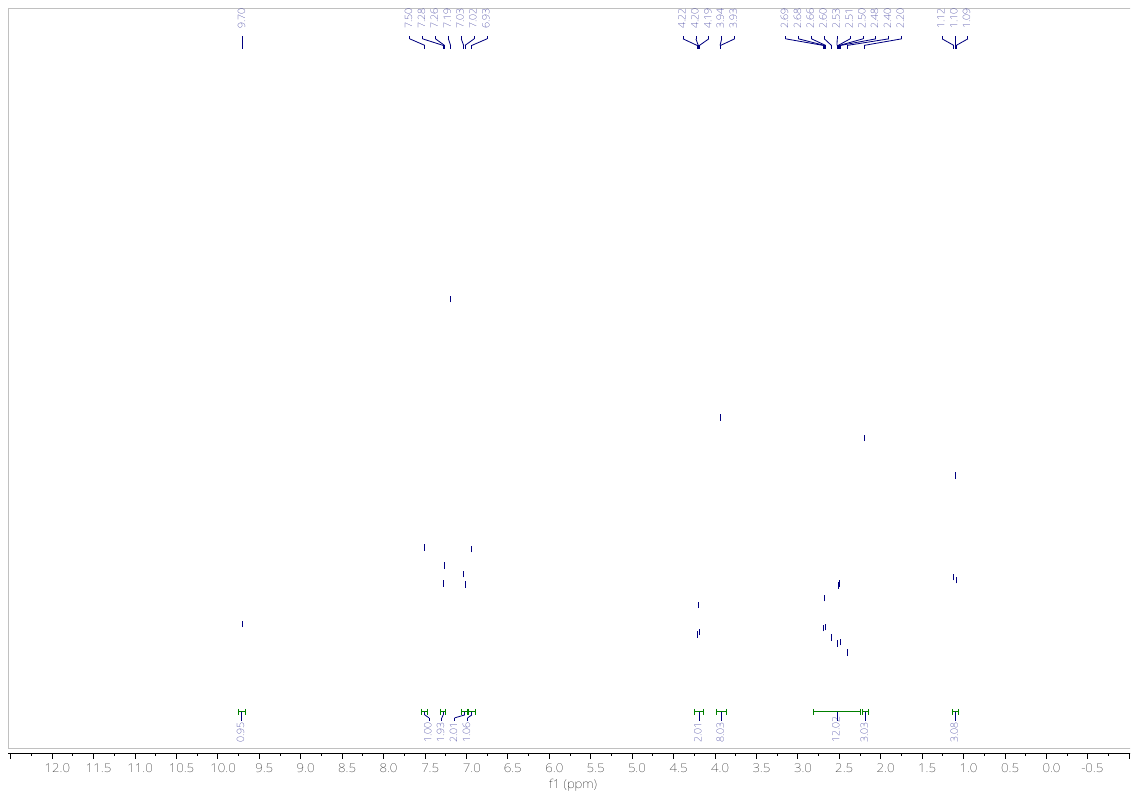

**Figure S33.** ^1^H NMR of **10r** in CDCl_3_.

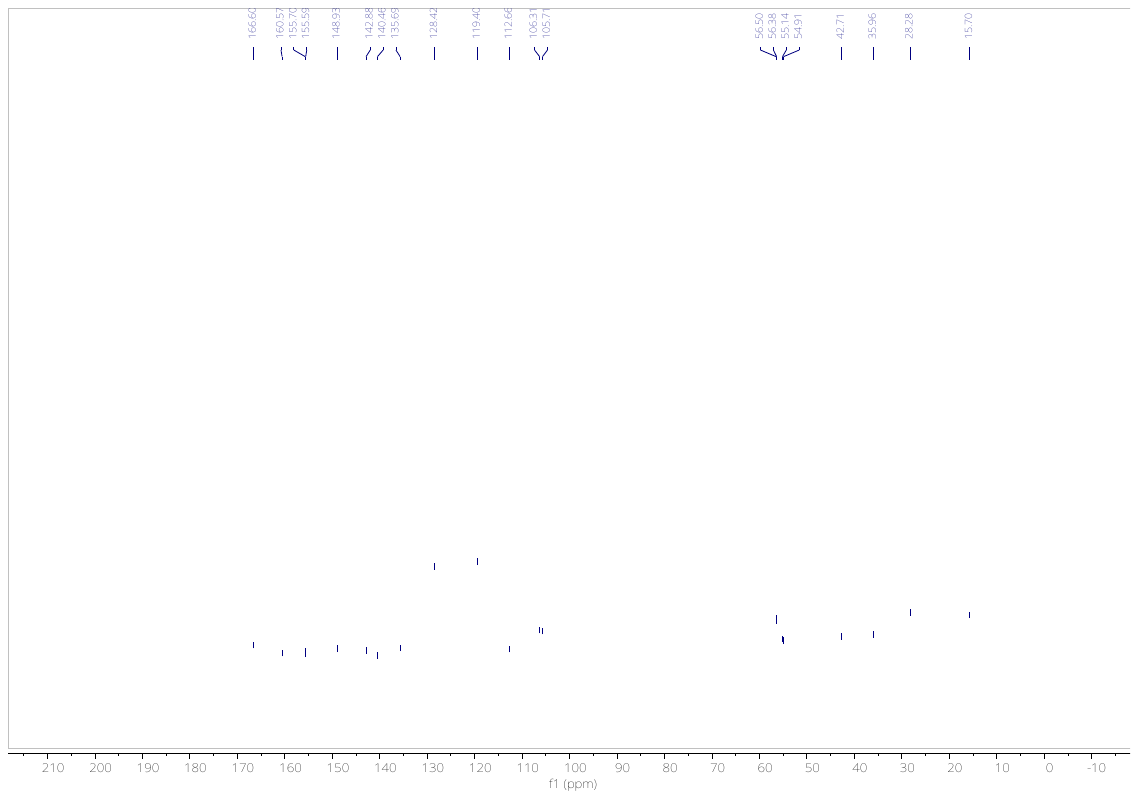

**Figure S34.** ^13^C NMR of **10r** in CDCl_3_.

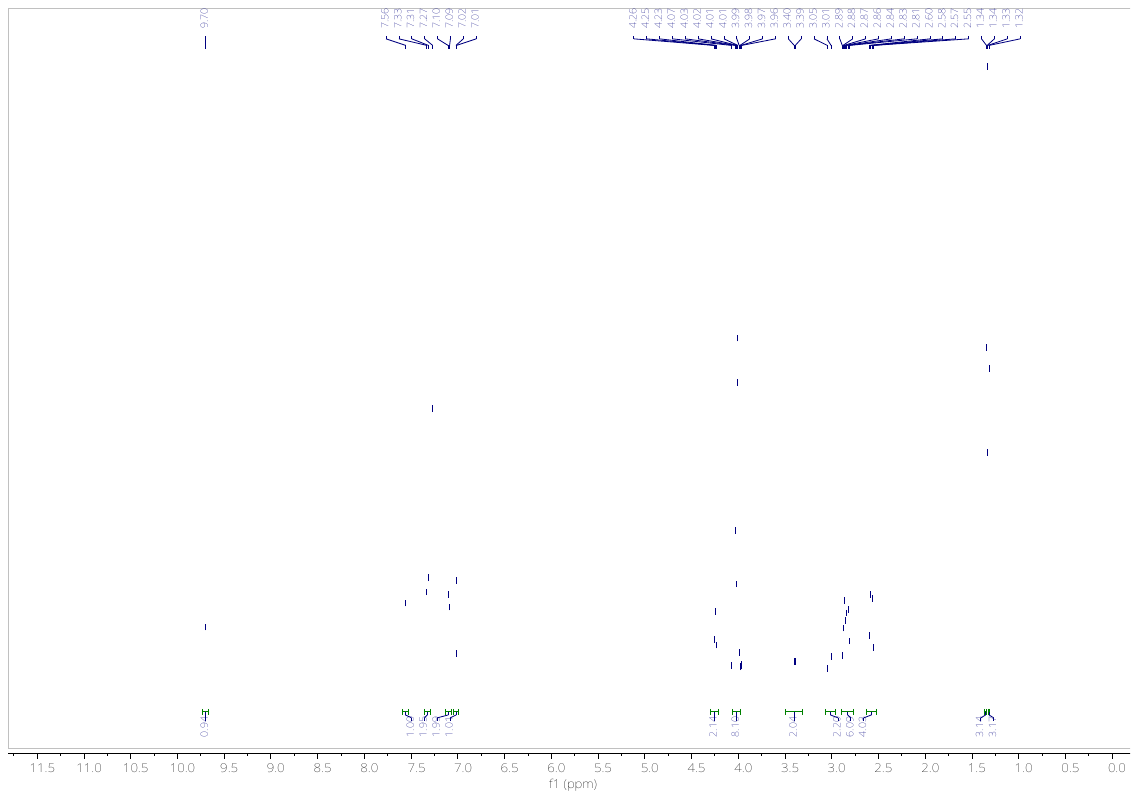

**Figure S35.** ^1^H NMR of **10s** in CDCl_3_.

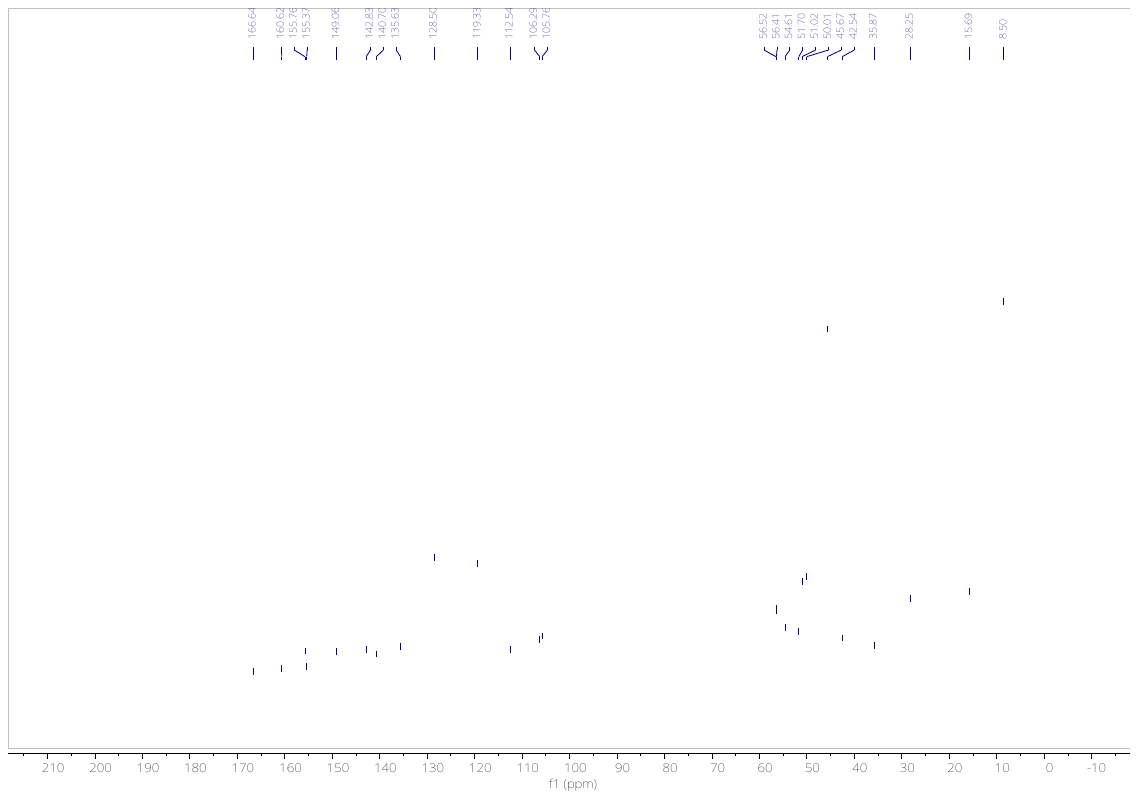

**Figure S36.** ^13^C NMR of **10s** in CDCl_3_.

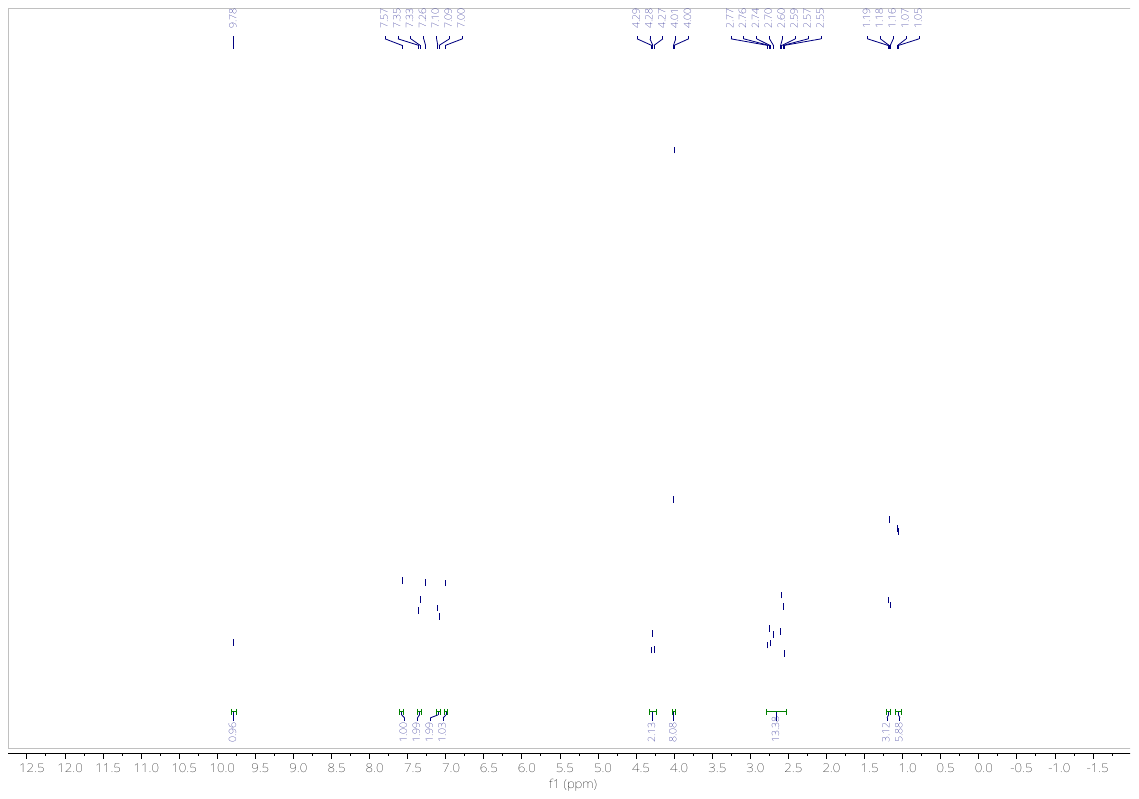

**Figure S37.** ^1^H NMR of **10t** in CDCl_3_.

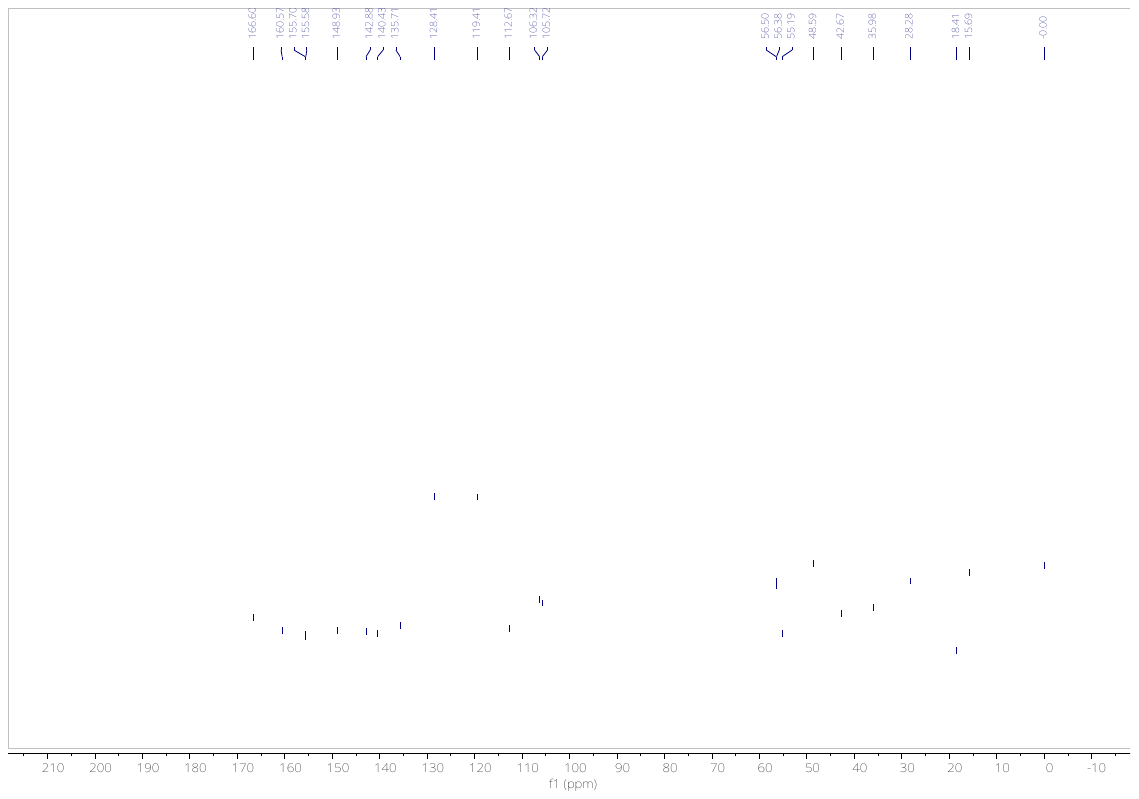
**Figure S38.** ^13^C NMR of **10t** in CDCl_3_.

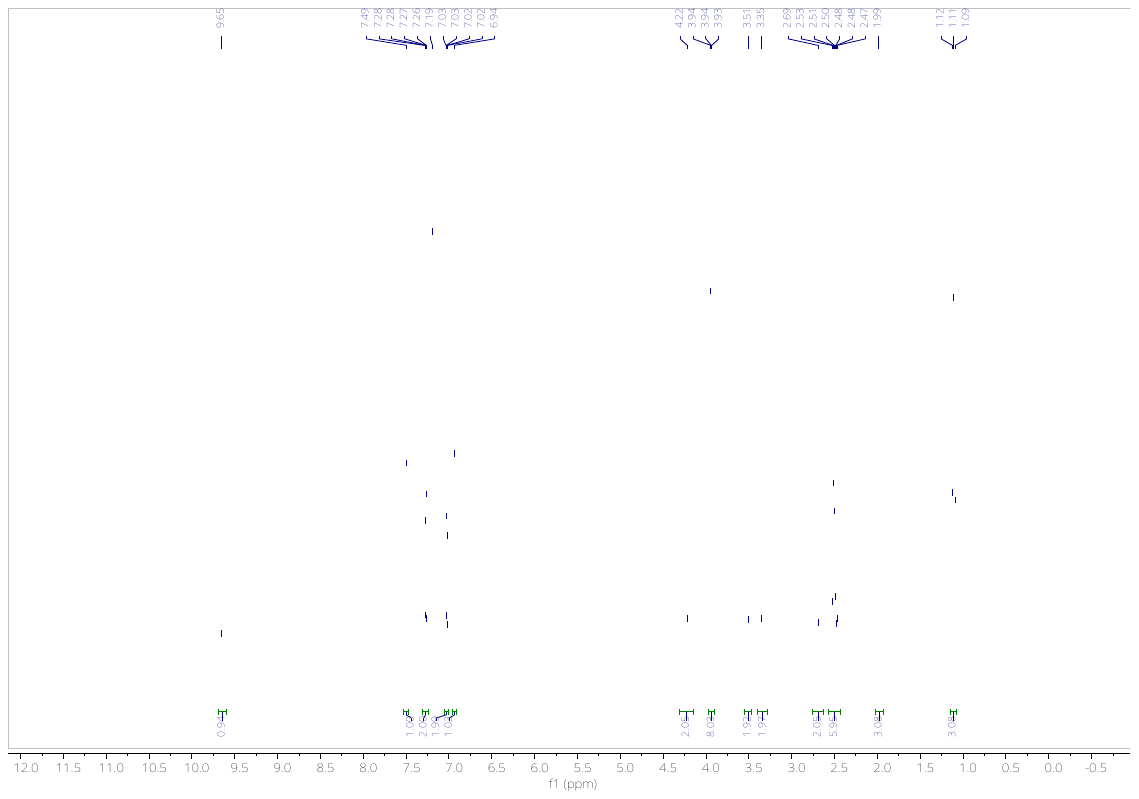

**Figure S39.** ^1^H NMR of **10u** in CDCl_3_.

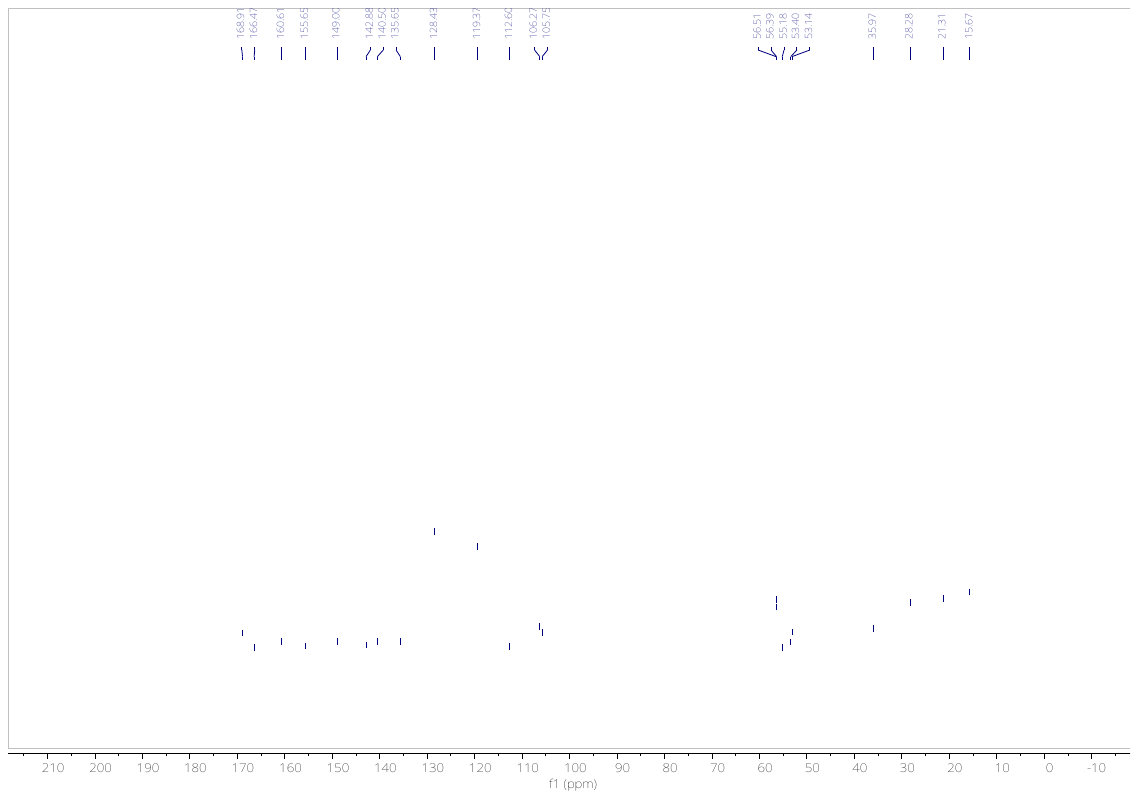

**Figure S40.** ^13^C NMR of **10u** in CDCl_3_.

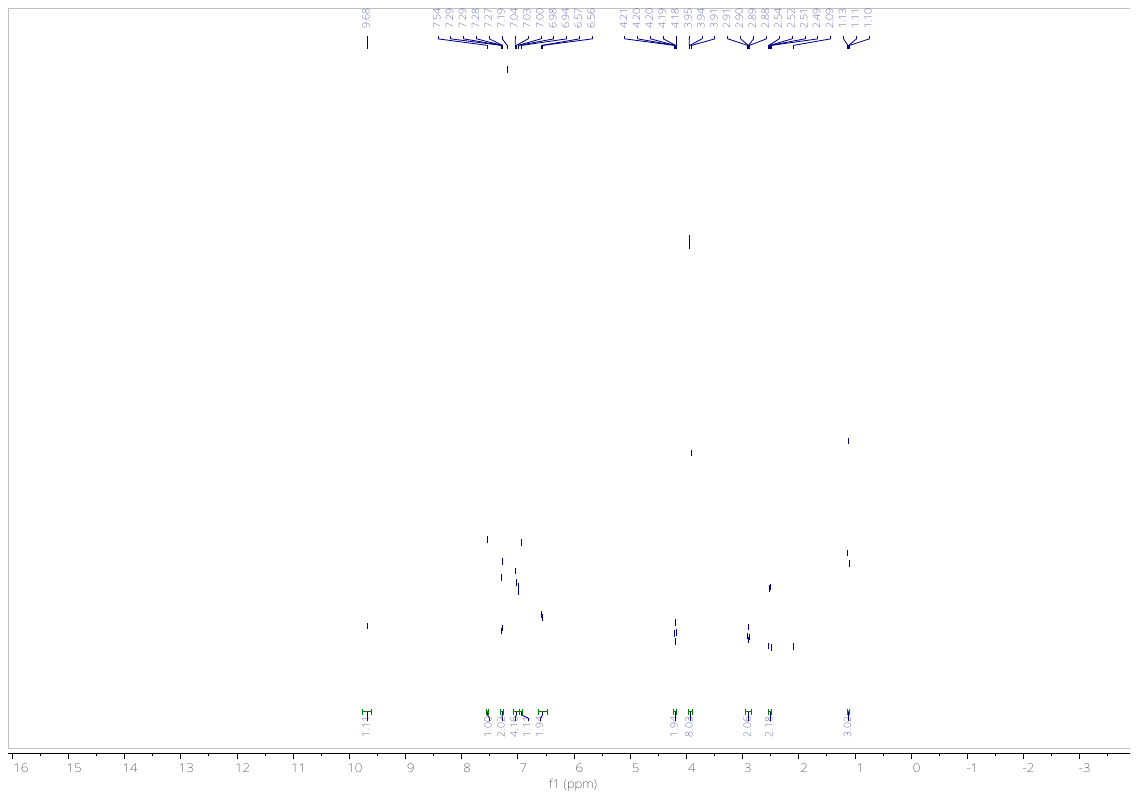

**Figure S41.** ^1^H NMR of **10v** in CDCl_3_.
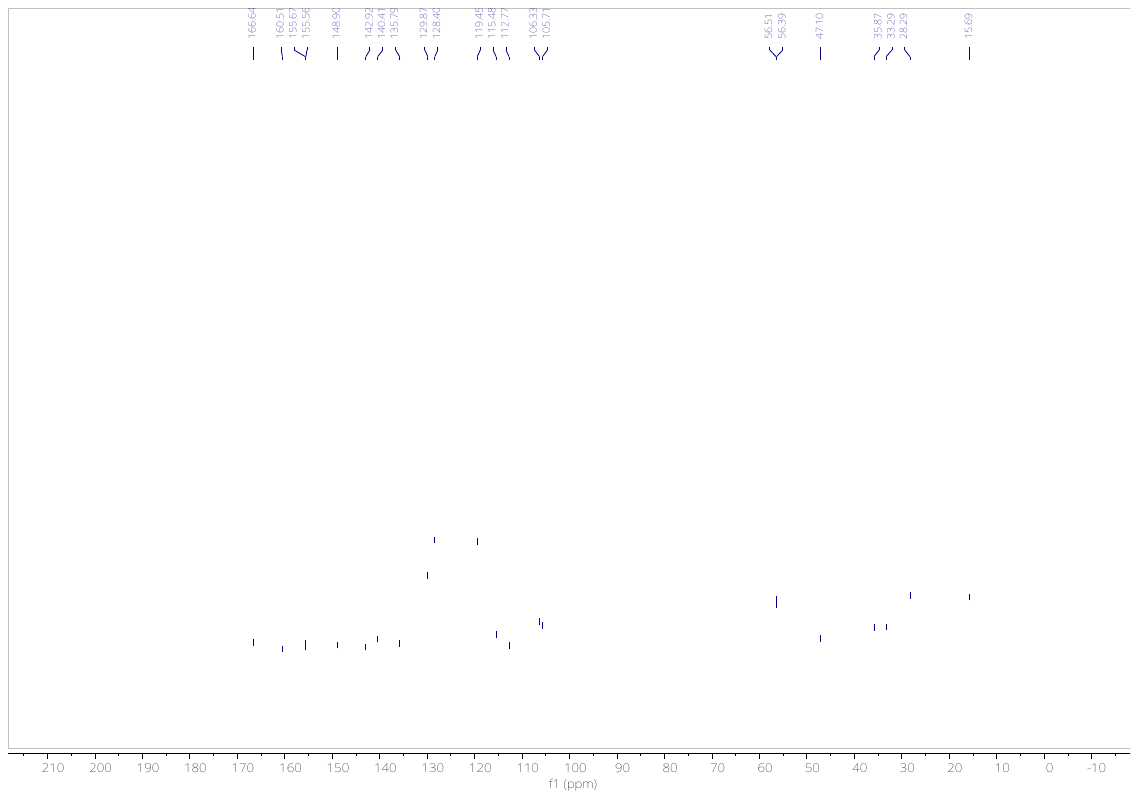

**Figure S42.** ^13^C NMR of **10v** in CDCl_3_.

**2. HRMS of compounds**

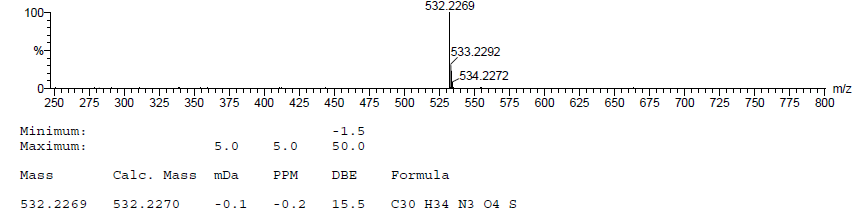
**Fig. S43.** HRMS of **10a**.

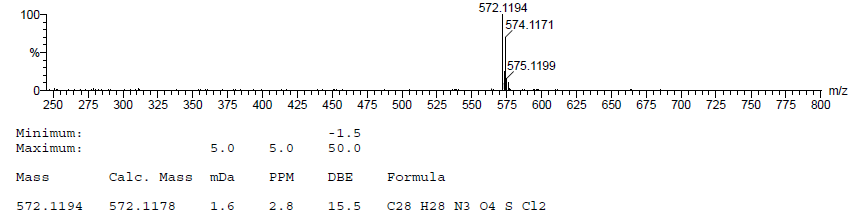
**Fig. S44.** HRMS of **10b**.

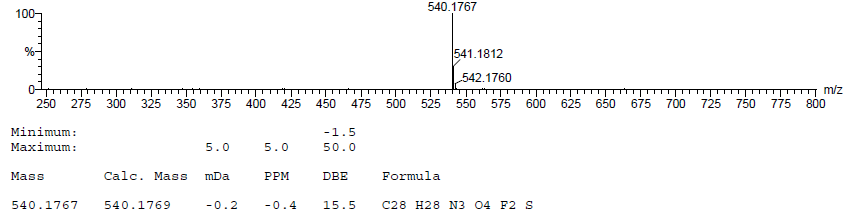

**Fig. S45.** HRMS of **10c**.

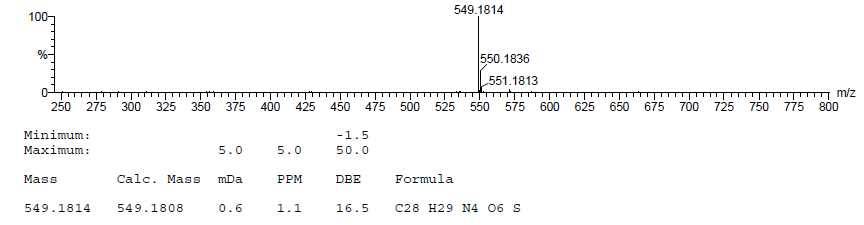
**Fig. S46.** HRMS of **10d**.

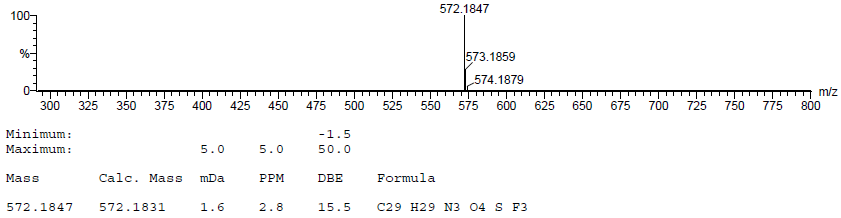

**Fig. S47.** HRMS of **10e**.

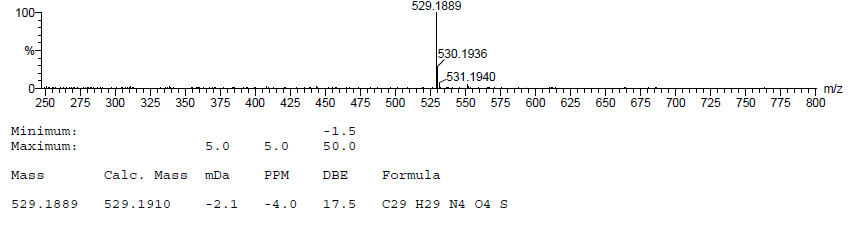
**Fig. S48.** HRMS of **10f**.

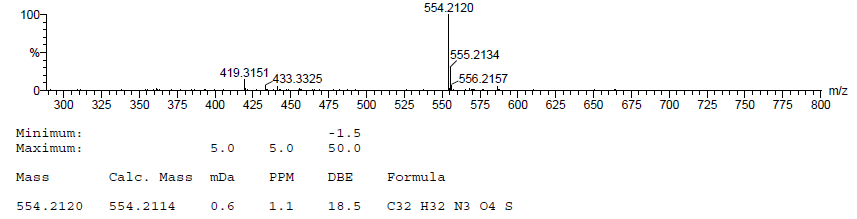
**Fig. S49.** HRMS of **10g**.

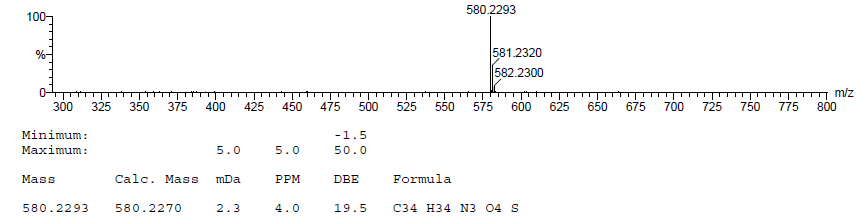
**Fig. S50.** HRMS of **10h**.

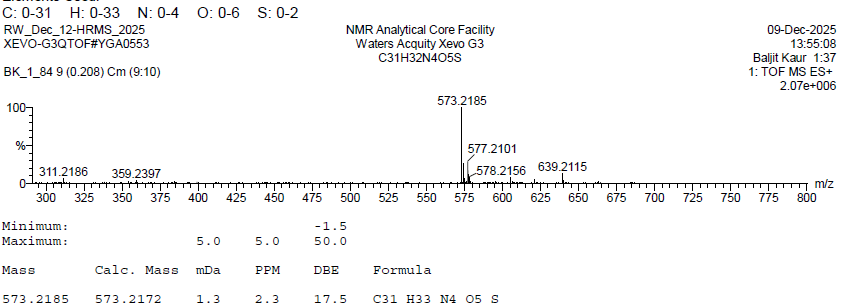

**Fig. S51.** HRMS of **10i**.

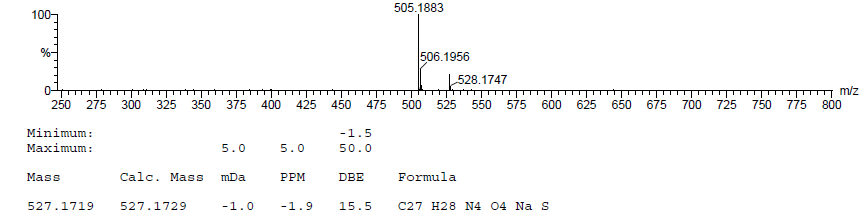
**Fig. S52.** HRMS of **10j.**

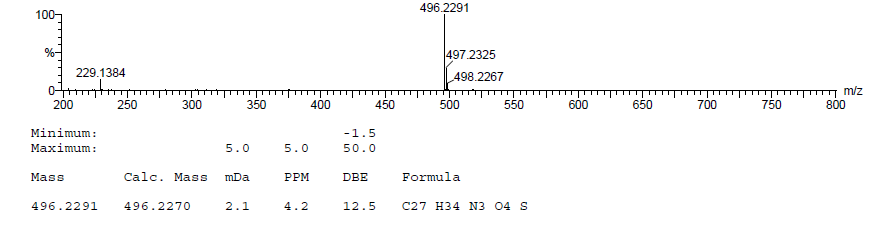
**Fig. S53.** HRMS of **10k**.

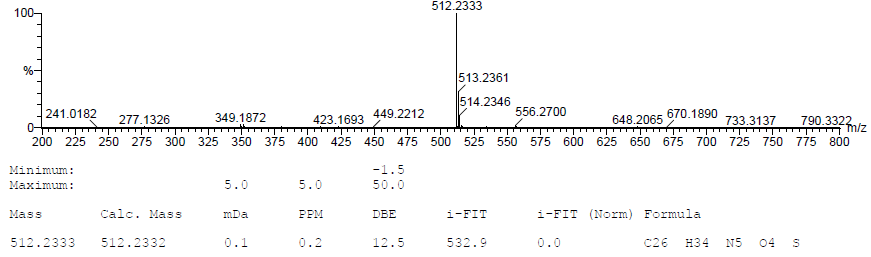

**Fig. S54.** HRMS of **10m**.

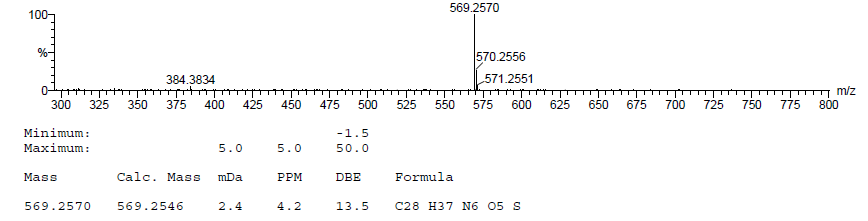
**Fig. S55.** HRMS of **10n**.

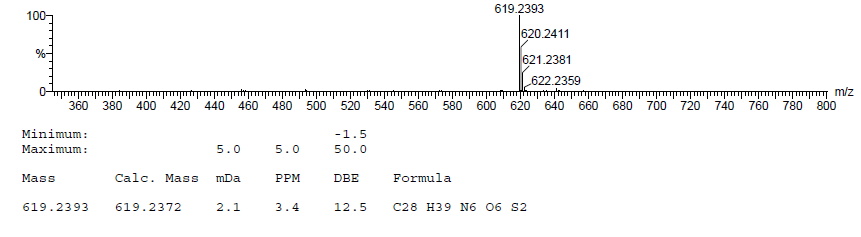
**Fig. S56.** HRMS of **10o**.

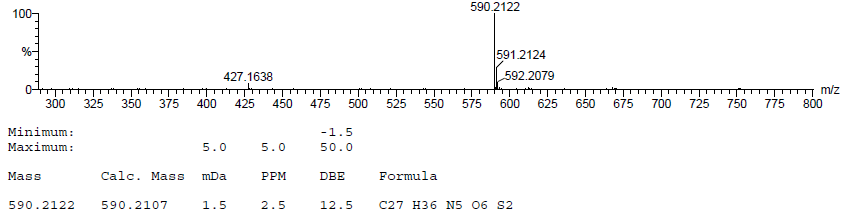
**Fig. S57.** HRMS of **10p**.

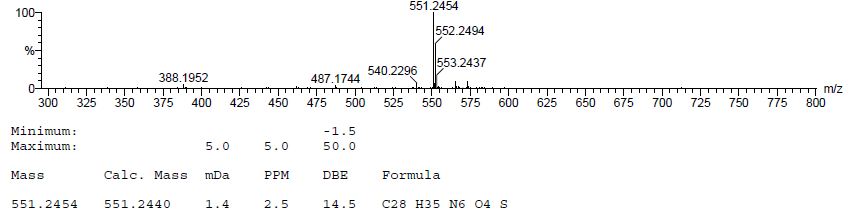
**Fig. S58.** HRMS of **10q**.

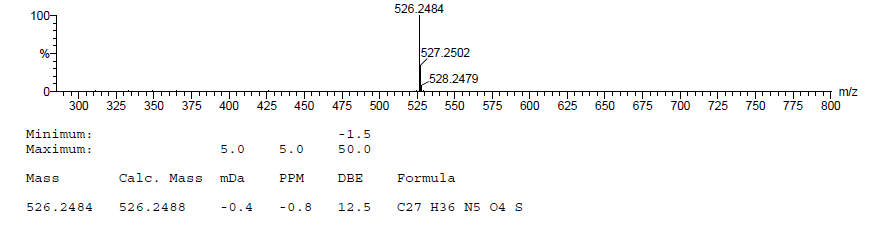
**Fig. S59.** HRMS of **10r**.

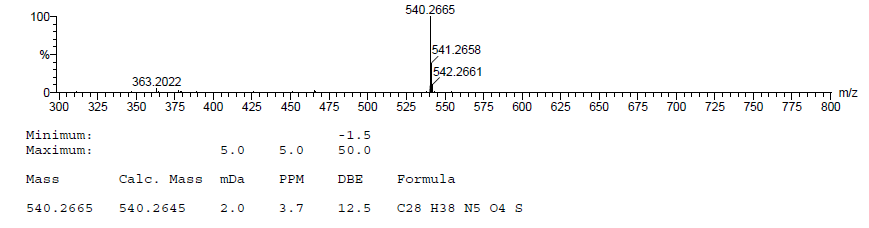
**Fig. S60.** HRMS of **10s**.

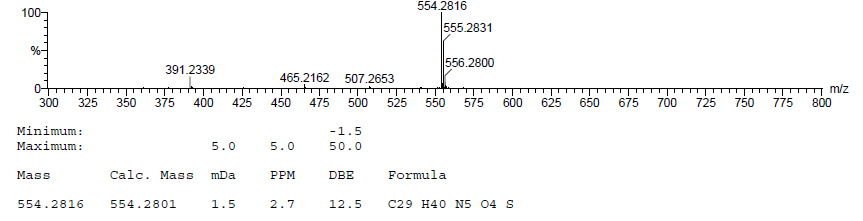
**Fig. S61.** HRMS of **10t**.

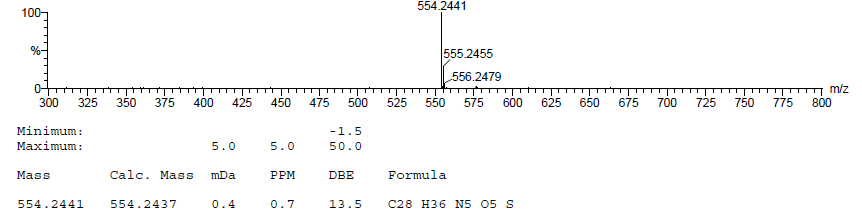
**Fig. S62.** HRMS of **10u**.

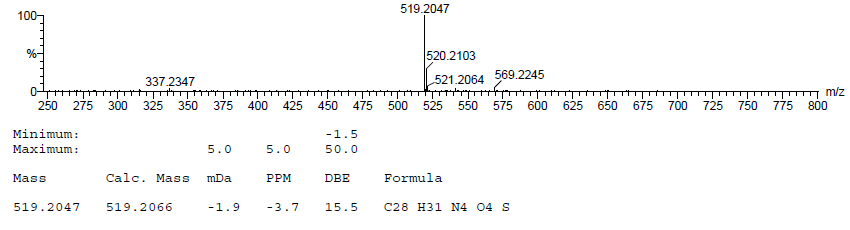
**Fig. S63.** HRMS of **10v**.
